# Diversification or collapse of self-incompatibility haplotypes as a rescue process

**DOI:** 10.1101/2020.03.30.017376

**Authors:** Alexander Harkness, Emma E. Goldberg, Yaniv Brandvain

**Affiliations:** Department of Ecology, Evolution, and Behavior, University of Minnesota; Department of Plant and Microbial Biology, University of Minnesota

## Abstract

In angiosperm self-incompatibility systems, pollen with an allele matching the pollen recipient at the self-incompatibility locus is rejected. Extreme allelic polymorphism is maintained by frequency-dependent selection favoring rare alleles. However, two challenges limit the spread of a new allele (a tightly linked haplotype in this case) under the widespread “collaborative non-self recognition” mechanism. First, there is no obvious selective benefit for pollen compatible with non-existent stylar incompatibilities, which themselves cannot spread if no pollen can fertilize them. However, a pistil-function mutation complementary to a previously neutral pollen mutation may spread if it restores self-incompatibility to a self-compatible intermediate. Second, we show that novel haplotypes can drive elimination of existing ones with fewer siring opportunities. We calculate relative probabilities of increase and collapse in haplotype number given the initial collection of incompatibility haplotypes and the population gene conversion rate. Expansion in haplotype number is possible when population gene conversion rate is large, but large contractions are likely otherwise. A Markov chain model derived from these expansion and collapse probabilities generates a stable haplotype number distribution in the realistic range of 10–40 under plausible parameters. However, smaller populations might lose many haplotypes beyond those lost by chance during bottlenecks.

## Introduction

Self-incompatibility (SI), a common strategy by which plants ensure outcrossing, is a classic example of extreme allelic polymorphism maintained by long-term balancing selection. An SI plant rejects self pollen, which is identified by a specificity phenotype encoded by a highly polymorphic self-incompatibility locus (S-locus). SI is widespread in plants: distinct non-homologous SI systems have been discovered in the Poaceae (Li et al. 1997), Papaveraceae (Foote et al. 1994), Solanaceae (McClure et al. 1989), Brassicaceae (Stein et al. 1991), and Asteraceae (Hiscock et al. 2003), and some form of SI is thought to be present in nearly 40% of plant species across 100 families (Igić et al. 2008). Rejection occurs when the pollen specificity matches the pistil specificity, which is likewise encoded by the S-locus. Pollen with a rare specificity has an advantage because it is less likely to encounter a pistil with a matching specificity and is thus less likely to be rejected. This advantage of rarity results in balancing selection, which maintains polymorphism at the S-locus by protecting S-locus alleles (S-alleles) from loss through drift (Wright 1939). While it is “fairly obvious that selection would tend to increase the frequency of any additional alleles that may appear” (Wright 1939), it is not obvious how any additional alleles may appear. The origin of novel S-alleles is most mysterious in the “collaborative non-self recognition” SI system, whose molecular mechanism has been recently unraveled. Collaborative nonself-recognition, which is described below in the section of the same name, appears inhospitable to novel S-alleles for two reasons. First, because a style with a novel specificity suffers from pollen limitation, and because pollen capable of fertilizing non-existent stylar specificities experience no siring advantage, the spread of a novel allele presents a “chicken-or-egg” problem. Second, we show that if a new S-allele can invade, pollen types incapable of fertilizing it risk extinction because of their siring disadvantage, and therefore a new S-allele can lead to a collapse in the number of S-alleles, rather than an expansion. We develop a population genetic model of the expansion, collapse, and longterm evolution of S-allele number under this widespread SI system, and show when and how these challenges can be overcome, providing many testable hypotheses for the diversification of S-alleles in this system.

While the SI system originally described in *Nicotiana* (East and Mangelsdorf 1925) is particularly well-studied, a molecular genetic understanding of the “collaborative non-self recognition” in this system has only been elucidated recently. Counts of 10–28 alleles have been directly observed in several other species in Solanaceae, and *Physalis crassifolia* has been estimated to harbor as many as 44 (Lawrence 2000). In this SI system, each haploid pollen grain carries one allele at the S-locus, and the pollen is rejected if its allele matches either of those carried by the diploid pollen recipient (East and Mangelsdorf 1925). It is classified as a form of gametophytic SI (as opposed to sporophytic) because the pollen’s phenotype is determined by the genotype of the haploid male gametophyte (i.e., the pollen’s own genotype) as opposed to the genotype of the diploid sporophyte plant that produced it. The SI systems in the Plantaginaceae and the distantly related Rosaceae have been found to be homologous to that in the Solanaceae, which implies that this system was present in the common ancestor of the asterids and rosids (Igić and Kohn 2001; Steinbachs and Holsinger 2002), estimated to have lived about 120 mya (Tank et al. 2015). We refer to it as the collaborative nonself-recognition system after its hypothesized mechanism (Kubo et al. 2010) to distinguish it from other, non-homologous gametophytic SI systems but note that this mechanism has only been definitively demonstrated in *Petunia*, and recognize that even homologous systems may make use of distinct molecular mechanisms. Nonetheless, S-alleles appear to be very long-lived, consistent with long-term balancing selection. A given allele is often more closely related to an allele in another species or even another genus than it is to other alleles in the same population, which is consistent with the alleles having persisted since the common ancestor of those species or genera (Igić and Kohn 2001). Understanding the forces shaping allelic diversity within and across species requires theory that incorporates our modern molecular genetic understanding of the structure and function of the S-locus.

### Conceptual challenges

A series of influential models investigated how a system of many incompatibility alleles could arise, but only very recently have theorists incorporated assumptions reflecting the contemporary understanding of the molecular genetics underlying SI. Under the simplest self-recognition system of SI, Wright (1939) showed that each fully functional S-allele is under negative frequency-dependent selection because pollen carrying rare alleles is compatible with a larger fraction of the population. As such, the number of S-alleles is limited only by their rate of input by mutation, and their stochastic loss from a population by drift. Charlesworth and Charlesworth (1979) showed that in SI system in which a single mutation can generate a new specificity, novel functional S-alleles can invade when inbreeding depression is high. However, single-mutation models of incompatibility are inconsistent with the separate pistil- (McClure et al. 1989) and pollen-expressed (Lai et al. 2002; Entani et al. 2003; Sijacic et al. 2004) products that were later discovered. Uyenoyama et al. (2001) and Gervais et al. (2011) showed that if a new specificity requires separate mutations in a pollen-function and a tightly linked pistil-function locus, new S-alleles can be generated through an self-compatible (SC) intermediate, but one or more S-alleles are also lost for many parameter combinations. Sakai (2016) showed that if each pollen-function mutation is more likely to be rejected by the pistil-function allele that was on its initial genetic background, then the number of S-alleles can increase from zero into the range observed in nature.

A critical common element among all these models is the assumption that SI functions through self-recognition. In a self-recognition system, such as the SCR/SRK system found in the *Brassicaceae* (Kachroo et al. 2002), pollen is accepted by default, but self pollen carries a product that induces rejection in the maternal plant. However, the SI system in question functions not through self-recognition, but rather through a “collaborative nonself-recognition mechanism” (Kubo et al. 2010). In a nonself-recognition system, pollen is rejected by default, but cross pollen carries a product that allows it to bypass rejection. The crucial difference from a theoretical perspective is in the behavior of a mutant that expresses a new, unrecognized specificity. A new specificity would be compatible with existing phenotypes under self-recognition (it is new, but still not self), whereas it would be incompatible with existing phenotypes under nonself-recognition (it is no longer recognized as nonself). Results from models of self-recognition systems might therefore not be directly applicable to nonself-recognition. Despite the ubiquity of collaborative non-self recognition-based SI, few models of S-allele evolution explicitly consider this system. Fujii et al. (2016) proposed that a pollen-function mutation complementary to a novel pistil-function mutation could arise on a single background and then spread to other backgrounds through gene conversion. Bod’ová et al. (2018) enumerated several possible evolutionary pathways to new S-alleles and demonstrated that the pathway hypothesized by Fujii et al. (2016) could either expand or reduce S-allele number. An explanation of the long-term trajectory of S-allele number must therefore account not only for the steps needed to create new fully functional alleles but also for dynamics that can lead to their loss.

The stochastic process involving the risk of collapse, which follows from the collaborative non-self recognition system, raises radically different questions than one where drift is the only force opposing continual growth. Are S-allele numbers repeatedly expanding and contracting, or do they reach equilibria where changes are rare? We might sometimes observe changes in progress if they are frequent, but infrequent changes might be buried in the past. When a contraction occurs, is it usually small or catastrophic? Large contractions, followed by re-expansion, should result in more turnover in alleles and fewer shared alleles among species than would small contractions. Since contraction is possible whether SI is maintained throughout (Bod’ová et al. 2018) or is temporarily lost in SC intermediates (Uyenoyama et al. 2001; Gervais et al. 2011; Bod’ová et al. 2018), neither pathway is unidirectional. Is the pathway maintaining SI still sufficient for expansion, or are SC intermediates necessary? If new proto-S-alleles are segregating in natural populations, are they self-compatible, self-incompatible, or both? Answers to these questions will aid interpretation of S-allele genealogies, which document both similarities and differences between species in their complement of S-alleles. For example, the large historical contraction in S-allele number in the common ancestor of *Physalis* and *Witheringia* has been interpreted as a demographic bottleneck (Paape et al. 2008), but a firmer theoretical investigation could assess whether a collapse restricted to the S-locus is a plausible alternative explanation.

We find that diversification and collapse of S-alleles result from a rescue process similar to evolutionary or genetic rescue of a population. To approximate the relative probabilities of collapse and expansion in S-allele number as well as the distribution of collapse magnitudes, we therefore expand the analogy of evolutionary rescue (Orr and Unckless 2008) to model the extinction of haplotypes and to incorporate the frequency-dependent dynamics associated with SI. We find a negative relationship between initial S-allele number and the probability of further expansion. By constructing and iterating a Markov chain out of these contraction and expansion probabilities, we find stable long-term distributions of S-allele number. We show that for large but plausible values of population size and rate of gene conversion, this process can generate numbers of S-allele numbers comparable to those found in nature (20–40). However, contractions can be very large when they occur, often eliminating the majority of S-alleles. These results suggest that an evolutionary opportunity to expand the number of S-alleles is intrinsically connected to the risk of a contraction in S-allele number.

Our results yield numerous novel predictions. First we predict that novel stylar rejection alleles will arise on SC haplotypes which are maintained by mutation-selection balance. Second, we predict that, when many but not all S-alleles are fortuitously capable of fertilizing plants carrying the novel S-allele, the S-alleles lacking this capability are very likely to be lost. This is because the alleles possessing this fortuitous compatibility eliminate their competitors when a novel specificity arises, thereby reducing the total number of surviving alleles. However, the fortuitously compatible S-alleles themselves are never expected to be lost at equilibrium. The joint effect of increasing the number of fortuitously compatible S-alleles is to increase the probability that some S-alleles are lost but also to increase the minimum number of surviving S-alleles. Third, we predict that intraspecific variation in S-allele number can sometimes be explained by an invasion of a runaway SI allele rather than a bottleneck or a transition to SC. Counterintuitively, we therefore predict that when allopatric populations exhibit large disparities in S-allele number, the population with the smaller number of S-alleles will harbor a stylar allele absent from and incompatible with pollen from the population with more S-alleles, while all S-alleles from the population with more S-alleles can be fertilized by pollen in the population with fewer S-alleles.

## Model and Results

### Collaborative Nonself-Recognition

The risk of contraction in S-allele number in the “collaborative nonself-recognition” incompatibility system is rooted in the system’s biological details. This system involves two kinds of complementary products: pistil-expressed ribonucleases (RNases) (McClure et al. 1989) and pollen-expressed F-box proteins (Lai et al. 2002; Entani et al. 2003; Sijacic et al. 2004). Together, these products achieve a form of collaborative nonself-recognition first described by Kubo et al. (2010). Under this system, each “allele” at the S-locus is actually a tightly linked haplotype containing one RNase gene and a collection of paralogous F-box genes. The RNase gene is highly polymorphic, and each functionally distinct haplotype possesses a different RNase allele. Nonself pollen is recognized through the complement of F-box proteins it expresses. Each F-box paralog produces a functionally distinct product, and each of these products is capable of detoxifying one or more forms of RNase. Pollen is only successful if it expresses both of the two F-box proteins to match the diploid pollen recipient’s two RNase alleles. Self-fertilization is prevented because each fully functional haplotype lacks a functional copy of the F-box gene that corresponds to the RNase on the same haplotype, and so each pollen grain necessarily lacks one of the two F-box proteins required to fertilize the plant that produced it. Rejection is not restricted to self-pollination: any two individuals that share one haplotype will reject half of each other’s pollen, while individuals that share both haplotypes are completely incompatible. Rejection is therefore more likely between closely related individuals. Tight genetic linkage across all components of the S-locus reduces the probability that recombination will cause a haplotype to lose a functional F-box paralog (reducing its siring opportunities) or gain the F-box paralog that detoxifies its own RNase (inducing SC).

This system presents two novel challenges that are absent in self-recognition systems. The first is a “chicken-egg problem.” A novel F-box specificity alone is at best neutral because it detoxifies an RNase that does not yet exist. A novel RNase alone is deleterious because it degrades all pollen and renders the plant ovule-sterile. Both of these mutations must invade in order to generate a new, fully functional S-haplotype. Second, for every novel RNase that arises, the corresponding F-box specificity must appear on every other haplotypic background in order to restore cross-compatibility among all haplotypes. Building on the hypothesis for expansion of haplotype number proposed by Fujii et al. (2016), Bod’ová et al. (2018) showed how cross-compatibility could be restored among all incompatibility classes after novel RNase and F-box mutations have invaded. If an initially neutral F-box mutation already exists when its complementary RNase mutation arises, the F-box mutation will then invade because it confers the advantage of compatibility with the new RNase. To restore full cross-compatibility among haplotypes, all haplotypes must acquire F-box paralogs complementary to all RNase alleles other than their own either through gene conversion (Fujii et al. 2016) or recurrent mutation (Bod’ová et al. 2018). This means that the haplotype bearing the RNase mutation must also acquire the F-box complementary to its ancestor. Once this occurs, the resulting haplotype is compatible with pollen recipients carrying all other haplotypes, but haplotypes still lacking the new F-box are not compatible with pollen recipients carrying the new RNase. As the RNase mutation increases in frequency, siring opportunities decrease for haplotypes that still lack the new F-box, and they are gradually driven extinct. If all doomed haplotypes acquire their missing F-box before they are lost, expansion has occurred. But if some doomed haplotypes go extinct, their RNase alleles are lost and contraction has occurred. Bod’ová et al. (2018) simulated the dynamics of this process along with several other expansion pathways and found that up to 14 haplotypes could be maintained.

Based on this biological background, we introduce a set of metaphors to make the interactions among haplotypes more intuitive. Each form of pistil-function RNase is a lock, and each form of pollen-function F-box protein is a key. A diploid plant codominantly expresses two different locks in its pistils. Each pollen grain expresses every paralogous key in its haploid genome, and these keys collectively form that pollen grain’s key ring. The pollen must unlock both of the pollen recipient’s locks in order to fertilize it, which requires keys to both locks. Each fully functional haplotype is SI because its key ring lacks the key to the lock on the same haplotype (fig. 1).

**Figure 1:**
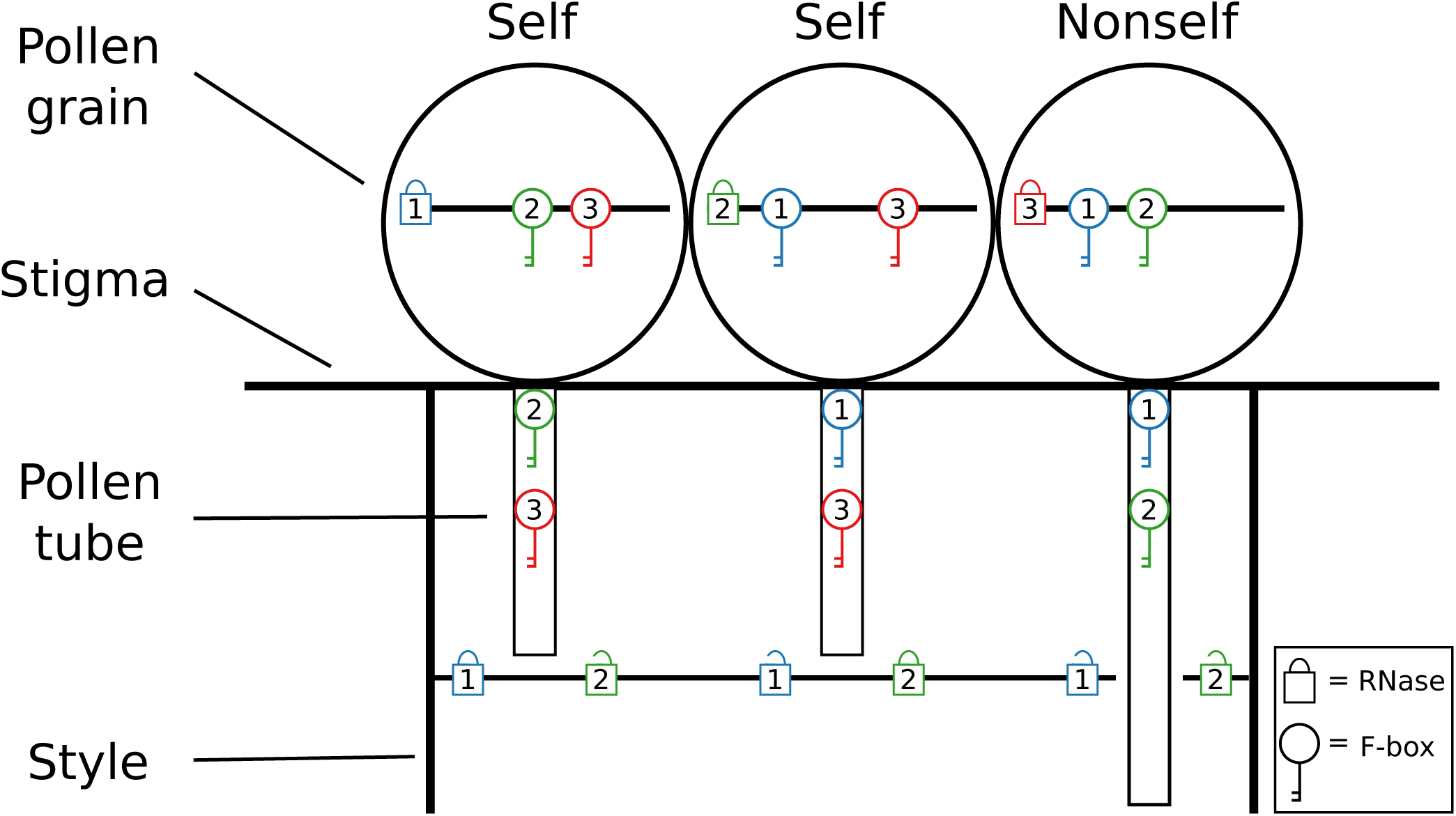
Rejection of self pollen in the style. Each haplotype contains a single lock and multiple paralogous keys. Each pistil expresses the two locks in its diploid genotype, and each pollen grain expresses all keys in its haplotype. A pollen tube is arrested in the style unless it contains keys to both of the pistil’s locks. Each haplotype lacks the key to its own lock, but contains the keys to all other locks. Self-fertilization is prevented because pollen necessarily lacks one of the keys expressed by the plant that produced it.

In contrast to Bod’ová et al. (2018) our model addresses the realistic possibility that novel RNase alleles (locks) will have ovular fitness below the population average because keys to this lock are initially rare. Thus, while Bod’ová et al. (2018) assumed the novel RNase was neutral, we model the case when this RNase suffers more intense pollen limitation than the rest of the population. How could such a stylar mutation, which increases pollen limitation, increase ovular fitness (as is required for its adaptive spread)? We suggest that novel locks can be favored on self-compatible genetic backgrounds otherwise maintained by the balance of selection against self-compatibility and gene conversion of F-box alleles which restore self-compatibility. By restoring SI and preventing self-fertilization, a new lock can provide an ovular advantage and can spread by natural selection. Concurrently, the complementary key spreads within and among haplotypes (“key rings”), and the number of S-allele increases if all S-alleles are “rescued” before S-haplotypes lacking this novel key are driven to extinction due to their mating disadvantage.

### Model outline

We identify six haplotype classes with regard to their phenotypes expressed in pollen and pistils (fig. 2), and we evaluate a three-step evolutionary pathway to expansion or contraction of S-haplotype number (fig. 3). We define “contraction” as any transition to a state with fewer lock alleles. We define “expansion” as the transition from an initial state in which all haplotypes are cross-compatible with all other haplotypes to a final state that is the same but with one additional cross-compatible SI haplotype.

**Figure 2:**
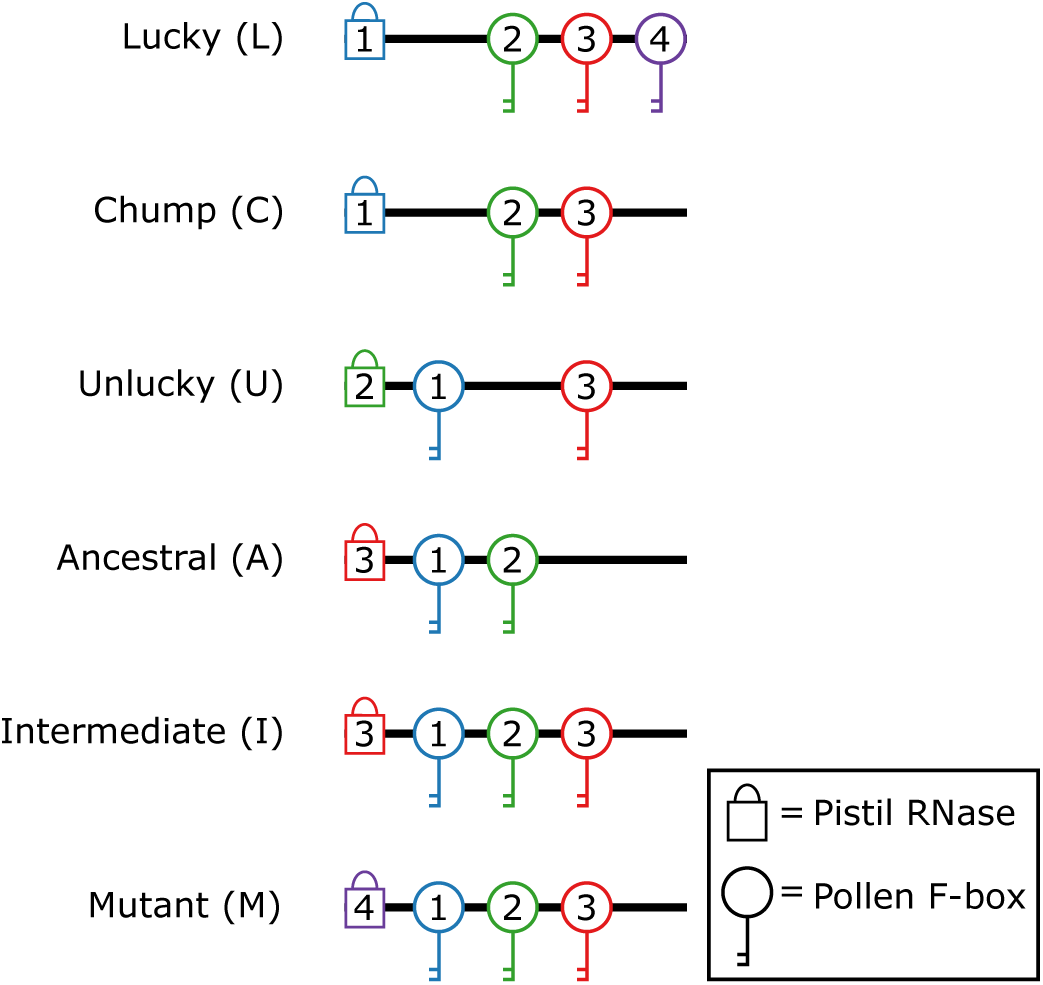
Haplotype classes. In this example, the population begins with three complete haplotypes (those with the key to every lock but their own). Gene conversion generates an SC “intermediate” haplotype from the “ancestral” haplotype by introducing the key to the ancestral haplotype’s own lock. The SC intermediate acquires a novel lock by mutation, restoring SI and generating the “mutant” haplotype. All haplotypes lack the key to the new lock except for those “lucky” enough to carry it in advance, which have an advantage over the “chump” haplotypes that bear their corresponding locks. Haplotypes that are “unlucky” merely lack the new key but don’t compete against other haplotypes with their same locks.

**Figure 3:**
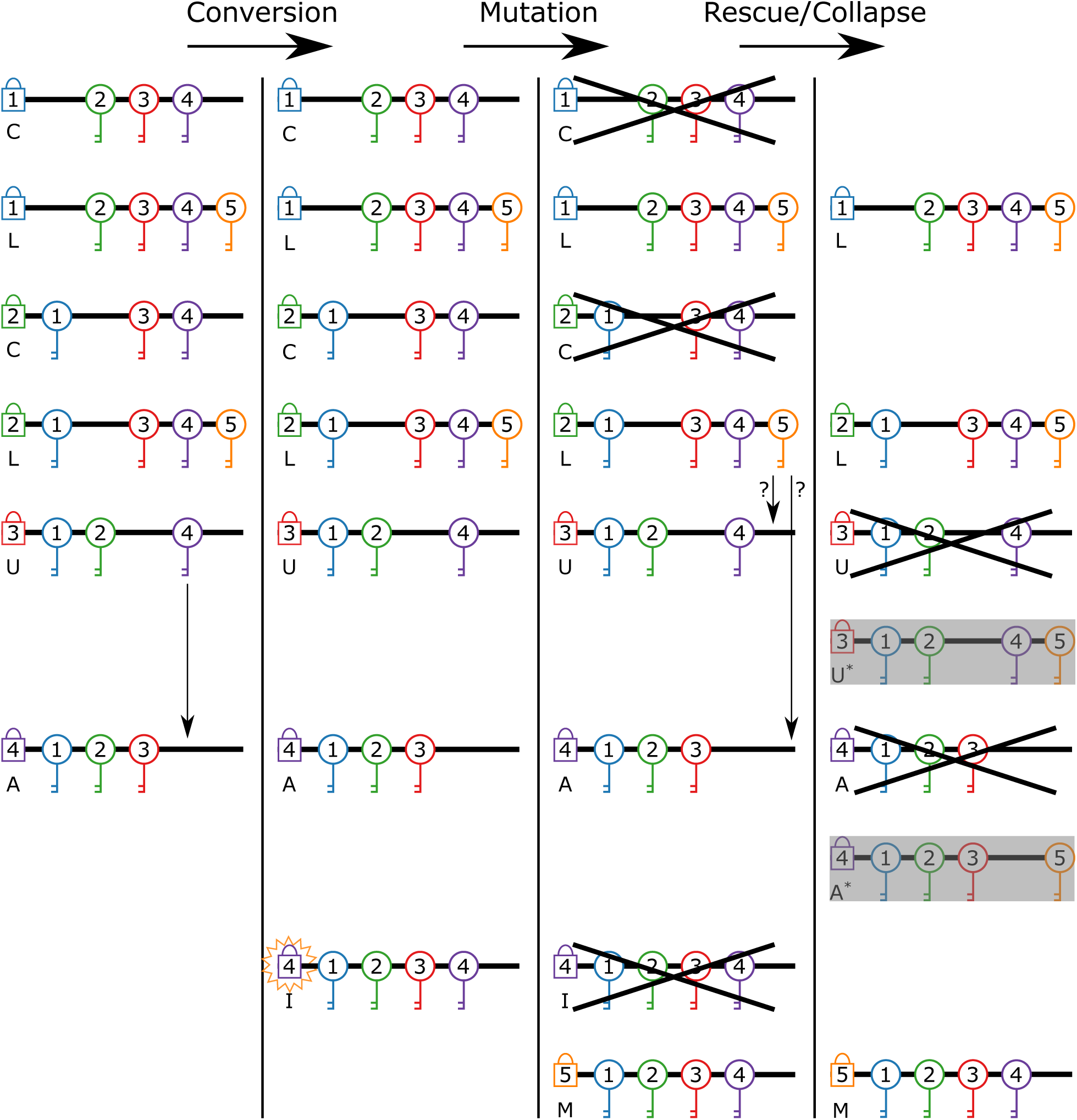
Model steps. First, gene conversion (vertical arrow) generates the SC intermediate I. Second, a mutation in the intermediate’s lock (starburst) generates a mutant SI haplotype M. The intermediate is deleterious and transient, though recurrent mutation continually produces phenotypically identical SC haplotypes. Now that their key has a complementary lock, L haplotypes increase in fitness and outcompete their corresponding C haplotypes. Third, the other haplotypes lacking the new key (U, A) are driven to extinction, but gene conversion might or might not “rescue” them (arrows with question marks) by producing convertant haplotypes carrying the new key (gray boxes).

In Step 1, the “conversion” step, a haplotype acquires the key to its own lock through gene conversion, rendering it SC. This ensures that when a lock mutation occurs on this haplotype, the resulting mutant will already possess every key but its own and will be cross-compatible as a sire with all other haplotypes as soon as it arises. A supply of such SC haplotypes is maintained at the balance between gene conversion and strong selection against inbreeding. The invasion and fixation of an SC haplotype should function identically under self- and nonself-recognition, and this process has already been thoroughly covered by Uyenoyama et al. (2001) and Gervais et al. (2011). We return to it briefly below. However, we focus on cases in which inbreeding depression is strong enough that SC is at a net disadvantage—ours is a model of diversification of S-haplotypes, not the transition to SC. For such cases, there is no risk that the population will become entirely SC: the only possible outcomes are increase, decrease, and stasis in the number of SI haplotypes.

In Step 2, the “mutation” step, the lock allele on the SC intermediate haplotype mutates to a new specificity, one that can be unlocked by an existing, previously neutral key variant. The new lock is a necessary component of a new haplotype. The SC intermediate previously had the key to every lock, but now that its own lock has mutated, the resulting mutant has the key to every lock but its own and is once again SI. The pre-existence of a complementary key on other haplotypes ensures there is at least some pollen compatible with the new lock. However, in the likely event that the complementary key is present on only a subset of other haplotypes, the new lock will reject more pollen than other locks. If the amount of compatible pollen received is a limiting factor on seed set, the lock mutation might reduce ovule success. But since the mutant is compatible as a sire with all other haplotypes (thanks to the “conversion” step), its increased pollen success can compensate for lost ovule success. This might seem to be asking a lot of nature in this step – the mutation to a new lock coincidentally occurs on a rare SC haplotype, and the lock mutation’s matching key already exists in the population despite being previously neutral. However, these events appear less contrived when we consider the alternatives and realize that there is always an opportunity for these rare events to occur. If the lock mutation occurs on an SI haplotype, the resulting mutant will derive no advantage because it was already SI, but it will still suffer additional pollen limitation. If the lock mutation lacks a complementary key, it will reject all pollen and suffer complete ovule failure. Such mutations should surely occur, but they should remain at low frequencies because they are strongly deleterious. Since there is a constant supply of SC haplotypes at gene conversion-selection balance, the population can wait indefinitely until one such haplotype mutates to a lock with a pre-existing key.

Finally, in Step 3, the “rescue/collapse” step, the remaining haplotypes lacking the key to the new lock are driven to extinction unless they can acquire the missing key through gene conversion in their remaining time. This restores mutual cross-compatibility among all surviving haplotypes, whether there are more (expansion) or fewer (contraction) than the initial number.

The six relevant haplotype classes are described in figure 2. Table 1 gives notation for all parameters and variables. There are initially *n* lock alleles, but diversity arises from which keys each haplotype possesses. The first haplotype class consists of *n*_L_ haplotypes that possess one of the initial *n* locks and a key ring that is “complete” (Bod’ová et al. 2018). That is, they possess the keys to all extant locks but their own, including the mutant lock. This first class is denoted *L* for “lucky” because its members fortuitously possess the key to the new lock. Each haplotype in this class initially occurs at frequency *d*_*L*_/*n*, where the parameter *d*_*L*_ is an arbitrary initial proportion. In nature, the parameter *n*_*L*_ would be determined by the extent of gene conversion that has already occurred: if the neutral novel key has been copied from its original background to many others, then *n*_*L*_ is high. Second, for each of these *n*_*L*_ complete haplotypes, there exists one haplotype that is identical except that it lacks the key to the new lock. These could represent ancestral versions of the complete haplotypes that have not yet gained the key to the new lock. This second class is denoted *C* for “chump” because members of this class are simply inferior versions of the corresponding “lucky” haplotypes. Each haplotype in this class occurs at frequency (1 − *d*_*L*_)/*n*. Third, there are *n*_*U*_ haplotypes that have one of the initial *n* locks and an incomplete key ring that lacks only the key to the mutant lock. Unlike the previous class, there exists no complete version of these haplotypes initially. This third class is denoted *U* for “unlucky” because members of this class cannot unlock the new lock like the “lucky” haplotypes can, but neither must they compete with a superior version of themselves like the “chump” haplotypes must. Each haplotype in this class occurs at frequency 1/*n*. Fourth, there is the ancestral haplotype from which the SC intermediate arose. It is like any other incomplete haplotype (e.g., a *U* haplotype), except that it has the same lock as the SC intermediate. This fourth class is denoted *A* for “ancestral” because its lock is ancestral to that of the lock mutant. It occurs at frequency 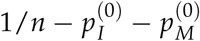, where 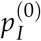 is the initial frequency of the SC intermediate, and 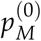 is the initial frequency of the lock mutant. Fifth, there is the SC intermediate. Pollen carrying this haplotype is compatible with all plants, and pistils carrying it reject only the “A” haplotype. This fifth class is denoted *I* for “intermediate” because it is the intermediate between the ancestral class and the new SI haplotype to be generated. Note that gene conversion should generate some SC haplotypes from all classes, not just Class A. We do not name these haplotypes because all SC haplotypes should remain at low frequency, and we only name Class I because it is the background on which the lock mutation arises. We lump all other SC haplotypes into the class from which they originated (*L* or *U*). Sixth, there is the haplotype bearing the lock mutation. It is identical to the intermediate haplotype, except that it has a novel lock and is thus SI. It is denoted *M* for “mutant” because of its novel lock mutation. This haplotype increases its own fitness over its SC predecessor by preventing selfing, but it also decreases the fitness of the “unlucky” and “chump” haplotypes by presenting a lock to which they lack the key. It occurs at initial frequency 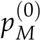 There are thus *n*_*L*_ + *n*_*U*_ + 1 = *n* pre-existing haplotypes (including the mutant’s ancestor) along with the mutant itself for a total of *n* + 1 haplotypes.

**Table 1:** Notation. Fixed parameters are set at the beginning of the diversification/collapse process and remain constant throughout. Initial conditions are set at the beginning of the process, but their only effect is to determine the initial genotype and haplotype frequencies. Dynamic variables vary throughout the simulation and continue to be tracked. The dynamic variables are not independent: the collection of genotype frequencies is sufficient to describe the current state of the population completely.

|  |  |  |
| --- | --- | --- |
| $n$ | Fixed parameter | Number of locks, excluding the mutant |
| $n_L$ | Fixed parameter | Number of lucky haplotypes |
| $n_U$ | Fixed parameter | Number of unlucky haplotypes |
| $G$ | Fixed parameter | Number of pollen grains received on each stigma |
| $\mu$ | Fixed parameter | Per-individual rate at which gene conversion produces SC haplotypes |
| $R_{conversion}$ | Fixed parameter | Total rate at which gene conversion produces rescuing haplotypes |
| $d_L$ | Initial condition | Initial frequency of new key within a lucky-unlucky pair |
| $p_X$ | Dynamic variable | Frequency of all haplotypes of class $X$ |
| $P_{XY}$ | Dynamic variable | Frequency of genotype with haplotypes of classes $X$ and $Y$ |
| $F_{XY}$ | Dynamic variable | Frequency of pollen compatible with genotype $XY$ |
| $D_{XY}$ | Dynamic variable | $P_{XY}/F_{XY}$ |
| $w_X$ | Dynamic variable | Fitness of any haplotype of class $X$ |
| $\bar{w}$ | Dynamic variable | Mean fitness of all haplotypes |
| $w_R$ | Dynamic variable | Fitness of a potentially rescuing gene convertant |
| $s$ | Dynamic variable | Additional fitness of a gene convertant, $w_R - \bar{w}$ |

Throughout out model, we assume that selection dominates random genetic drift, so we use deterministic equations to describe the change in haplotype frequency by selection. As such, the frequency of the SC intermediate after Step 1 (fig. 3), “conversion,” is controlled by the deterministic equilibrium between selection and gene conversion, and the decay of doomed haplotypes after Step 2, “mutation,” occurs through deterministic genotype frequency trajectories (see Appendix). However, our model includes numerous stochastic forces that play a critical role in mediating the diversification or collapse of S-allele diversity. The first event we model as a random variable is the distribution of compatible pollinations on the lock mutant after Step 2. This distribution is used to demarcate parameter values in which compatible pollen supply does or does not limit ovule success. Most critically, we model the probability of survival of new gene convertants in Step 3, the “rescue/collapse” step, as a stochastic process. Specifically, we use the branching processes approximation of Fisher (1923) and Haldane (1927) to approximate this survival probability based on the fitness benefit of the convertant, though we check the accuracy of this approximation through stochastic simulation. The third and final random variable is the number of surviving haplotypes after rescue/collapse, which is itself calculated from the survival probability and the expected supply of gene convertants. We use this distribution to parameterize a transition matrix, and iterate this as a Markov Chain to determine the long-run stable distribution of haplotype number numerically. Though we do not model long-term drift in allele frequencies (only the short-term stochastic loss of new alleles), we nevertheless refer to a population size parameter in order to convert the proportion of gene convertants into a whole number of copies (the relevant quantity for rescue probability). If the long-term number of haplotypes predicted by Bod’ová et al. (2018) is limited by the supply of gene convertants in the small populations that can feasibly be simulated, a predominantly deterministic approach may allow us to examine populations with greater supplies of gene convertants and possibly greater long-term haplotype numbers.

### Equilibrium frequency of SC intermediate

Step 1, the “conversion” step, consists of the generation and equilibration of the SC intermediate (fig. 3). The SC intermediate provides the advantage of compatibility with other individuals carrying the same lock: this advantage can counteract pollen limitation when the SC intermediate later acquires a new lock by mutation. However, SC also carries the disadvantage of allowing some self-fertilization. We are not interested in the situation in which SC haplotypes invade and fix in the population. We therefore focus on cases in which SI is maintained by extreme inbreeding depression: specifically, all selfed offspring are eliminated before they can reproduce. In this case, for many values of S-haplotype number and primary selfing rates, the SC intermediate is maintained at low frequency by the balance between gene conversion and selection. What is this frequency?

We determine conversion-selection balance of the SC intermediate using a simple model. A population contains *n* SI haplotypes that are all mutually cross-compatible. Since all haplotypes are cross-compatible, a diploid’s maternal and paternal haplotype carry the key to the other’s lock. This state does not normally result in SC pollen because, though the diploid possesses keys to both of its own locks, these keys are on separate haplotypes. The absence of recombination at the S-locus then ensures that none of the plant’s pollen carries both keys. However, gene conversion allows a lock to be copied from one haplotype to another, potentially generating an SC haplotype from any diploid carrying at least one SI haplotype. In this population, gene conversion in zygotes generates SC haplotypes at rate *µ*. Individuals carrying two SC haplotypes self at rate *σ*, and those carrying one self at rate *σ*/2. We model extreme inbreeding depression such that all selfed offspring are inviable, and so the selfing rate is equivalent to a fecundity penalty. However, SC haplotypes gain an advantage in pollen. Assuming all individuals are heterozygous at the lock locus, SI haplotypes are rejected with probability 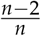, whereas SC haplotypes are never rejected because they carry all keys. If SC haplotypes are rare, the expected change in frequency of all SC haplotypes, *p*, is

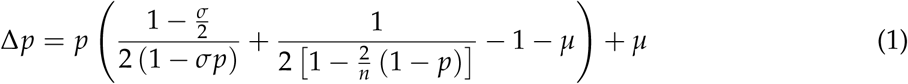

If *n, µ*, and *σ* are treated as constant parameters, Δ*p* is a cubic function of *p*. We determine the equilibrium frequency by setting Δ*p* = 0 and solving numerically using Mathematica’s NSolve function. We solve for the equilibrium frequency of SC haplotypes, varying *µ* from 10^−4^ to 10^−3^ in steps of 10^−4^, *σ* from 0.1 to 1 in steps of 0.1, and *n* from 3 to 40 in steps of 1 (fig. 4). The biologically meaningful domain of Δ*p* is the real numbers between zero and one, so we ignore solutions outside this interval.

**Figure 4:**
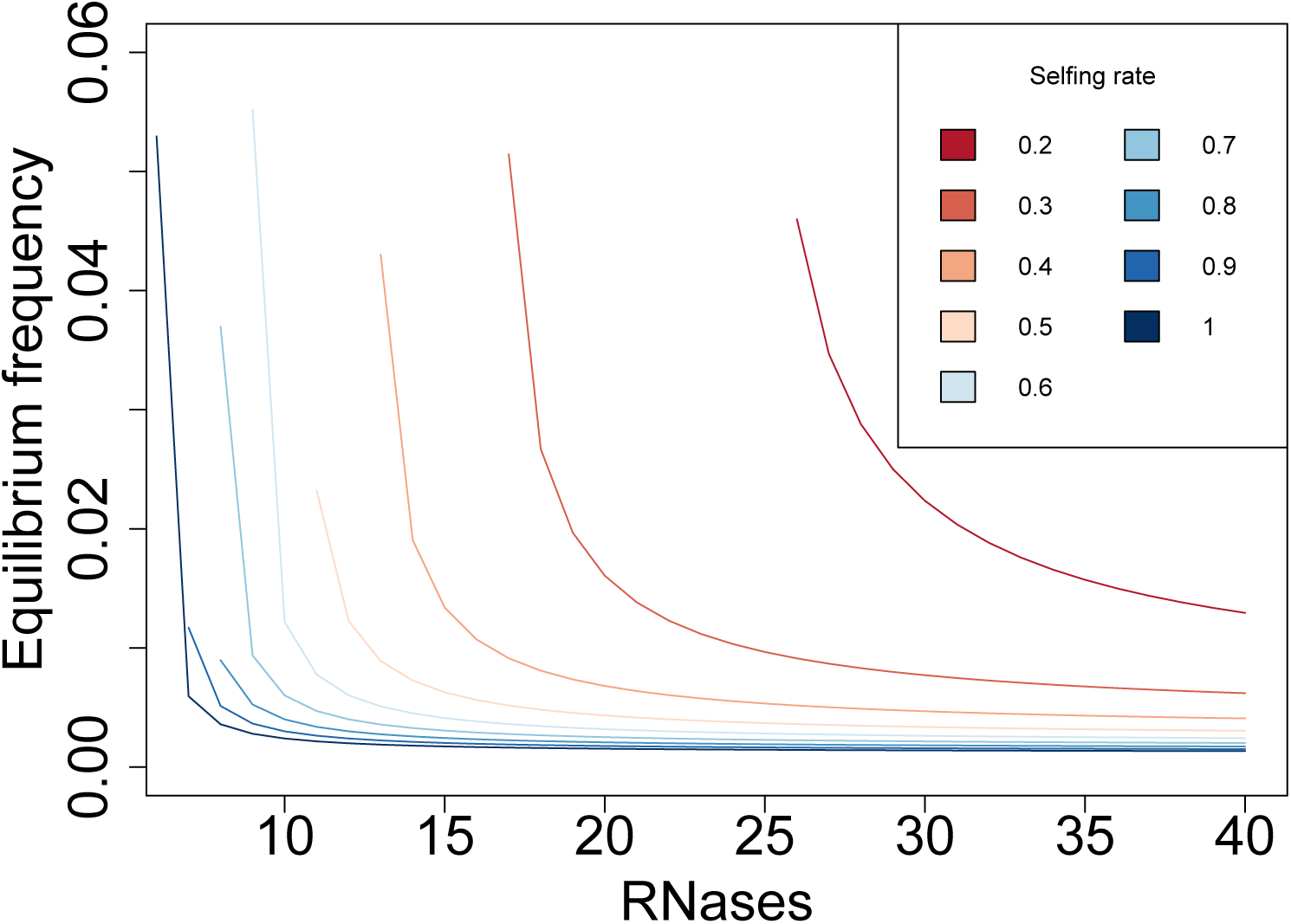
Conversion-selection balance of SC gene convertants. Equilibrium frequency was calculated assuming the gene conversion rate *µ* = 0.0003. Equilibrium frequency declined with increasing RNase number and selfing rate. The left-most point of each curve represents the threshold RNase number below which an SC haplotype will rise to fixation rather than equilibrate at low frequency. This threshold appears as an abrupt stop rather than an asymptote because RNase number can only take on integer values. No internal equilibrium was found for selfing rate *σ* = 0.1.

We find two broad categories of outcomes depending on the parameter values. First are cases in which Δ*p* is positive from *p* = 0 to *p* = 1, and there are no internal equilibria. In such cases, SC haplotypes are expected to rise to fixation despite intense inbreeding depression. This occurs when SC confers a large pollen advantage because *n* is small, or when it incurs a small ovule disadvantage because *σ* is small (Charlesworth and Charlesworth 1979; Uyenoyama et al. 2001). Second are cases in which there are two solutions between *p* = 0 and *p* = 1. In these cases, the smaller solution is stable, and the larger is unstable. The unstable solution tends to be fairly large, usually between 0.1 and 0.5, so we warn that predicted frequencies of unstable equilibria are likely imprecise. Because this internally unstable equilibrium cannot be approached by any biological process, we report and investigate only the stable solution. Stable equilibria for *µ* = 0.0003, with *σ* from 0.1–1.0, and *n* from 0–40 are plotted in (fig. 4).

### Invasion of lock mutant

With the existence of SC haplotypes at conversion-selection balance, we can generate a new SI haplotype through mutation. If an SC haplotype’s lock mutates to a novel specificity, it will no longer have the key to its own lock, making it SI. It will, however, possess the key to every other lock, making it universally cross-compatible as a sire. The SC haplotype therefore acts as an SC “intermediate” haplotype *I* to the new SI “mutant” haplotype *M* (fig. 2). As Bod’ová et al. (2018) showed, competition between complete haplotypes possessing all others’ keys and incomplete haplotypes lacking some keys can result in extinction of the incomplete haplotypes. This occurs because the incomplete haplotypes have pollen success below the population average. In our model, two haplotypes possess the keys to all other locks: the intermediate *I* and mutant *M*, haplotypes. These haplotypes are compatible as sires with every other haplotype, but only a subset of haplotypes are compatible with them. In *I*, the siring advantage partially compensates for the loss of ovule success through selfing, resulting in a smaller but still deleterious net effect on fitness. The mutation from *I* to *M* restores SI while maintaining the siring advantage. However, this mutation also potentially increases pollen limitation because the new lock rejects the majority of pollen in the population. In order for *M* to invade and for the population to progress to the rescue/collapse step, *M* must have fitness above the population average despite the tradeoff of pollen limitation.

We use a simple model of pollen limitation to determine if *M* will invade when rare. As SC haplotypes are rare and of fitness below the population average, so the invasion of *M* depends only on whether *M* is superior to *L* and *U* haplotypes. We assume *L, U*, and *M* haplotypes are equally efficient at rejecting self pollen, and so the quantity of self pollen is irrelevant in determining these haplotypes’ fitnesses relative to each other, and since self pollen acts simply to ‘waste’ ovules, we do not consider self-pollen as a source that alleviates pollen limitation. For simplicity, we assume that each flower carries 10 ovules, and receives a number of pollen grains equal to the parameter *G*. The proportion of *G* that is compatible with the maternal flower is a binomial random variable, but the expectation of this variable is sufficient to determine fitness, so we treat ovule success as a deterministic function of the proportion of the compatible pollen. The proportion of pollen that is compatible with the maternal flower is *n*_*L*_*d*_*L*_/ (*n*_*L*_ + *n*_*U*_) for *UM* genotypes, *d*_*L*_ (*n*_*L*_ − 1) / (*n*_*L*_ + *n*_*U*_) for *LM* genotypes, and (*n*_*L*_ + *n*_*U*_ − 2) / (*n*_*L*_ + *n*_*U*_) for all other genotypes. Fertilization occurs through the following process: one ovule subtracts one compatible pollen grain from the flower’s stock of compatible pollen to produce a seed, then the next ovule does the same, and this process continues until either ovules run out (full seed set) or compatible pollen runs out (partial seed set). We therefore define a flower’s ovule success as min(ovules, compatible pollen)/(ovules). The ovule success of *M* is always less than or equal to the ovule success of *L* and *U* because a smaller proportion of pollen is compatible with it.

The net fitness of *M* also depends on its pollen success. When *M* is rare, *M* pollen is almost never rejected because it is compatible with all non-*M* plants. We therefore define the pollen success of *M* as 1, the maximum value. If *M* is rare, all *L* and *U* haplotypes have a pollen success of approximately (*n* − 2) /*n*, the probability that the pollen matches neither of a random maternal plant’s haplotypes. We then divide the ovule and pollen success of *M* by the mean values of pollen and ovule success (approximated by the success of *L* or *U*, which are equivalent when *M* is rare) to get a measure of its fitness relative to the mean. The *M* haplotype invades if half the sum of these quantities, a measure of relative fitness, is greater than one. We calculate this relative fitness measure for *nL* = 2, *G* = 100–1000, *dL* = 0.01, 0.1, or 0.5, and *nU* = 3, 5, or 7 (fig. 5). We find that the relative fitness of *M* increases as *G* or *dL* increases, and it decreases as *nU* increases. Depending on the choice of *dL* and *nU, M* might have fitness below the population average throughout *G* = 100–1000, fitness above the population average throughout, or there might be a threshold value of *G* within 100-1000 above which *M* is favored.

**Figure 5:**
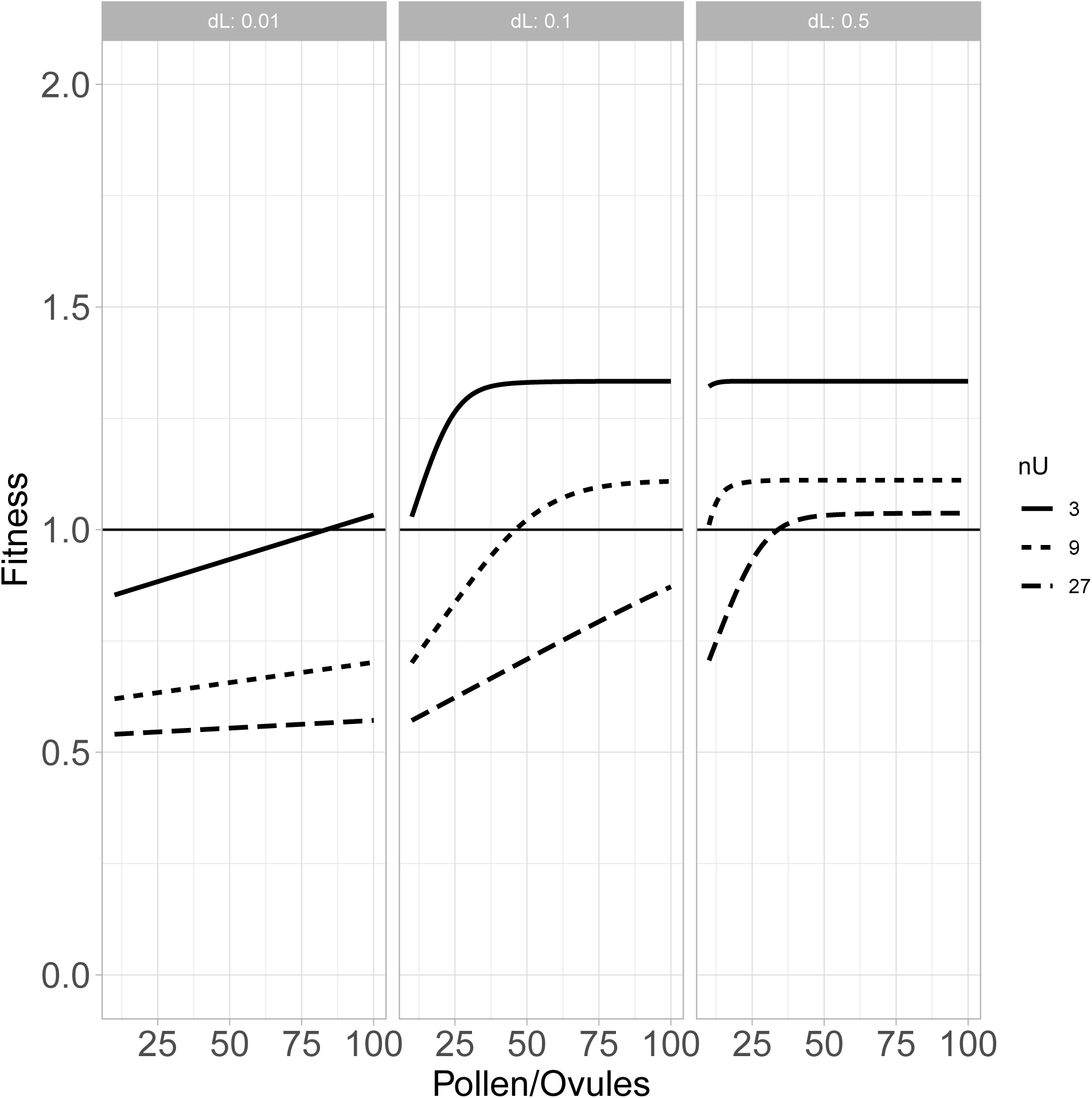
Fitness of a pollen-limited SI mutation. The mutation from the SC intermediate to the new SI haplotype prevents selfing but induces pollen limitation, resulting in a tradeoff. Mutant fitness is calculated relative to that of non-mutant SI haplotypes, which we use as an approximation of population mean fitness. All flowers received equal quantities of pollen drawn proportionally from the pollen pool, and all differences in ovule success were determined by the proportion of that pollen that was compatible. Fitness is plotted against the ratio of pollen received to ovules in a flower: ovule number was held constant at 10, and the pollen supply *G* was varied. Fitness was calculated assuming *n*_*L*_ = 2.

We note that if M is disfavored, this novel specificity goes extinct, and we await a mutation to another novel specificity (or a recurrent mutation to this specificity which could be favored if the number of “‘lucky’ haplotypes’ is, by chance, higher). As such, with every novel specificity, this process repeats itself until one increases in frequency.

### Rescue of doomed haplotypes

Numerical iteration of genotype frequencies when *M* is beneficial shows that *M* will rise to high frequencies and eliminate every haplotype that lacks the key to the new lock (fig. 6). These lost haplotypes include all of the *U* haplotypes along with the lock alleles they carry. Therefore, the overall process results in a collapse in haplotype number at equilibrium rather than expansion. This is consistent with the deterministic analytical results of Bod’ová et al. (2018) which showed that without mutation and with more than seven S-alleles, a novel female incompatibility would always decrease the number of S-alleles when not all haplotypes were initially complete.

**Figure 6:**
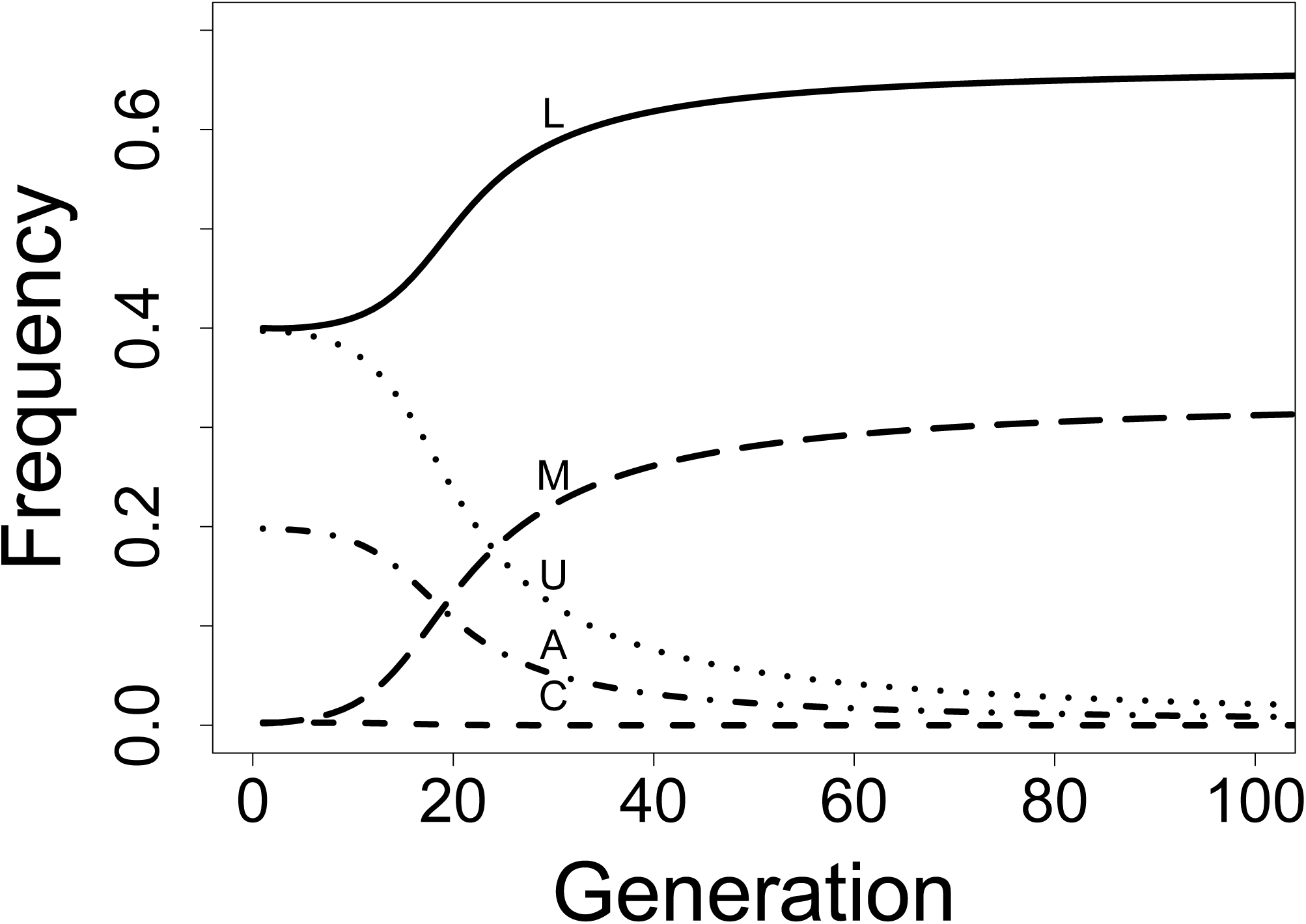
Doomed haplotypes. When the low-frequency self-compatible intermediate (not plotted) acquires a novel RNase by mutation, the resulting haplotype (M) invades. Those haplotypes possessing the key to the new lock (L) also benefit and reach high frequencies, while haplotypes lacking the key (C, U, A) are driven extinct. All trajectories were generated by numerical iteration with parameters *d*_*L*_ = 0.01, 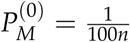, and *n*_*L*_ = *n*_*U*_ = 2.

Multiple haplotypes are now doomed to extinction at equilibrium. However, if a doomed haplotype acquires the key to the new lock before equilibrium is reached, the resulting haplotype’s fitness increases with the frequency of the new lock. This should protect the resulting gene convertant from being driven from low frequencies to extinction. It might therefore be possible for doomed haplotypes to be rescued by gene conversion. If all haplotypes are rescued before any one of them is lost, then the number of lock alleles at equilibrium is one greater than the initial number, and expansion has occurred. Bod’ová et al. (2018) previously noted the role of a rescue-like process in this pathway, though they modeled it through recurrent mutation rather than gene conversion.

The underlying mathematics of this rescue process are similar in form to the evolutionary rescue of a declining population by new mutation. The origin of a new lock is a kind of change in the (genetic) environment, the declining frequency of a doomed haplotype is analogous to a declining population, and the acquisition of the new key by the doomed haplotypes is analogous to the generation of new beneficial mutations. We therefore base our model on Orr and Unckless’ (2008) model of evolutionary rescue by new mutations. There are two components of this model: the supply of beneficial mutations and the probability that any one of them will survive. The probability that the population is rescued is the probability that at least one of these mutations survives. Similarly, the probability that a given haplotype is rescued is the probability that at least one copy adds the new key to its key ring and survives. Orr and Unckless (2008) used a model of survival probability in a population of decreasing size (Otto and Whitlock 1997), which is itself a modification of the branching-process model used by Fisher (1923) and Haldane (1927). However, we modify the original model in a different way because the declining frequency alters the frequency-dependent fitness of the potentially rescuing gene convertants and thus the survival probability of a new mutant increases over time. Thus, rather than varying the population size over time, we determine the time-dependent selection coefficient and the input of new convertants as a function of genotypic frequencies.

In the original model (Orr and Unckless 2008), selection for the rescuing mutation is unconditionally positive. Survival thus implies eventual fixation, and the survival probability equals the fixation probability. In contrast, SI involves balancing selection among haplotypes, so survival does not imply fixation. But since the branching-process approximation is actually the probability of surviving early loss by drift, it can be used in either case. Haldane (1927) defined *s* as the number of additional offspring produced by the mutant haploid individual above the population average. We model survival of a haplotype and its descendants rather than a diploid individual, so we retain Haldane’s assumption of haploidy. In our model, *s* can be re-interpreted as the additional number of descendant copies of a gene convertant above the average for all haplotypes. This advantage *s* depends on the changing frequency of genotypes that accept the gene convertant but reject its ancestor. The survival probability for a single gene convertant is therefore also frequency-dependent, which complicates the calculation of the probability that at least one gene convertant survives.

In the model of Orr and Unckless (2008), the population declines at a predictable rate determined by a standard model of negative population growth. This allowed them to express the number of mutations per generation as a function of time. We cannot do the same with the number of gene conversions per generation because we do not have an explicit formula for genotype frequencies as a function of time. We can, however, numerically iterate recursion equations (see Appendix) to get a trajectory of genotype frequencies. We can then retroactively calculate the expected number of gene conversions each generation in this hypothetical history as a product of the per-individual gene conversion rate, the total number of individuals, and the frequency of heterozygotes for the key to the new lock.

Population size and gene conversion rate have no effect on the survival or rescue probability except through their product, so this product is reported as a single compound parameter *R*_conversion_ (conversions per individual per generation: the exact number of conversions per generation depends on the number of heterozygotes for the key). We include in *R*_conversion_ only those gene conversion events that add a key to a key ring while maintaining SI, but there are at least two other possible outcomes of gene conversion. First, gene conversion might replace a functional F-box paralog with a non-functional or deleted allele: i.e., it might remove a key. Such a conversion event should reduce the siring success of the haplotype and be unconditionally deleterious. Second, gene conversion might induce SC by uniting a key with its complementary lock, as occurred in the previous step at rate *µ*. We do not track these gene convertants in this step because they should be maintained at low frequency at conversion-selection balance. The parameter *R*_conversion_ is therefore smaller than the raw rate of functional gene conversion, which also includes the production of short-lived deleterious convertants.

The probability of rescue depends on the probability of survival of a new gene convertant. We use deterministic expectations for the genotype frequency trajectory and the supply of potentially rescuing gene convertants each generation. All stochasticity in our model therefore arises from whether each potentially rescuing convertant actually survives. We numerically approximated the survival probability of a new gene convertant using

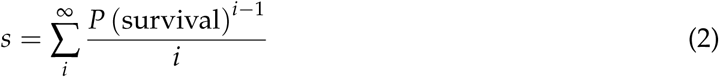

from Haldane (1927). The survival probability of a hypothetical gene convertant was approximated through the following procedure. Proposed survival probabilities were taken from the sequence from 0 to 1 in increments of 0.01. Each generation, we calculated the pollen success of every haplotype and defined *s* as the difference between the fitness of a rescuing gene convertant and the mean fitness. Variation in fitness was completely determined by pollen success because ovule success was equal for all SI haplotypes, save the novel S-allele modeled above. We then chose a survival probability that, when substituted into equation (2) (truncated to the first 100 terms of the sum), resulted in the value of *s* closest to the value calculated from the genotype frequencies. We checked this approximation against simulations for each generation of a common genotype frequency trajectory with *n*_*L*_ = 2, *n*_*U*_ = 2, and a population size of 10,000. The actual survival probability was determined by simulating the survival of a new gene convertant conditional on arising in a given generation. The convertant was judged to have survived if it survived for at least ten generations. The approximation closely followed the simulated results (fig. S1), so we used the approximation for all future calculations.

Once the genotype frequencies, the expected number of gene convertants, and the survival probability of a new gene convertant were known for every generation, we calculated the probability that none of them survived as the product of the complements of the survival probabilities. The complement of this probability is the probability that one doomed haplotype is rescued at least once, assuming the expected number of gene convertants under the expected genotype frequency trajectory. We approximated the probability that a given number of haplotypes were rescued as the probability that each of them was rescued independently. The rescue probabilities are not strictly independent because one rescue event alters the trajectory of genotype frequencies. Rescuing one haplotype essentially creates another lucky haplotype, which could outcompete the remaining doomed haplotypes. However, if the rescues happen in relatively rapid succession, each rescued haplotype will have less effect on subsequent rescues because it is still at low frequency. Orr and Unckless (2008) found that rescue is most likely to occur while the population is still relatively large and mutation supply is highest. That is, there is a short period when the probability of rescue is highest. If rescue in our model is also most likely in a short window (when the doomed haplotypes are still at high frequency), then it is plausible that the multiple rescues occur at similar times and are approximately independent. With this assumption of independence, we generated a probability distribution of the number of surviving haplotypes after rescue or collapse. A natural question is, are large collapses guaranteed, or is there a non-trivial probability of expansion?

For all iterations, we assumed that at least two haplotypes had already acquired the key to the new lock. This is the minimum number for the lock-mutation not to confer ovule-sterility because, if only one haplotype could fertilize it, all ovules carrying the lock-mutation would be fertilized by the same pollen haplotype. The maternal haplotype of the resulting offspring would reject all pollen except the paternal haplotype, which would reject itself. Therefore, when there is exactly one haplotype of class *L*, all ovules carrying the lock mutation grow up to be ovule-sterile adults. In this case, the lock mutation is deleterious rather than neutral and should be rapidly lost. Though this scenario might often occur, it results in no change in the number of S-haplotypes and can be safely ignored in this model.

We calculated the probability that all haplotypes survived for all combinations of *n*_*L*_ = 2, 5, 10, 20, *n*_*U*_ = 2, 5, 10, 20, and *R*_conversion_ in the range of 0.1 – 1 in increments of 0.01 and the range of 1 – 10 in increments of 0.1. We found that the probability that all haplotypes were rescued (the expansion probability) decreased with the number of doomed haplotypes and increased with the population rate of gene conversion *R*_conversion_ (fig. 7). For *n*_*L*_ = 2, the expansion probability rapidly saturated near 1.0 as *R*_conversion_ increased regardless of *n*_*U*_. The expansion probability greatly decreased for *n*_*L*_ = 3 relative to *n*_*L*_ = 2 and continued decreasing gradually for larger values of *n*_*L*_. The effect of *n*_*U*_ was also to decrease the expansion probability, though this effect was less than that of *n*_*L*_.

**Figure 7:**
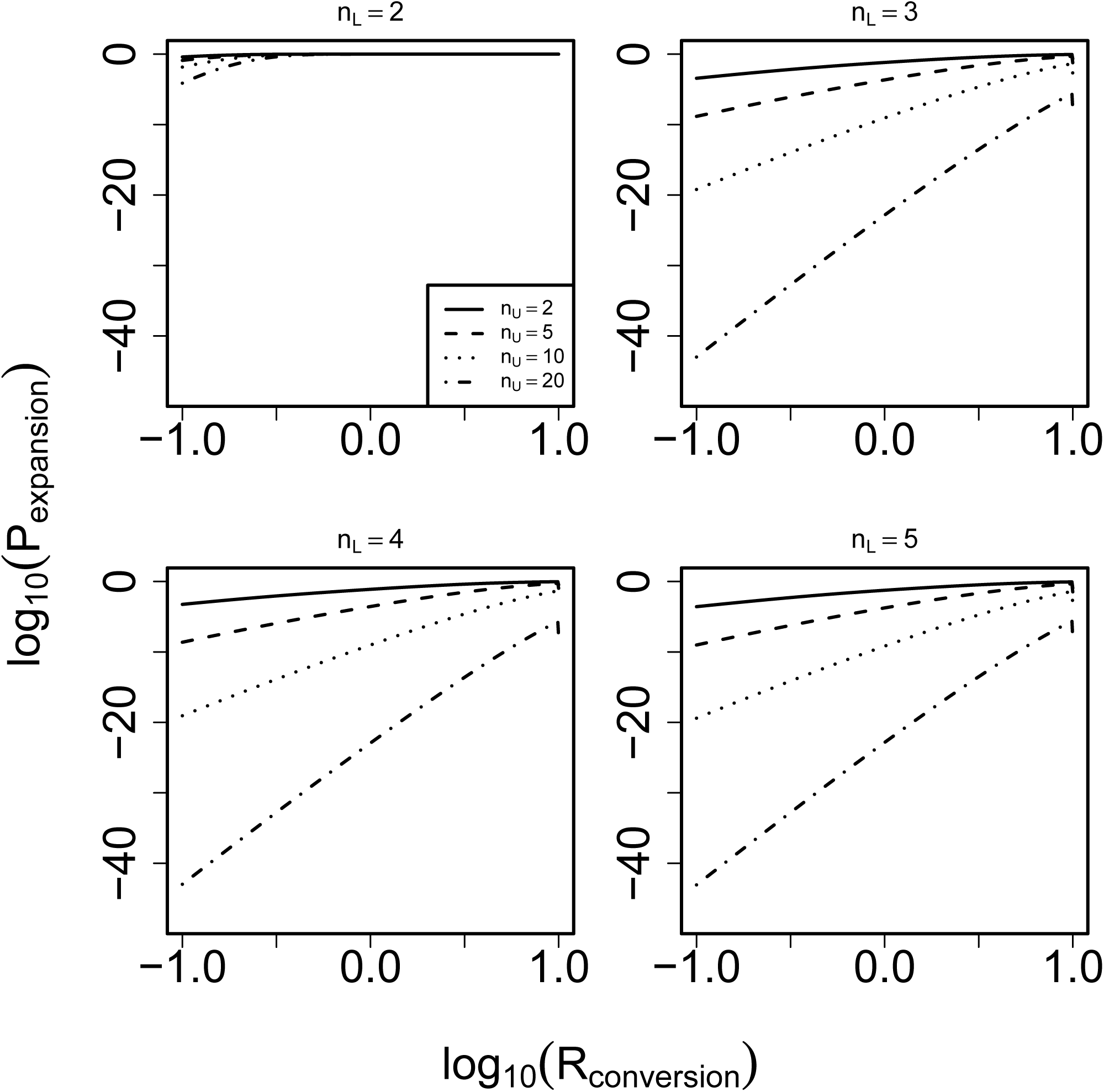
Expansion probability. Each curve represents the probability all doomed haplotypes are rescued for a given number of doomed haplotypes. The probability that all are rescued decreases as the number of doomed haplotypes (*n*_*U*_) increases, and it also usually decreases as the number of “lucky” haplotypes increases. Rescue is very unlikely unless either the population rate of gene conversion (*R*_conversion_) is very high or *n*_*L*_ is small.

The distribution of haplotype number after rescue/collapse responded similarly to *R*_conversion_ for different values of *n*_*L*_, gradually shifting rightward toward maintenance of the initial number of haplotypes, but never reaching high probabilities of expansion for *R*_conversion_ ≤ 0.1 (fig. S2). For *n*_*L*_ = 2 and *n*_*L*_ = 5, the modal outcome was loss of all or almost all doomed haplotypes.

### Long-term behavior

The population has now passed through the final “rescue/collapse” step of the three-step pathway (fig. 3). However, natural populations might traverse this pathway repeatedly, and they must have undergone successive rounds of expansion to produce their current haplotypic diversity. The distribution of haplotype number in nature should therefore be governed by the long-term balance between collapse and expansion. We can make crude predictions for the longterm evolution of haplotype number by treating the diversification process as a Markov chain. Each state of this system is a positive integer representing a number of fully functional haplotypes currently present in the population. The transition probabilities out of each state are given by the distribution of haplotype number after a rescue/collapse event (calculated above). The stationary distribution of this Markov chain represents the probability distribution of haplotype number at expansion-collapse equilibrium. The stable distribution can be obtained without reference to the waiting times between transitions, which we do not model explicitly. In order to create transition matrices of finite size, we enforce a lower reflecting boundary of three haplotypes, the biological minimum that allows any fertilization, and an upper reflecting boundary of 43 haplotypes, near the upper range observed for taxa with SI presumed to function by collaborative non-self recognition (Lawrence 2000). We then iterate each transition matrix for 1000 introductions of a new lock to estimate its stationary distribution.

Our previous rescue probability calculations showed that expansion was likely even at large haplotype numbers when *n*_*L*_ = 2 and *R*_conversion_ ≥ 0.1 (fig. 7). Such large expansion probabilities are necessary to produce the many haplotypes observed in nature. To cover a range of expansion probabilities, we generate transition matrices for *R*_conversion_ = 0.1–0.4 in intervals of 0.1, as well as *R*_conversion_ = 0.01 and 1. We again assume that the initial frequency of each *L* haplotype is low, *d*_*L*_ = 0.01. We also assume that *n*_*L*_ = 2 remains a constant parameter between states, and that all changes in haplotype number are represented by changes in *n*_*U*_. This value of *n*_*L*_ allowed non-trivial expansion probabilities. In reality, the transition matrix should be partly determined by several random variables: the number of *L* haplotypes and the initial frequency of each. The number would be determined by the extent to which gene conversion had already spread the new key before the corresponding lock originated. Likewise, the frequencies would be determined by the outcome of drift between corresponding *L* and *C* haplotypes. We sidestep this complication and assume constant parameters *n*_*L*_ and *d*_*L*_.

We found that the stationary distribution shifted toward greater numbers of haplotypes as *R*_conversion_ increased (fig. 8). At *R*_conversion_ = 0.01, the modal haplotype number was eight, while at *R*_conversion_ = 1, it was 37. In the interval from *R*_conversion_ = 0.2–0.4, the stable distribution was centered around intermediate values near the 20–30 range. At *R*_conversion_ = 0.3, the transition matrix showed that diversification was almost guaranteed at low haplotype numbers, while partial collapses were likely at high haplotype numbers (fig. 9). To visualize the typical history of a population, we also simulated five haplotype-number trajectories at *R*_*conversion*_ = 0.3 for 100 transitions starting at three initial haplotypes. We found that haplotype number increased rapidly for about the first 20 transitions in all simulations, after which the trajectories fluctuated in the range of 20–30 haplotypes (fig. 10).

**Figure 8:**
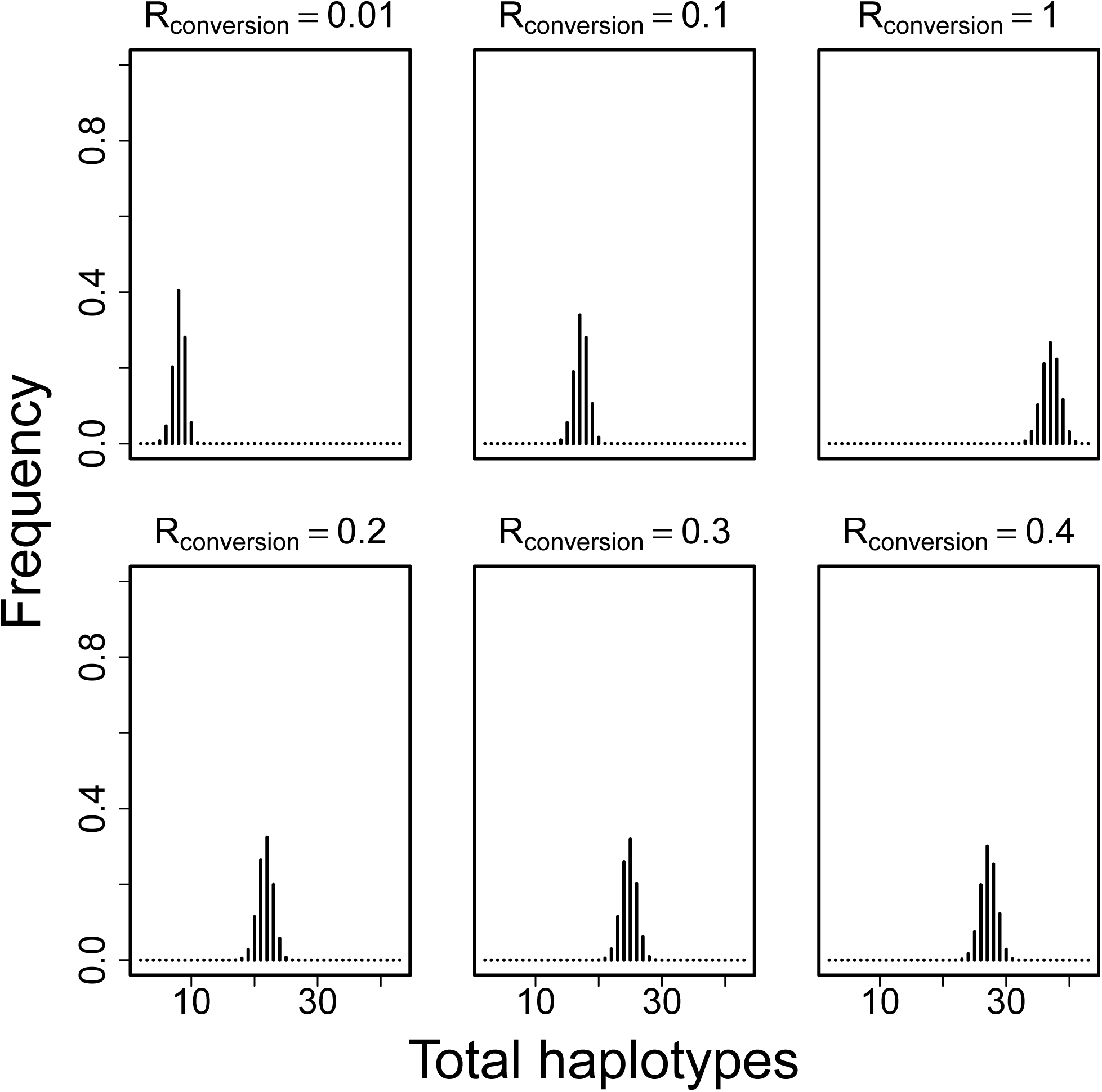
Stable distribution of haplotype number. The stable number of haplotypes increased as *R*_conversion_, the supply of gene convertants, increased. The haplotype number came to exceed the biological minimum near *R*_conversion_ = 0.1, and produced 20–40 haplotypes around *R*_conversion_ = 0.3 or 0.4. For all panels, the Markov chain was run for 1000 steps (expansions or collapses), and *n*_*L*_ = 2. All probability of producing more than 40 incomplete haplotypes (i.e., 43 total haplotypes, counting the *L* and *M* haplotypes) is binned into the 43rd category.

**Figure 9:**
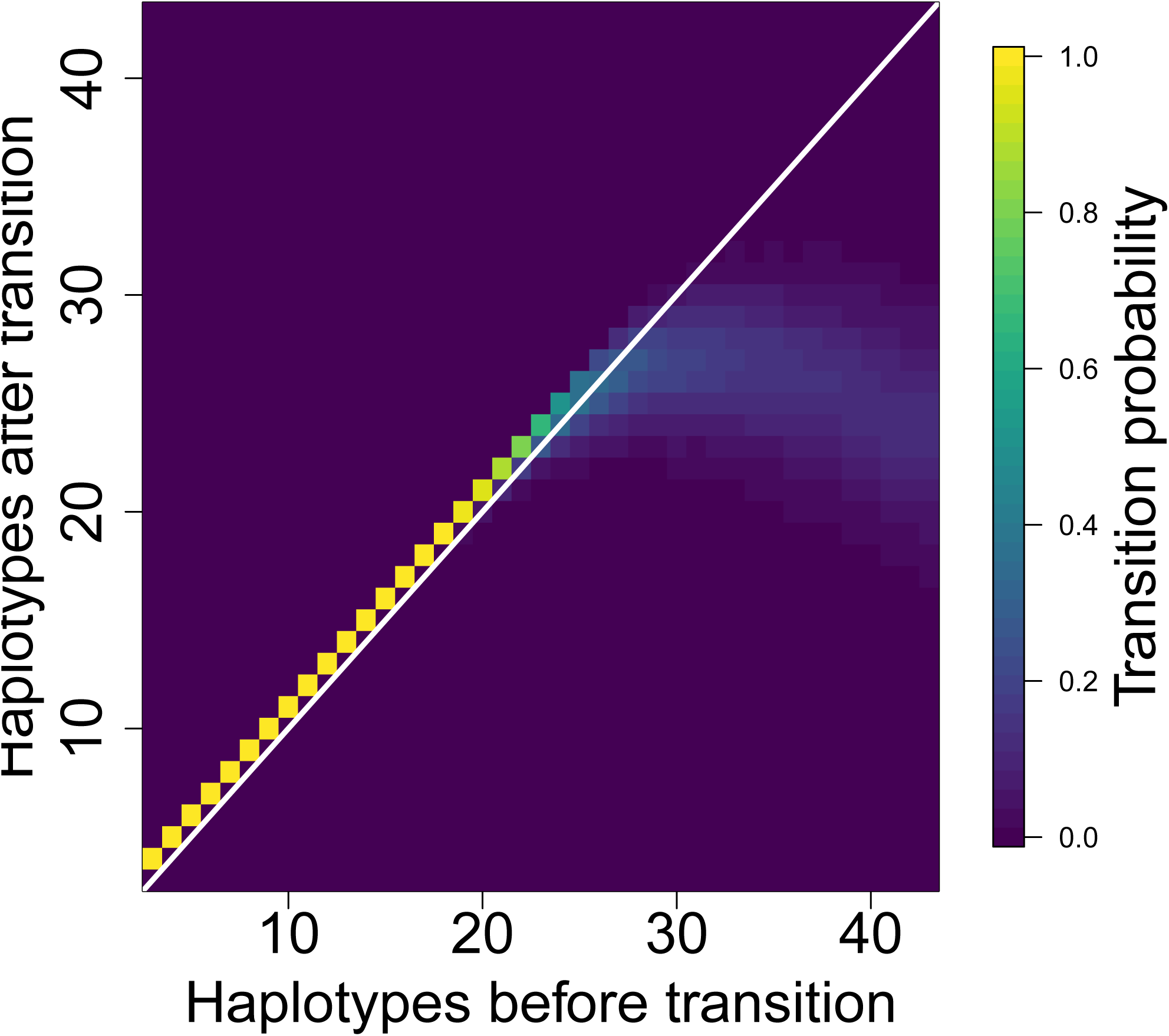
Haplotype number transition matrix for *R*_*conversion*_ = 0.3. The transition probability is from an initial state (horizontal axis) to the next state (vertical axis), where each state is a number of S-haplotypes. For all states, *n*_*L*_ = 2, and only *n*_*U*_ changes between states. The diagonal from bottom-left to top-right (white line) represents no change in haplotype number but could result from either loss of a novel haplotype (i.e., stasis) or maintenance of a novel haplotype and loss of an old one (i.e., turnover). The off-diagonal immediately above that represents gain and maintenance of single novel haplotype. The probability of diversification decreases with increasing haplotype number. It remains greater than 0.5 until *n*_*L*_ + *n*_*U*_ = 18 and remains the most likely outcome until *n*_*L*_ + *n*_*U*_ = 22.

**Figure 10:**
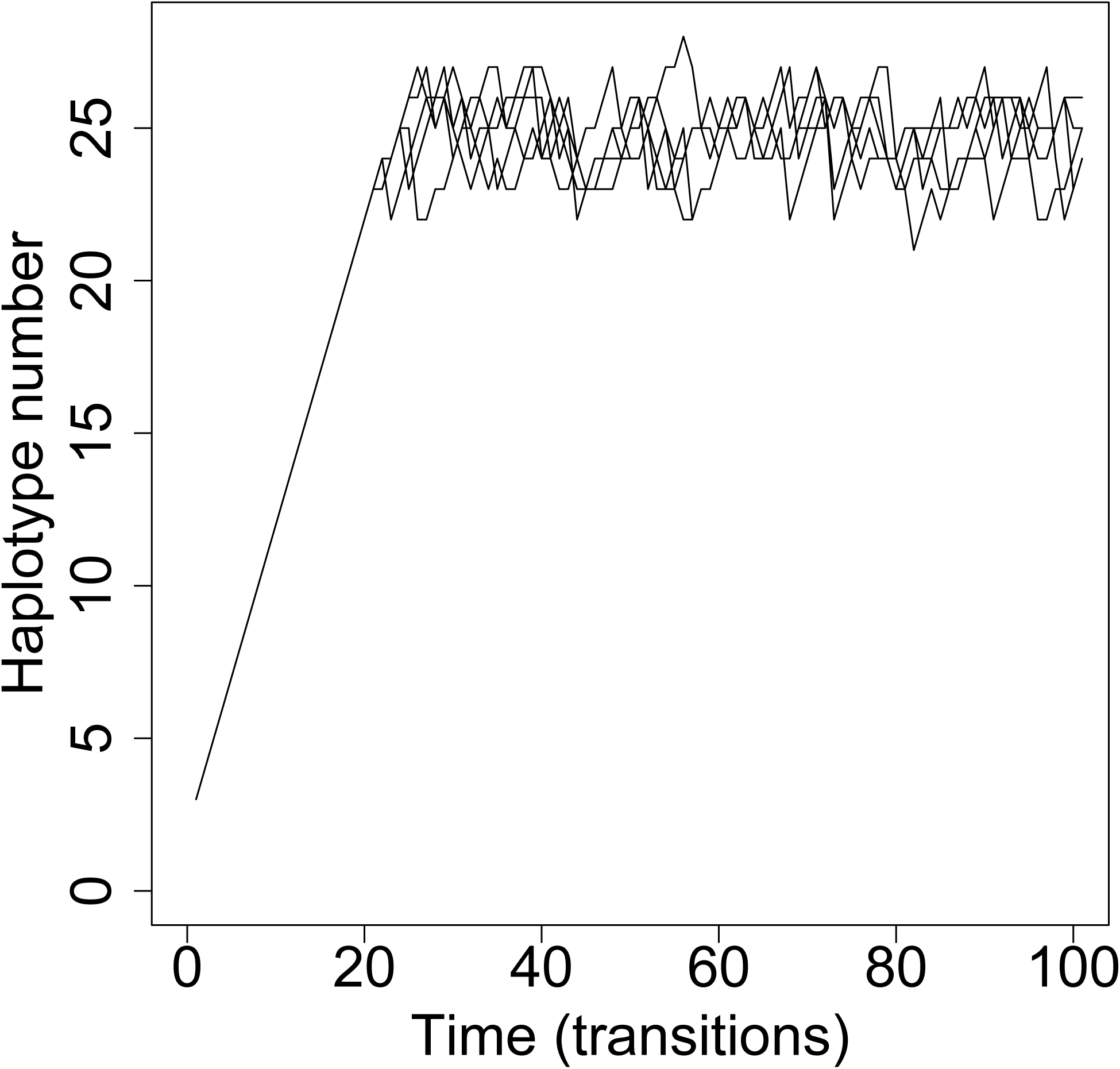
Long-term simulated haplotype number trajectories. Five trajectories were simulated for 100 transitions from the transition matrix for *R*_*conversion*_ = 0.3 and *n*_*L*_ = 2. The initial state was three haplotypes (*n*_*L*_ = 2, *n*_*U*_ = 1). Haplotype number increased rapidly at first and continued to fluctuate after entering the long-term stable range.

The forgoing model only considers changes in the number of S-haplotypes, but another relevant process is the transition from SI to SC. A transition to SC is expected when SC alleles are beneficial and their frequency at conversion-selection balance is at fixation. We repeat the previous model of long-term haplotype number evolution, but additionally assume that any population below a threshold number of S-haplotypes will transition to SC, losing all other S-haplotypes as the SC haplotype rises to fixation. We assume transitions to SC are irreversible, as appears to be the case for the collaborative nonself-recognition system (Igić et al. 2004), and so SC populations leave the Markov process entirely. Thus, any probability mass below this threshold is eliminated every generation, and the remaining probabilities are divided by their sum so they add to one. The resulting distribution describes the probability of each haplotype number conditional on remaining SI. We set this lower threshold at 11 haplotypes, the minimum number for which we found an internal equilibrium frequency of an SC intermediate given *µ* = 0.0003 and *σ* = 0.5 (fig. 4). We also set the starting haplotype number at 11 haplotypes and note that, though these parameters would not allow diversification to 11 haplotypes in the first place, initial diversification could have occurred if parameters were originally even less favorable to SC (e.g., larger *σ*). All long-run distributions were unchanged after adding this threshold except for *R* = 0.01, for which the probability was strongly concentrated at the new minimum of 11 (fig. S3).

## Discussion

We developed a new model of ‘haplotypic rescue’ to estimate the relative probabilities of expansion and contraction in S-haplotype number, calculate the distribution of contraction magnitudes, and predict the long-term evolution of haplotype number. We find that expansion from low haplotype number is possible if gene conversion is frequent or the population is large. When collapses occur instead, they can be large, easily resulting in the loss of the majority of S-haplotypes. A unique prediction of this model is that recently bottlenecked populations should have a “debt” such that, once the appropriate gene conversion event occurs, they will experience a sudden reduction in haplotype number in addition to the reduction directly caused by the bottleneck.

Despite the possibility of collapse, we found that stable distributions of haplotype number within the range of 20–40 were possible in the long term. These numbers are greater than the upper limit of 14 haplotypes found in the model of Bod’ová et al. (2018), and are within the range found in nature. This discrepancy is most likely due to the large supplies of rescuing gene convertants we modeled, and it suggests that arbitrarily many haplotypes can be maintained for a sufficiently large population size or high rate of gene conversion. Population size and S-haplotype number have rarely been estimated for the same populations. However, complete sampling of three populations of *Pyrus pyraster*, a rare woody perennial, revealed 9–25 S-haplotypes per population despite small (8–88 individuals) population sizes (Hoebee et al. 2011). Our model fails to predict the maintenance of so many haplotypes in such small populations at equilibrium, but it is possible that these populations are not at equilibrium. If the waiting times between successive RNase invasions are long, then it is possible that collapses in haplotype number lag far behind reductions in population size. In this case, we should predict that S-haplotype number will drop precipitously in *P. pyraster* in the (possibly distant) future. Alternatively, some populations might be locked at a long-term static haplotype number. For example, we found that a new S-haplotype cannot invade if pollen limitation is too great, in which case neither expansion nor collapse is possible.

A major barrier to the evolution of new S-haplotypes is that a new RNase specificity is expected to reject the majority of pollen in the population because compatibility with the RNase was previously neutral. In the complete absence of compatible pollen, RNase mutations cannot invade because mutant plants cannot set seed. Both Harkness et al. (2019) and “pathway five” in Bod’ová et al. (2018) accounted for this complication by supposing that F-box variants complementary to the RNase preceded the RNase itself, thereby providing a supply of compatible pollen. However, both of these models, as well as all other pathways modeled in Bod’ová et al. (2018) assumed that pollen limitation was binary: plants that received any pollen had the maximum ovule success, and plants that received no pollen had no ovule success. That is, RNase mutants have ovule success equal to that of non-mutants despite rejecting the majority of pollen. Instead, we explore the gradation between total ovule-sterility when pollen is limiting and equal undiminished ovule success when pollen is abundant. We find that a novel RNase can invade despite causing increased pollen limitation as long as compatible pollen is sufficiently abundant. We also find that the relative fitness of an RNase mutation decreases as the number extant RNases increases: the frequency-dependent advantage of rarity is less when the competing RNases are numerous and rare than when they are few and common. This diminishing advantage of RNase novelty could act as a negative feedback that limits how many RNase alleles can be produced in a single population. We therefore predict a negative relationship between pollen limitation and RNase number in nature. Nevertheless, pollen limitation appears to be a surmountable barrier to RNase diversification.

Once the novel RNase invades, S-haplotype number might still either contract or expand. The probability of expansion depends on the population size and gene conversion rate. However, the rate at which gene conversion copies a functional F-box from one haplotype to another is unknown. Per-nucleotide rates of gene conversion have been estimated to be 400 times the rates of crossovers in *Drosophila* (Gay et al. 2007). Assuming a crossover rate of 10^−8^ per meiosis per nucleotide, this yields a gene conversion rate on the order of 10^−6^. A functional gene conversion rate of 10^−6^ would produce few haplotypes (*≈* 8) for populations on the order of 10,000, *≈* 10 − 30 for populations on the order of 100,000, and larger numbers for larger populations. Such populations are large but not unrealistic. Note, however, that relevant quantity is not merely the per-nucleotide rate, but the rate at which a functional F-box is gained. This rate excludes gene conversion events that eliminate a pollen specificity as well as events that fail to affect specificity at all. It is therefore smaller than the per-nucleotide rate of gene conversion, though how much smaller is unknown. The rate at which F-box paralogs are copied would constrain not only the probability of rescue (through *R*_*conversion*_) but also the input of SC gene convertants (through *µ*). It might be possible to estimate the rate of functional gene conversion indirectly. To perform such an estimate, a study would need to measure both the frequency and seed set of wild SC individuals relative to SI individuals in a predominantly SI population. Since the equilibrium frequency of SC haplotypes is determined by their ovule fitness and the gene conversion rate, it should be possible to estimate the gene conversion rate given the observed frequency and fitness of SC.

Other than large populations and rapid gene conversion, another possible explanation of high S-haplotype diversity is that gene flow buoys species-wide diversity despite local contractions. Uyenoyama et al. (2001) found that, similar to our results, S-allele number was unlikely to expand within a single population because coexistence was rarely possible between a mutant SC intermediate and its ancestral haplotype. They proposed that expansion might instead occur through local turnover in S-alleles followed by introduction of the novel alleles into the broader metapopulation. However, this model assumed that incompatibility operates through self-recognition, in which introgressed S-alleles gain an advantage because they are compatible with resident plants that do not recognize them as self. It is thus not obvious whether the conclusion translates to nonself-recognition. Castric et al. (2008) found less divergence between putative shared ancestral variants of S-locus genes in *Arabidopsis lyrata* and *A. halleri* than between non-S orthologs, consistent with elevated introgression at the S-locus in this self-recognition system. An obvious empirical question is whether this result could be replicated in a species with a nonself-recognition system. What would our model predict? Under collaborative nonself-recognition, each novel RNase requires a corresponding F-box. A novel introgressed RNase allele will not necessarily be detoxified by local F-box proteins, and it might thus inflict ovule-sterility. If there is insufficient compatible pollen to support the migrant haplotype, SI might act as a barrier to introgression. This barrier could be overcome, but it would require either that the key corresponding to the foreign lock already exists in the population (e.g., as a dual-function key or as a segregating neutral variant), or that there is sufficient migration to supply the corresponding foreign keys. Given these complications, the impact of gene flow on the diversification / collapse of S-allele diversity and the impact of S allele diversity on hybridization and introgression in collaborative nonself recognition based SI require additional research.

We found that the expansion probability greatly decreased as the number of complete haplotypes increased, a second form of negative feedback in addition to that imposed on the invasion of the RNase in the first place. Greater numbers of complete haplotypes reduce the expansion probability in at least two ways. First, by increasing the number and thus the frequency of fit haplotypes, they increase the mean siring success. When compatibility with the new RNase is more common, the competitive advantage of this compatibility is reduced. Second, with more complete haplotypes, the equilibrium frequency of the novel RNase is lesser. These two effects reduce the advantage of compatibility with the new RNase and thus the survival probability of the rescuing gene convertant. This essential parameter of the model, the initial number of haplotypes possessing the F-box complementary to the novel RNase specificity, has not been systematically estimated. The existence of the complementary F-box on multiple backgrounds is a prerequisite for both expansion and collapse. The prevalence of pre-existing complementary F-boxes in a population could be determined by experimentally applying pollen to plants carrying a novel, functional RNase allele. Successful fertilization would be consistent with a pre-existing F-box complementary to the RNase. An alternative explanation for fertilization, the loss of RNase function in the dam, could be ruled out with the appropriate control: F-box knockouts should be compatible with RNase loss-of-function mutants but not with novel RNase alleles. Novel RNase alleles could be generated by mutagenesis, or they could be introduced from distantly related populations transgenically or through introgression.

Besides bounding the parameters of the model, empirical research could also test for evidence of recent historical collapses. Consider a pair of allopatric populations that initially shared all S-haplotypes. However, a novel RNase has recently invaded Population A and eliminated several pre-existing haplotypes, resulting in a contraction in haplotype number. This RNase never arose in Population B, which retains the original complement. Every remaining haplotype in Population A other than the novel haplotype itself should carry the F-box complementary to the novel RNase. Of course, these vestigial F-boxes would eventually become pseudogenized, but they might still be functional shortly after the collapse. In Population B, several haplotypes should either be polymorphic for the novel RNase’s complementary F-box or should lack it entirely. This situation could be tested by reciprocally crossing individuals from the two populations. A substantial proportion of B pollen should be rejected by recipients in Population A: a proportion in excess of that predicted by the number and frequency of shared S-haplotypes. A potentially dramatic test would be to introgress the novel RNase from Population A to Population B. In the short run, the siring success of all haplotypes not already compatible with the new RNase would decrease. In the long run, the frequency of these unlucky haplotypes would decrease to extinction. That is, we predict that S-haplotype collapse should be contagious between closely related populations, even if both populations are SI.

This work builds on that of Bod’ová et al. (2018), who model invasion and coexistence conditions for S-haplotypes and also directly simulate the dynamics of the diversification process. One of their key insights from their coexistence model is that, if some haplotypes are more complete than others (i.e., possess more functional pollen specificities), the less complete haplotypes can be driven to extinction at equilibrium because of their inferior pollen success. They show that a new S-haplotype cannot coexist indefinitely with all pre-existing haplotypes except in the limited cases that either all haplotypes are complete or exactly one haplotype is complete. They note, however, that even transient coexistence might provide a window in which further mutations might make all haplotypes complete and thus capable of permanent coexistence. This rescue process is implicitly incorporated in their stochastic simulations, and we model it analytically in this study. Their coexistence model also shows that SC haplotypes will reach fixation except for high values of inbreeding depression (≥ 0.75), which informs our focus on intense inbreeding depression. Bod’ová et al. (2018) consider two kinds of pathways: one that maintains SI throughout and several that go through an SC intermediate. There is good reason to believe that, although the pathway maintaining SI is traversed moderately often in the simulations of Bod’ová et al. (2018), this pathway should be rare in nature. This is because when an RNase mutation occurs on an SI background, it reduces the amount of pollen accepted but does not provide any benefit. Such a mutation is neutral if ovule success is not limited by the supply of compatible pollen (Bod’ová et al. 2018) but deleterious if it is. In contrast, we have shown that a pathway through an SC intermediate is possible even with the barrier of pollen limitation because the RNase mutation prevents self-fertilization. Finally, we have followed up on the suggestion by Bod’ová et al. (2018) to incorporate gene conversion of F-box paralogs as hypothesized by Kubo et al. (2015) and Fujii et al. (2016). Although recurrent gene conversion in our model functions similarly to recurrent mutation in their model (Bod’ová et al. 2018), an important difference is that functional gene conversion can only occur in heterozygotes for an F-box locus. We show that, despite this limitation on the supply of potentially rescuing variants, diversification is still possible.

Even though we assume maximal inbreeding depression throughout, we find that populations are susceptible to invasion and fixation of SC haplotypes unless selfing rates are also high (fig. 4). The possibility of transition to SC does not seem to affect the distribution of haplotype numbers of those populations that remain SI unless the stable distribution crosses the minimum for maintaining SI (fig. S3), likely because there is a strong upward pull when haplotype numbers start below the stable range (fig. 9, fig. 10). However, populations can only diversify from low to moderate numbers of haplotypes in the first place if selfing rate is high. Thus, though diversification is possible through this pathway alone, transitions to SC might be very common at low haplotype numbers in nature. A trait that is frequently lost but rarely or never regained can still common among many species if species with the trait speciate more frequently or species without the trait go extinct more frequently. This macroevolutionary process, species-level selection, is hypothesized to maintain SI among species despite frequent and possibly irreversible transitions to SC (Igić et al. 2008; Goldberg et al. 2010). But if diversification is precarious at low numbers, how have high numbers been reached? And if high inbreeding depression and selfing rate are required, why would a population not have either purged the inbreeding depression or evolved simpler mechanisms of reducing selfing? Future models of the origin of the polyallelic S-locus ought to answer these questions, but the present model is best viewed as one of continued diversification rather than one of origin.

SI has developed into a rich study system through the continued interaction between theory and empirical research. Experimental demonstration of the genetic control of SI (East and Mangelsdorf 1925) and field research showing the number of S-alleles in natural populations (Emerson 1938, 1939) inspired theoretical explanations of the balancing selection capable of maintaining this diversity (Wright 1939; Charlesworth and Charlesworth 1979). The theoretical potential for balancing selection indicated the S-locus as a candidate for long-term polymorphism, and S-allele phylogenies confirmed this possibility (Igić and Kohn 2001; Steinbachs and Holsinger 2002). The discovery of the fine-scale genetic basis of nonself-recognition (Kubo et al. 2010, 2015) necessitated new theoretical explanation of the expansion process, and this theory now points to the unanticipated possibility of S-allele collapse through runaway gene convertants.

## Supporting information

All scripts

## Appendix genotype frequency recursions

The following genotype frequency recursions governed changes in the frequencies of the classes of SI haplotypes. They were used to produce the trajectories leading to extinction of doomed haplotypes (fig. 6), which were in turn used to calculate rescue probabilities. SC haplotypes were assumed to be rare and were neglected in these recursions. Each *P*_*XY*_ represents the frequency of a diploid genotype consisting of one haplotype of class *X* and one of class *Y*. The genotype frequency recursions are

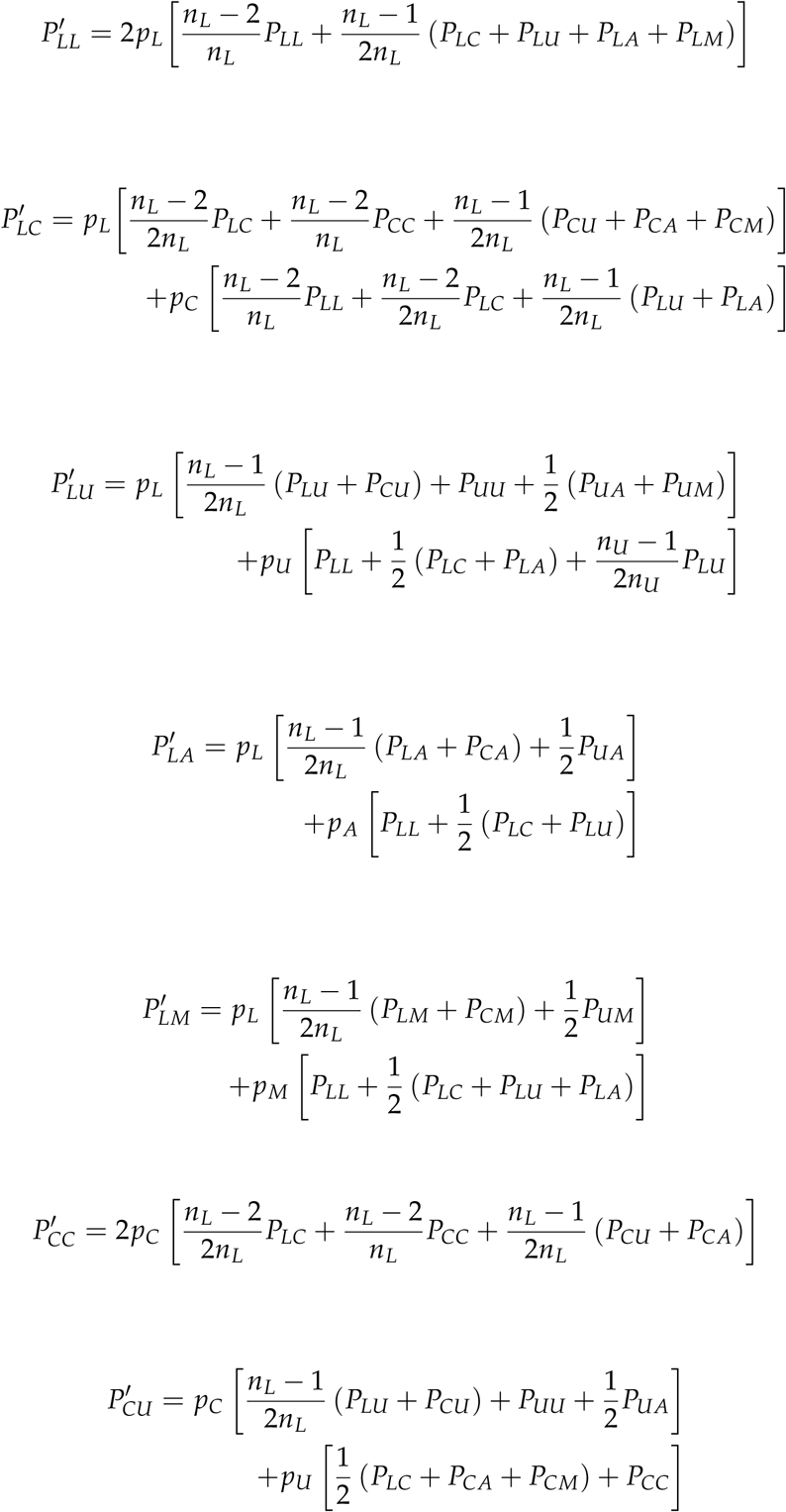

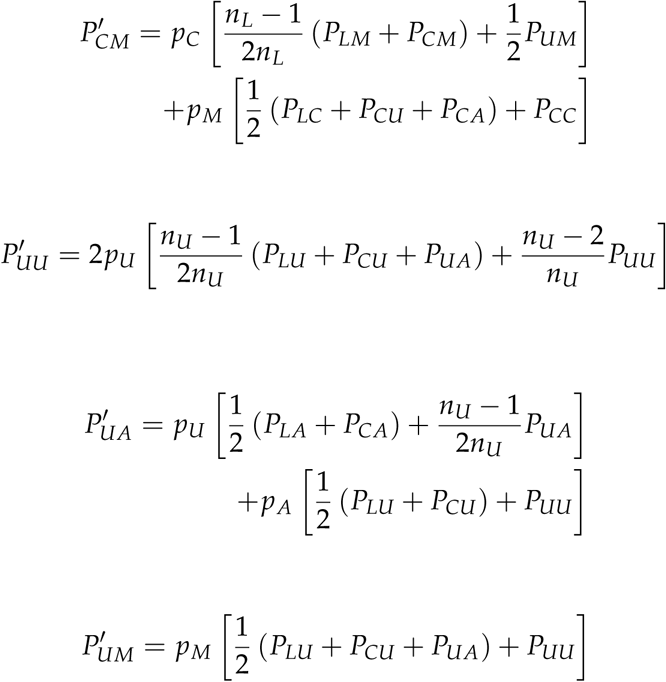

---

**Figure S1:**
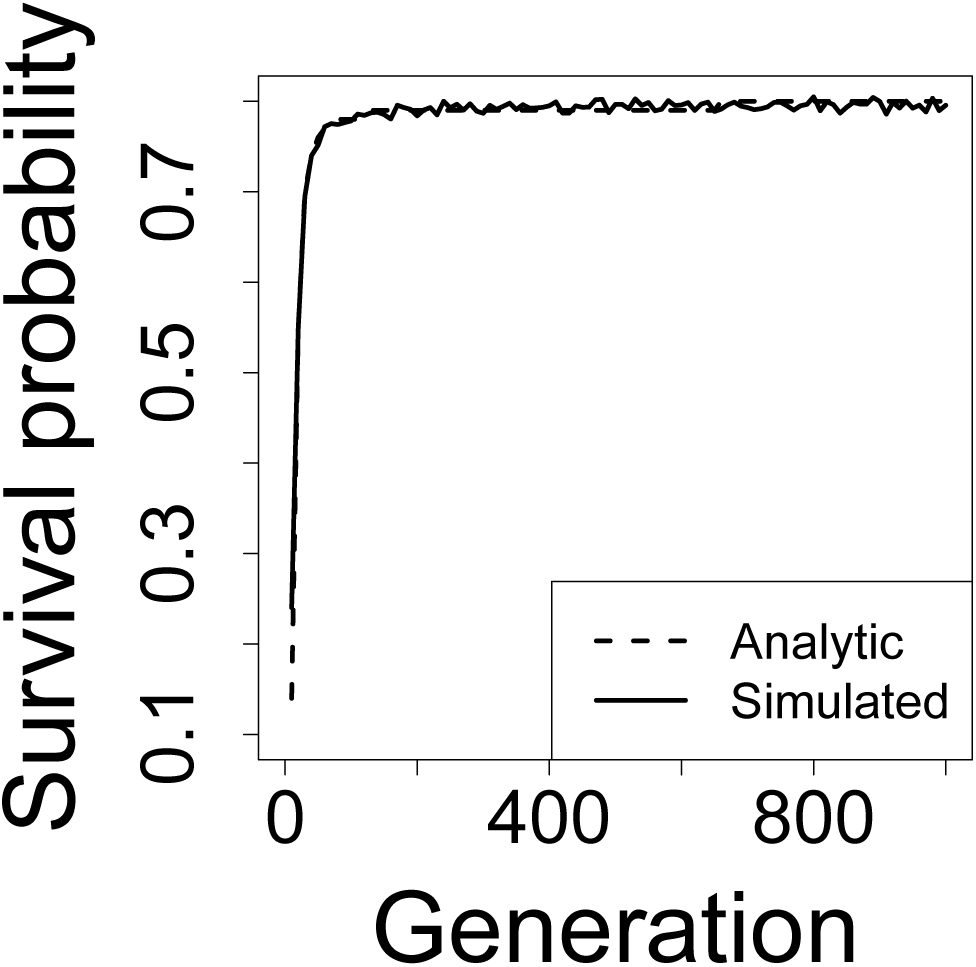
Survival probability of a new gene convertant. The history of a population of 10,000 individuals with *n*_*L*_ = 2, *n*_*U*_ = 2, during the rescue phase was first determined by iterating the recursion equations. Survival probability was determined each generation using Haldane’s series expansion (analytic) or by simulating survival of a new gene convertant for 10 generations over 10,000 replicate simulations (simulated).

**Figure S2:**
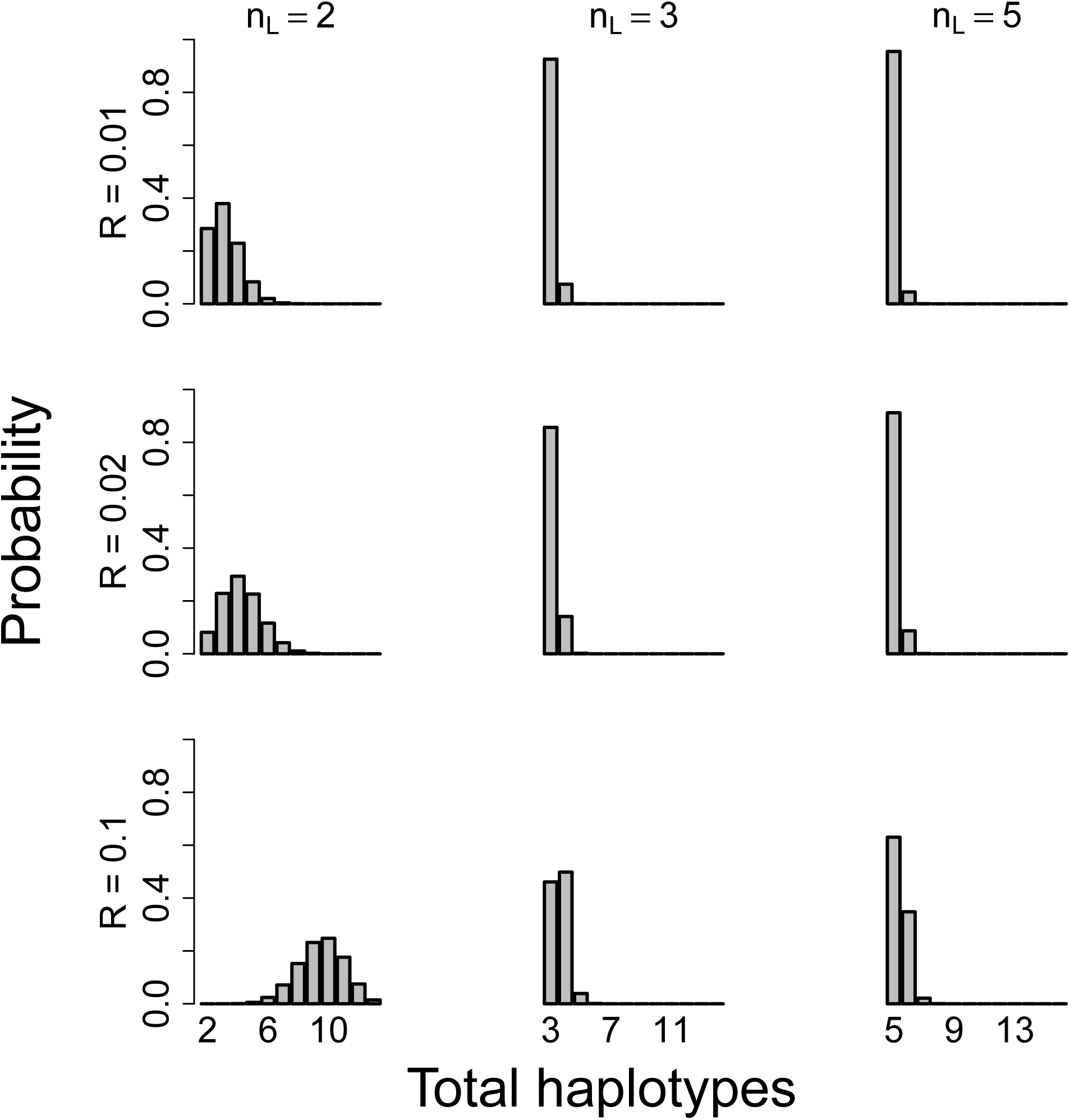
Distribution of haplotype number after a single rescue/collapse event. This probability distribution of outcomes is conditional on the population rate of gene conversion *R*_*conversion*_, and on the current numbers of lucky *n*_*L*_ and unlucky *n*_*U*_ haplotypes. For all panels, *n*_*U*_ = 10. Expansion only occurs when the final number of haplotypes is *n* + 1 (the rightmost bin in each histogram). For *n*_*L*_ > 2 and small population gene conversion rates (*R*_conversion_), the most likely outcomes is either the loss of all doomed haplotypes or the loss of all but one.

**Figure S3:**
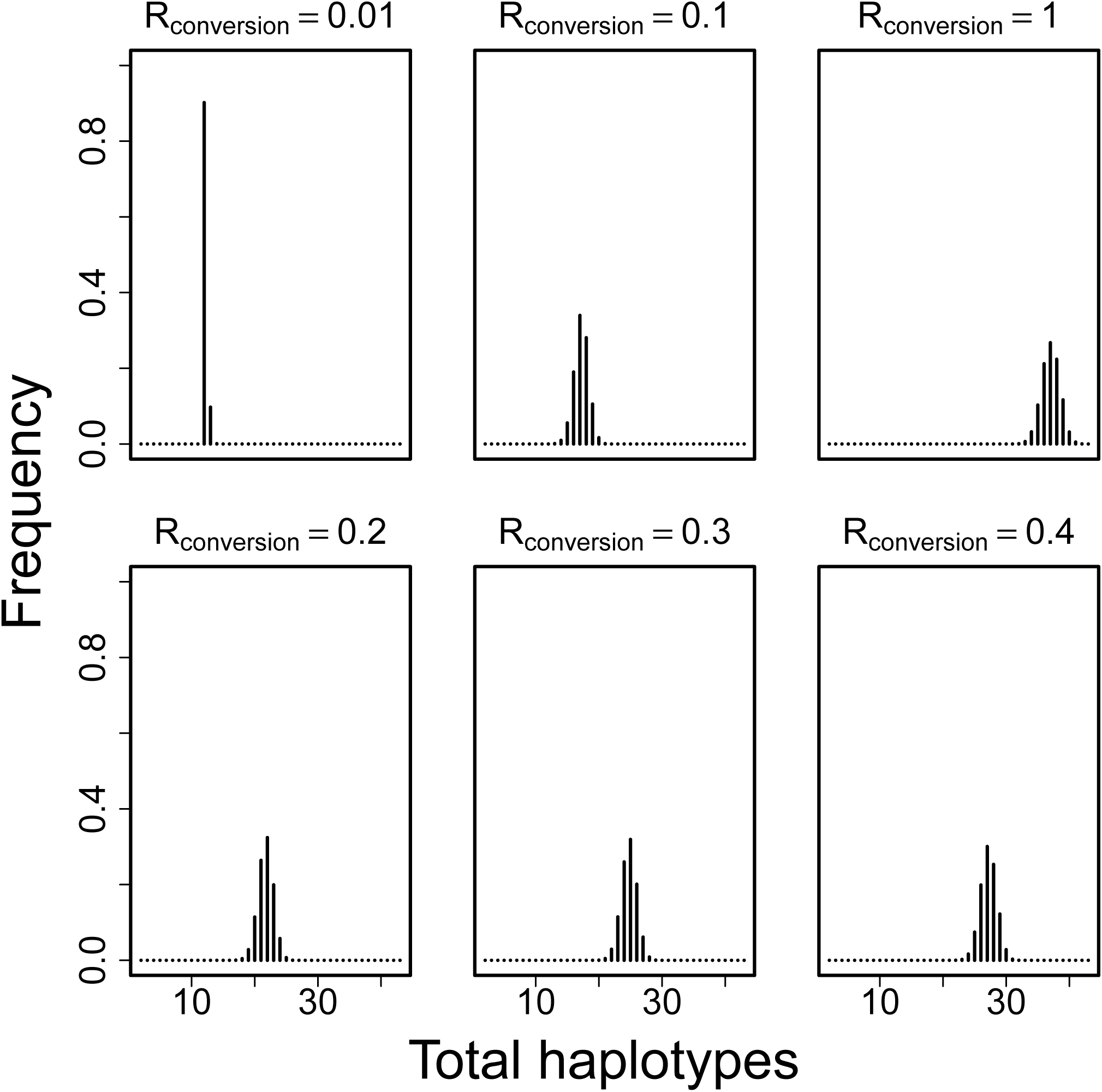
Stable distribution of haplotype number with lower threshold. Figure is identical to (fig. 8), except that all probability mass below 11 haplotypes was removed each generation and the remaining probabilities were divided by their sum.

