## Supplementary material for "Diversification or collapse of self-incompatibility haplotypes as a rescue process": All scripts: analytic_survival.R

```
# This script determines the survival probability of a new gene convertant using Haldane's approximation.

source('rnase_functions.R')

# Load the expected trajectory.

args <- commandArgs(trailingOnly=TRUE)

L <- eval(parse(text=args[1]))
U <- eval(parse(text=args[2]))
luck_frac <- 1
spite_frac <- 1
my_generations <- 4000

trajectory_name <- paste('../data/', 'L=', L, '_U=', U, '_split_traj.csv', sep = '')
expected_trajectory <- as.matrix(read.csv(trajectory_name, row.names = 1))

# Initialize the population from those parameters.
my_genos <- generate.genos(L,U, monster = T)
my_pollen_prod <- generate.pollen.production(my_genos)
my_hap_comp <- generate.haplo.compatibility(my_genos, monster = T)
my_dip_comp <- generate.diplo.compatibility(my_hap_comp, my_genos)

analytic_longitudinal <- vector(length = nrow(expected_trajectory))
for(i in 1:length(analytic_longitudinal)){
  current_s <- get.sel(expected_trajectory[i,], my_pollen_prod, my_dip_comp, 'L3')
  analytic_longitudinal[i] <- series.solver(current_s, seq(from = 0, to = 1, by = 0.01), 100)
}

survival_name <- paste('../data/', 'L=', L, '_U=', U, '_survive.csv', sep = '')
# write.table(t(c('start_gen','survival_probability')), file = survival_name, sep = ',', col.names = F, row.names = F)
write.table(analytic_longitudinal, file = survival_name, sep = ',', col.names = F, row.names = T)

################
# REPEAT FOR A #
################

# We need a separate survival probability for A.
# A gene convertant of A produces essentially another L haplotype, except that it can fertilize all L and U but not A.
# To emulate this, we replace A with a new RNase class containing a U (the unrescued A) and an L (the rescued A).
# I'm pretty sure that would have exactly the same mechanics.

# First, create genotypes and compatibilities.
# Start by making genotypes on the assumption that there is ONE MORE class of unlucky haplotypes than there really is.
modified_genos <- generate.genos(L, U+1, monster = T)
modified_pollen_prod <- generate.pollen.production(modified_genos)
modified_hap_comp <- generate.haplo.compatibility(modified_genos, monster = T)
modified_dip_comp <- generate.diplo.compatibility(modified_hap_comp, modified_genos)

# Now, whatever genotypes match the modified trajectory, fill them in.
# But then replace all As with Us of the extra type.
modified_trajectory <- matrix(nrow = nrow(expected_trajectory), ncol = nrow(modified_genos))
modified_trajectory[] <- 0
colnames(modified_trajectory) <- paste(modified_genos[,1], modified_genos[,2], sep = '_')
for(i in colnames(expected_trajectory)){
  modified_trajectory[,i] <- expected_trajectory[,i]
}

# Loop through each A-bearing genotype and move the contents to the genotype that bears fake_U but is otherwise identical.
fake_U <- paste('U', L+U+1, sep = '')
# Need haplotype names to weed out lookalikes.
haps <- rownames(modified_hap_comp)
for(i in colnames(modified_trajectory)){
  if(grepl('A', i)){
    # Find out the identity of the other hapltoype in this genotype besides the A.
    # Do this by deleting As and _s.
    other_haplo <- gsub('_', '', gsub('A', '', i))
    # Find the haplotype that contains fake_U and other_haplo.
    # for A_fake_U, fake_U and other_haplo refer to the same haplotype.
    # Therefore, anything with fake_U and contains both fake_U and other_haplo when A_fake_U is reached.
    # Therefore, do no replacement when fake_U and other_haplo refer to the same haplotype.
    # Regular expressions are dangerous.
    # If the first digit of fake_U or other_haplo is shared by multiple haplotypes of the same class,
    # they will accidentally match a grep for fake_U or other_haplo.
    # Be therefore vigilant against lookalikes.
    lookalike_haps <- haps[(grepl(fake_U, haps) & fake_U != haps) | (grepl(other_haplo, haps) & other_haplo != haps)]
    lookalike_dips <- paste(modified_genos[,1], modified_genos[,2], sep = '_')[(modified_genos[,1] %in% lookalike_haps) | (modified_genos[,2]) %in% lookalike_haps]
    if(fake_U == other_haplo){}
    else{
      replacement <- colnames(modified_trajectory)[grepl(fake_U, colnames(modified_trajectory)) & grepl(other_haplo, colnames(modified_trajectory))]
      # Remove all lookalikes from replacement.
      replacement <- replacement[!(replacement %in% lookalike_dips)]
      modified_trajectory[,replacement] <- modified_trajectory[,i]
      modified_trajectory[,i] <- 0
    }
  }
  else{}
}

# Now that the modified trajectory is known, calculate the survival probability of a hypothetical fake_L.
fake_L <- paste('L', L+U+1, sep = '')

analytic_longitudinal_A <- vector(length = nrow(modified_trajectory))
for(i in 1:length(analytic_longitudinal_A)){
  current_s <- get.sel(modified_trajectory[i,], modified_pollen_prod, modified_dip_comp, fake_L)
  analytic_longitudinal_A[i] <- series.solver(current_s, seq(from = 0, to = 1, by = 0.01), 100)
}

# Now write the survival probability for A convertants.
survival_name_A <- paste('../data/', 'L=', L, '_U=', U, '_survive_A.csv', sep = '')
write.table(analytic_longitudinal_A, file = survival_name_A, sep = ',', col.names = F, row.names = T)
```
