## Supplementary material for "Diversification or collapse of self-incompatibility haplotypes as a rescue process": All scripts: calculate_rescue.sh

```
#!/bin/bash
for L in 5; do
	for U in 1 2 3 4 5 6 7 8 9 10 11 12 13 14 15 16 17 18 19 20 21 22 23 24 25 26 27 28 29 30 31 32 33 34 35 36 37 38 39 40; do
		for N in 1000; do
			for CONV in 0.00001 0.0001 0.001 0.01 0.1; do
				Rscript rescue_from_trajectory.R $L $U $N $CONV
			done
		done
	done
done
```
