## Supplementary material for "Diversification or collapse of self-incompatibility haplotypes as a rescue process": All scripts: calculate_survival.sh

```
#!/bin/bash
for L in 5; do
	for U in 1 2 3 4 5 6 7 8 9 10 11 12 13 14 15 16 17 18 19 20 21 22 23 24 25 26 27 28 29 30 31 32 33 34 35 36 37 38 39 40; do
        Rscript analytic_survival.R $L $U
    done
done
```
