## Supplementary material for "Diversification or collapse of self-incompatibility haplotypes as a rescue process": All scripts: check_sim.R

```
# This script compares the survival probabilities calculated analytically with those determined by simulation.

source('rnase_functions.R')

# Load the expected trajectory.

expected_trajectory <- as.matrix(read.csv('../data/L=2_U=2_split_traj.csv', row.names = 1))

L <- 2
U <- 2
luck_frac <- 1
spite_frac <- 1
my_generations <- 4000
#my_N <- 10000

# Initialize the population from those parameters.
my_genos <- generate.genos(L,U, monster = T)
my_pollen_prod <- generate.pollen.production(my_genos)
my_hap_comp <- generate.haplo.compatibility(my_genos)
my_dip_comp <- generate.diplo.compatibility(my_hap_comp, my_genos)
my_geno_dist <- sim.initialize(my_genos, luck_frac = luck_frac, spite_frac = spite_frac)
names(my_geno_dist) <- paste(my_genos[,1], my_genos[,2], sep = '_')

# Examine all the hapltoypes at a given instant.
sels <- vector(length = 10)
names(sels) <- c('A','C1','C2','L1','L2','L3','L4','S','U3','U4')
for(i in 1:length(sels)){
  sels[i] <- get.sel(expected_trajectory[2,], my_pollen_prod, my_dip_comp, names(sels)[i])
}

# Examine how the survival probability changes through time.
# inferred_longitudinal <- vector(length = nrow(expected_trajectory))
# for(i in 1:length(inferred_longitudinal)){
#   # current_s <- get.sel(expected_trajectory[i,], my_pollen_prod, my_dip_comp, 'L3')
#   current_s <- realized.sel(my_genos, expected_trajectory[i,], my_pollen_prod, my_dip_comp, my_N)
#   inferred_longitudinal[i] <- series.solver(current_s, seq(from = 0, to = 1, by = 0.01), 100)
# }

analytic_longitudinal <- vector(length = nrow(expected_trajectory))
for(i in 1:length(analytic_longitudinal)){
  current_s <- get.sel(expected_trajectory[i,], my_pollen_prod, my_dip_comp, 'L3')
  # current_s <- realized.sel(my_genos, expected_trajectory[i,], my_pollen_prod, my_dip_comp, my_N)
  analytic_longitudinal[i] <- series.solver(current_s, seq(from = 0, to = 1, by = 0.01), 100)
}

pdf('../figures/sim_check.pdf')
# Plot the analytic survival probability versus the simulated survival probability.
par(mar = c(8,9,4,2), mgp = c(6, 2, 0), cex.axis = 3, cex.lab = 4)
# par(cex.axis = 2,
#     cex.lab = 2,
#     #cex = 2,
#     mar = c(5,5,4,2))
sim <- read.csv('../data/sim_survival.csv', header = F)
#detailed <- sim[201:nrow(sim),]
# Remember that analytic_longitudinal calculates survival probability every generation,
# whereas sim calculates every ten generations.
# Therefore, we want to subset analytic_longitudinal
plot(analytic_longitudinal, type = 'l', xlab = 'Generation', ylab = 'Survival probability',
     ylim = c(0,0.8),
     xlim = c(0,1000),
     lwd = 3,
     lty = 2)
points(y = sim[,2], x = sim[,1], type = 'l',
       lwd = 3,
       lty = 1)
legend('bottomright', legend = c('Analytic', 'Simulated'), lty = c(2, 1), lwd = 3, cex = 2)
#points(analytic_longitudinal, type = 'l', col = 'red')
dev.off()


library(tidyverse)
tibble( Generation = c(sim[,1],10:length(analytic_longitudinal)) ,
        `Surival Probability` = c(sim[,2], analytic_longitudinal[-(1:9)]),
        type = rep(c("Simulation", "Analytical"), 
                   times = c(nrow(sim), length(analytic_longitudinal)-9))) %>%
  ggplot(aes(x = Generation, y = `Surival Probability`, color = type)) +
  geom_line(data = . %>% filter(type != "Simulation"), lwd = 2, alpha = .5)+
  geom_point(data = . %>% filter(type == "Simulation"))+
  theme_classic()+
  scale_x_continuous(trans = "log10",limits = c(10,1000))+
  scale_y_continuous(trans = "log10",limits = c(.1,1), breaks = seq(.1,.8,.1))
#  coord_cartesian(xlim = c(0,1000))


#write.table(t(c('start_gen','survival_probability')), file = '../data/expected_survival.csv', sep = ',', col.names = F, row.names = F)
#write.table(analytic_longitudinal, file = '../data/expected_survival.csv', append = T, sep = ',', col.names = F, row.names = T)
```
