## Supplementary material for "Diversification or collapse of self-incompatibility haplotypes as a rescue process": All scripts: conversion_sim.R

```
# Load packages
library('gtools')

# This script is intended to simulate the diversification of GSI S-RNase S-haplotypes under the collaborative nonself-recognition model.

# I assume that the S-locus is a single locus, even though it is technically composed of one S-RNase locus and multiple SLF paralogs.
# Each combination of an RNase specificity and a complement of SLF specificities is treated as an allele at this locus.

# The first event, invasion of a new RNase, is governed by a deterministic model.
# A new RNase mutation invades a population that already contained SLF variants that could detoxify it.
# No further mutations occur.

# The simulation begins with a population that has already been invaded by a new RNase, and that has attained the analytically derived equilibrium.
# Each individual heterozygous for the ability to detoxify the new RNase undergoes gene conversion with some probability.
# This kind of gene conversion produces a single gamete with the larger SLF complement, and the parental RNase that cannot be detoxified by this SLF complement.
# There are other possible products of gene conversion (smaller SLF complement or loss of SI), but these are assumed to be so deleterious that they are instantly eliminated.
# Haplotypes with the new RNase acquire the ability to detoxify its immediate ancestor with the same gene conversion probability.

# Besides gene conversion, the other process occurring is frequency-dependent selection.
# Pollen competition occurs between haplotypes.
# Haplotypes that can detoxify more individuals have an advantage over those that can detoxify fewer.
# Pollen is assumed to be produced and distributed in infinite quantities, such that all ovules are fertilized.
# The exception is that individuals incapable of being detoxified by any pollen haplotype are completely ovule-sterile.
# All individuals have equal viability.

# The population size is held constant at some chosen level.
# For each individual that needs to be generated each generation, a seed parent is chosen at random.
# Then, a pollen grain is chosen at random out of the subset of the pollen pool capable of detoxifying both the seed parent's RNases.
# Each haplotype in the pollen pool occurs at a frequency equal to that haplotype's frequency in the parental generation.

# The simulation stops when every haplotype can detoxify every haplotype but itself, or when a specified maximum generation is reached.
# The difference between the final and initial number of RNases is recorded (max = +1).
# This simulation is repeated a number of times to generate a distribution of results.
# The goal is to determine the probability of increasing through this process.

##########
# LEGEND #
##########

# Each diploid genotype is represented as a 2x3 matrix.
# Each row represents a parental haplotype.
# Ignoring the nonfunctional SLF paralog associated with each RNase, there are three polymorphic loci.
# The first column represents the RNase locus.
# The second column represents the SLF that detoxifies the RNase ancestral to the new one.
# The third column represents the SLF that detoxifies the new RNase.
# The first column contains the letter "A" or "D", or a positive integer.
# "A" represents the ancestor of the new RNase; "D" represents the new RNase.
# All other RNases are represented by positive integers.
# The second and third columns contain either a "-" or a "+".
# "+" represents detoxification ability; "-" represents the absence of detoxification ability.
# It would be possible to use 0 and 1, but "-" and "+" visually distinguish the second and third from the first column.

# IMPORTANT NOTE ABOUT GENE CONVERSION RATES!
# convA and convD are the probabilities that a gamete capable of acquiring a new SLF without becoming SC does so.

##############
# PROPERTIES #
##############

# I will write several functions that identify an object as having a certain list of properties.
# These will mainly be used in assertions.
# This is a substitute for classes.

# A genotype is a 2x3 matrix of strings.
# Its first column is named "S" and contains the letters "A" and "D", as well as positive integers (still character data type).
# Its second column is named "A" and contains the characters "-" and "+".
# Its third column is named "D" and contains the characters "-" and "+".
# Its two row names are identical and represent the individual's number in the population.
# The row names are strings "IndX", where "X" is a placeholder for a positive integer.

# A population is a matrix an nx3 matrix with an even number of rows.
# Its first column is named "S" and contains the letters "A" and "D", as well as positive integers (still character data type).
# Its second column is named "A" and contains the characters "-" and "+".
# Its third column is named "D" and contains the characters "-" and "+".
# The row names are strings "IndX", where "X" is a placeholder for a positive integer.
# Each row name is repeated once, and the two instances of that row name are contiguous.

# A fraction is a positive scalar between 0 and 1.

is.fraction <- function(fraction){
  return(is.vector(fraction) &
           length(fraction) == 1 &
           fraction >= 0 &
           fraction <= 1
  )
}

# Diploid genomes all use the same colnames.

loci.names <- c('S','A','D')

#####################
# FUNCTION JUNCTION #
#####################

# The function is.compatible will tell you whether a pollen genotype can fertilize an ovule parent.

is.compatible <- function(ovule.parent, pollen){
  must.detoxify <- ovule.parent[,'S']
  can.detoxify <- length(which(pollen[must.detoxify] == '+'))
  return(can.detoxify == 2)
}

# The function fertilize will fertilize an ovule parent with compatible pollen.
# It will choose randomly among compatible pollen, weighted by their frequencies.
# This includes recombinant pollen.
# Once it has chosen compatible pollen, the ovule parent will pick an ovule at random.
# This ovule may or may not be recombinant.
# It also takes into account the self-POLLINATION rate (s) and the inbreeding depression (delta).
# NOTE: this model assumes prior selfing.
# Very rare SC pollen genotypes might successfully fertilize self ovules if the self-pollination rate is high.
# This would be a huge advantage for SC haplotypes if inbreeding depression is low.
# It would be a huge disadvantage for SC haplotypes if inbreeding depression is high because it crowds out good outcross seeds with garbage selfed seeds.
# The function could be rewritten to accomodate competing selfing by instead mixing self and outcross pollen together in some proportion.
# However, I don't know how I would 

fertilize <- function(ovule.parent, pollen.pool, delta, conv, s){
  # First, generate a self-pollen pool to see if some self-pollen is self-compatible.
  ovule.parent.as.pop <- ovule.parent
  rownames(ovule.parent.as.pop) <- c(1,1)
  # Create the self pollen pool, which is actually the whole self gamete pool.
  self.pollen.pool <- generate.pollen.pool(ovule.parent.as.pop, conv)
  # Subset both self and outcross pollen pools to contain only the genotypes that actually occur.
  self.pollen.pool <- self.pollen.pool[which(self.pollen.pool[,'p'] > 0),]
  pollen.pool <- pollen.pool[which(pollen.pool[,'p'] > 0),]
  # If only one pollen genotype can be produced, self.pollen.pool will be a vector.
  # In that case, we'll need to coerce it into a 1xn matrix.
  if(is.null(dim(self.pollen.pool))){self.pollen.pool <- t(as.matrix(self.pollen.pool))}
  self.pollen.genotypes <- split(self.pollen.pool,(1:nrow(self.pollen.pool)))
  # Give names.
  self.pollen.genotypes <- lapply(self.pollen.genotypes, name.names, nom = colnames(ovule.parent))
  # Subset only the compatible self pollen.
  compatible.self.pollen <- as.matrix(self.pollen.pool[unlist(lapply(self.pollen.genotypes,
                                                                is.compatible,
                                                                ovule.parent = ovule.parent)
                                                         ),
                                                  ]
                                      )
  # If compatible.self.pollen was originally a vector, it needs to be transposed into a row vector.
  if(ncol(compatible.self.pollen) == 1){compatible.self.pollen <- t(compatible.self.pollen)}
  # Flag whether the ovule parent can self.
  can.self <- nrow(compatible.self.pollen) > 0
  # Now identify which of the outcross pollen genotypes are compatible.
  outcross.pollen.genotypes <- split(pollen.pool,1:nrow(pollen.pool))
  # Give names.
  outcross.pollen.genotypes <- lapply(outcross.pollen.genotypes, name.names, nom = colnames(ovule.parent))
  # Subset only the compatible outcross pollen.
  compatible.outcross.pollen <- as.matrix(pollen.pool[unlist(lapply(outcross.pollen.genotypes,
                                                          is.compatible,
                                                          ovule.parent = ovule.parent)
                                                   ),
                                            ]
                                          )
  # If compatible.outcross.pollen was originally a vector, it needs to be transposed into a row vector.
  if(ncol(compatible.outcross.pollen) == 1){compatible.outcross.pollen <- t(as.matrix(compatible.outcross.pollen))}
  # Flag whether the ovule parent can outcross.
  can.outcross <- nrow(compatible.outcross.pollen) > 0
  # Determine the probabilities of selfing and outcrossing.
  # The probability of selfing is s unless either selfing or outcrossing is impossible.
  selfing.prob <- NA
  if(can.self & can.outcross){
    selfing.prob <- s
  }
  # If it can only self, then it is guaranteed to self.
  if(can.self & !can.outcross){
    selfing.prob <- 1
  }
  # If it can only outcross, then it is guaranteed to outcross.
  if(!can.self & can.outcross){
    selfing.prob <- 0
  }
  # If it can't self nor outcross, then it can't be fertilized.
  if(!can.self & !can.outcross){return('ovule sterile')}
  # Determine if it actually selfs.
  does.self <- rbinom(n = 1, size = 1, prob = selfing.prob) > 0
  # Initialize output.
  out <- matrix(nrow = 2, ncol = ncol(ovule.parent))
  colnames(out) <- colnames(ovule.parent)
  # If it selfs,
  if(does.self){
    # Determine if inbred offspring die from inbreeding depression.
    does.die <- rbinom(1, 1, delta) > 0
    if(does.die){return('dead')}
    # Pick a pollen grain from the possible self pollen genotypes weighted by its frequency.
    parental.pollen <- compatible.self.pollen[sample(1:nrow(compatible.self.pollen),
                                                            size = 1,
                                                            prob = compatible.self.pollen[,'p']
                                                     ),
                                              ]
    # The function generate.pollen.pool should work for ovules, too.
    # I guess I should have called it generate.gamete.pool.
    # However, the ovule genotype is not required to be one of the SC genotypes.
    parental.ovule <- self.pollen.pool[sample(1:nrow(self.pollen.pool),
                                                           size = 1,                                              
                                                           prob = self.pollen.pool[,'p']
                                                    ),
                                             ]
    out[1,] <- parental.pollen[names(parental.pollen)!='p']
    out[2,] <- parental.ovule[names(parental.ovule)!='p']
  }
  # If it outcrosses,
  if(!does.self){
    # Pick a pollen grain from the possible outcross pollen genotypes weighted by its frequency.
    parental.pollen <- compatible.outcross.pollen[sample(1:nrow(compatible.outcross.pollen),
                                                            size = 1,
                                                            prob = compatible.outcross.pollen[,'p']
                                                     ),
                                              ]
    # The function generate.pollen.pool should work for ovules, too.
    # I guess I should have called it generate.gamete.pool.
    # However, the ovule genotype is not required to be one of the SC genotypes.
    parental.ovule <- self.pollen.pool[sample(1:nrow(self.pollen.pool),
                                                     size = 1,
                                                     prob = self.pollen.pool[,'p']
                                              ),
                                       ]
    out[1,] <- parental.pollen[names(parental.pollen)!='p']
    out[2,] <- parental.ovule[names(parental.ovule)!='p']
  }
  return(out)
}


# The function advance.generation will repeat mate.individual on random individuals until the population is regenerated.

advance.generation <- function(population, delta, conv, s){
  # First, determine the population size.
  pop.size <- nrow(population)/2
  # Initialize the output.
  out <- matrix(nrow = nrow(population), ncol = ncol(population))
  colnames(out) <- colnames(population)
  rownames(out) <- rep(1:pop.size, times = 1, each = 2)
  # Create a new individual for every one that exists currently.
  i <- 1
  pollen.pool <- generate.pollen.pool(population, conv)
  while(i <= pop.size){
    # Pick a random ovule.parent.id
    ovule.parent.id <- as.character(sample(1:pop.size,1))
    ovule.parent <- population[which(rownames(population) == ovule.parent.id),]
    # Mate it.
    mating.outcome <- fertilize(ovule.parent, pollen.pool, delta, conv, s)
    # If the ovule parent is ovule sterile or produces a dead offspring, that ovule fails to contribute to the next generation.
    # The outputs "ovule sterile" and "dead" aren't matrices, so they will result in trying again to create the offspring.
    # Any output that is a matrix will be included.
    if(is.matrix(mating.outcome)){
      out[rownames(out) == i,] <- mating.outcome
      i <- i + 1
    }
  }
  return(out)
}

# The function simulate.slf.spread will repeat advance generation for a desired number of generations.

simulate.slf.spread <- function(population, delta, conv, s, gens){
  for(i in 1:gens){
    # Update the population with advance.generation.
    population <- advance.generation(population, delta, conv, s)
  }
  return(population)
}

# The function generate.haplotype will create a vector representing an S-haplotype.
# It needs to know how many RNases there are and which one this is.
# The argument kind tells the function whether the haplotype is adapted, not adapted, ancestral, or derived.

generate.haplotype <- function(numHaplotypes, thisHaplotype, kind){
  # First, intialize a vector of the appropriate length.
  out <- vector(length = numHaplotypes + 1)
  # Name the elements.
  names(out) <- c('S', 'A', 'D', as.character(1:(numHaplotypes-2)))
  # The RNase is specified by thisHaplotype.
  out['S'] <- as.character(thisHaplotype)
  # The SLF genotypes depend on what kind of haplotype we're given.
  if(kind == 'notAdapted'){
    # A normal haplotype can detoxify everything but itself and the derived RNase.
    out['A'] <- '+'
    out['D'] <- '-'
    out[as.character(1:(numHaplotypes-2))] <- '+'
    out[as.character(thisHaplotype)] <- '-'
  }
  if(kind == 'Adapted'){
    # An adapted haplotype can detoxify the derived RNase.
    out['A'] <- '+'
    out['D'] <- '+'
    out[as.character(1:(numHaplotypes-2))] <- '+'
    out[as.character(thisHaplotype)] <- '-'
  }
  if(kind == 'ancestral'){
    out['A'] <- '-'
    out['D'] <- '-'
    out[as.character(1:(numHaplotypes-2))] <- '+'
  }
  if(kind == 'adaptedAncestral'){
    out['A'] <- '-'
    out['D'] <- '+'
    out[as.character(1:(numHaplotypes-2))] <- '+'
  }
  if(kind == 'derived'){
    out['A'] <- '-'
    out['D'] <- '-'
    out[as.character(1:(numHaplotypes-2))] <- '+'
  }
  if(kind == 'adaptedDerived'){
    out['A'] <- '+'
    out['D'] <- '-'
    out[as.character(1:(numHaplotypes-2))] <- '+'
  }
  return(out)
}

# Since it's a pain to generate population by hand, the function generate.pop will make one for you.
# It takes as arguments several vectors of haplotype frequencies.
# The vector notAdapted represents the frequencies of all haplotypes that have not acquired the ability to detoxify the new RNase.
# The vector adapted represents the frequencies of haplotypes that have acquired the ability to detoxify the new RNase.
# Haplotypes ancestral to the adapted haplotypes are assumed to have gone extinct.
# The scalar ancestral represents the frequency of the haplotype ancestral to the new RNase.
# The scalar derived represents the frequency of the new RNase.
# The argument AorD tells you whether the ancestral or the derived RNase has acquired the ability to detoxify the other.
# AorD can take the take the values 'A' (ancestral), 'D' (derived), or 'N' (neither).
# You must also specify the population size.

# generate.pop <- function(pop.size, notAdapted, adapted, ancestral, derived, AorD){
#   # First create a matrix of the appropriate size to hold the population.
#   # There will be 2*pop.size rows because individuals are diploid.
#   # There will one column the RNase and one column for each SLF.
#   numSLF <- length(notAdapted) + length(adapted) + 2
#   out <- matrix(nrow = 2*pop.size, ncol = numSLF + 1)
#   # Each individual should get two rows.
#   rownames(out) <- rep(1:pop.size, times = 1, each = 2)
#   # One column for the RNase and one for each SLF.
#   colnames(out) <- c('S', 'A', 'D', as.character(1:(numSLF-2)))
#   # Now create each kind of haplotype based on the information known.
#   
# }

# The function pollen.probability tells you the probability of generating pollen of a given genotype given a parental genotype and a gene conversion probability.
pollen.probability <- function(parent, pollen, conv){
  # The pollen genotype should be as long as the parental genome.
  stopifnot(length(pollen) == ncol(parent)
  )
  pollen <- lapply(pollen, name.names, nom = names(parent))
  # Count the SLF loci.
  num.slf <- ncol(parent) - 1
  # The procedure starts by picking the base chromosome.
  # If the parent is heterozygous for the RNase, the base chromosome is unambiguously the one with the same RNase allele as the pollen.
  base.id <- pollen['S'] == parent[,'S']
  # If neither parent has the RNase, the probability is zero because we aren't modeling RNase mutation.
  if(length(which(base.id))==0){return(0)}
  # If only one parent has it, the base parent is unambiguous.
  if(length(which(base.id))==1){
    base.chrom <- parent[base.id,]
    base.chrom.slfs <- base.chrom[names(base.chrom) != 'S']
    other.chrom <- parent[!base.id,]
    other.chrom.slfs <- other.chrom[names(other.chrom) != 'S']
    # Initialize a vector of locus probabilities.
    loc.probs <- vector(length = ncol(parent))
    names(loc.probs) <- names(pollen)
    # Then, we calculate the probability of getting the observed RNase.
    # If neither chromosome has it, the probability is zero.
    if(pollen['S'] != base.chrom['S'] & pollen['S'] != other.chrom['S']){loc.probs['S'] <- 0}
    # If both chromosomes have it, the probability is one (this can only occur after SC haplotypes are created by gene conversion).
    if(pollen['S'] == base.chrom['S'] & pollen['S'] == other.chrom['S']){loc.probs['S'] <- 1}
    # If only the base chromosome has it, the probability is one half (the probability of drawing that base chromosome).
    if(pollen['S'] == base.chrom['S'] & pollen['S'] != other.chrom['S']){loc.probs['S'] <- 0.5}
    # If only the other chromosome has it, the probability is zero (RNase gene conversion and mutation are not modeled).
    if(pollen['S'] != base.chrom['S'] & pollen['S'] == other.chrom['S']){loc.probs['S'] <- 0}
    # Now we find loci at which the SLF equals the base, the other chromosome, both, or neither
    slf.locs <- pollen[names(pollen) != 'S']
    slf.loc.probs <- vector(length = length(slf.locs))
    names(slf.loc.probs) <- names(slf.locs)
    same.as.base.only <- slf.locs == base.chrom.slfs & slf.locs != other.chrom.slfs
    same.as.other.only <- slf.locs != base.chrom.slfs & slf.locs == other.chrom.slfs
    same.as.both <- slf.locs == base.chrom.slfs & slf.locs == other.chrom.slfs
    same.as.neither <- slf.locs != base.chrom.slfs & slf.locs != other.chrom.slfs
    # Then, at each SLF locus, we assign a probability of getting the observed pollen genotype.
    # If neither chromosome has the pollen's genotype, the probability is zero since we aren't modeling mutation.
    slf.loc.probs[same.as.neither] <- 0
    # If both chromosomes have the pollen's genotype, the probability is one.
    slf.loc.probs[same.as.both] <- 1
    # If the base chromosome has it, the probability is the complement of the gene conversion probability.
    slf.loc.probs[same.as.base.only] <- 1 - conv
    # If the other chromosome has it, the probability is the gene conversion probability.
    slf.loc.probs[same.as.other.only] <- conv
    # Assuming the probability of gene conversion at each locus is independent, the probability of the observed pollen genotype is the product of the locus probabilities.
    loc.probs[names(loc.probs) != 'S'] <- slf.loc.probs
    return(prod(loc.probs))
  }
  # The most complicated possibility occurs when the RNase could have come from either chromosome.
  if(length(which(base.id)) == 2){
    # First we calculate the probability of the pollen genotype given that one chromosome is the base.
    # Then we repeat this calculation assuming the other is the base.
    possible.probs <- vector(length = 2)
    for(i in 1:2){
      base.chrom <- parent[c(1,2)[i],]
      base.chrom.slfs <- base.chrom[names(base.chrom) != 'S']
      other.chrom <- parent[c(2,1)[i],]
      other.chrom.slfs <- other.chrom[names(other.chrom) != 'S']
      # Initialize a vector of locus probabilities.
      loc.probs <- vector(length = ncol(parent))
      names(loc.probs) <- names(pollen)
      # Then, we calculate the probability of getting the observed RNase.
      # If neither chromosome has it, the probability is zero.
      if(pollen['S'] != base.chrom['S'] & pollen['S'] != other.chrom['S']){loc.probs['S'] <- 0}
      # If both chromosomes have it, the probability is one (this can only occur after SC haplotypes are created by gene conversion).
      if(pollen['S'] == base.chrom['S'] & pollen['S'] == other.chrom['S']){loc.probs['S'] <- 1}
      # If only the base chromosome has it, the probability is one half (the probability of drawing that base chromosome).
      if(pollen['S'] == base.chrom['S'] & pollen['S'] != other.chrom['S']){loc.probs['S'] <- 0.5}
      # If only the other chromosome has it, the probability is zero (RNase gene conversion and mutation are not modeled).
      if(pollen['S'] != base.chrom['S'] & pollen['S'] == other.chrom['S']){loc.probs['S'] <- 0}
      # Now we find loci at which the SLF equals the base, the other chromosome, both, or neither
      slf.locs <- pollen[names(pollen) != 'S']
      slf.loc.probs <- vector(length = length(slf.locs))
      names(slf.loc.probs) <- names(slf.locs)
      same.as.base.only <- slf.locs == base.chrom.slfs & slf.locs != other.chrom.slfs
      same.as.other.only <- slf.locs != base.chrom.slfs & slf.locs == other.chrom.slfs
      same.as.both <- slf.locs == base.chrom.slfs & slf.locs == other.chrom.slfs
      same.as.neither <- slf.locs != base.chrom.slfs & slf.locs != other.chrom.slfs
      # Then, at each SLF locus, we assign a probability of getting the observed pollen genotype.
      # If neither chromosome has the pollen's genotype, the probability is zero since we aren't modeling mutation.
      slf.loc.probs[same.as.neither] <- 0
      # If both chromosomes have the pollen's genotype, the probability is one.
      slf.loc.probs[same.as.both] <- 1
      # If the base chromosome has it, the probability is the complement of the gene conversion probability.
      slf.loc.probs[same.as.base.only] <- 1 - conv
      # If the other chromosome has it, the probability is the gene conversion probability.
      slf.loc.probs[same.as.other.only] <- conv
      # Assuming the probability of gene conversion at each locus is independent, the probability of the observed pollen genotype is the product of the locus probabilities.
      loc.probs[names(loc.probs) != 'S'] <- slf.loc.probs
      possible.probs[i] <- prod(loc.probs)
    }
    # Now add the probabilities of getting this genotype given that either parental chromosome could have been the base.
    # Note that the elements of possible.probs are not the probabilities conditional on receiving those parental probabilities.
    # I have already multiplied by half.
    return(sum(possible.probs))
  }
}

# The function name.names will take a vector or a matrix and give it names.
# I want to use it to give all the vectors or matrices in a list the same names.

name.names <- function(vec.or.mat, nom){
  out <- vec.or.mat
  # If it's a vector, use names()
  if(is.null(dim(vec.or.mat))){names(out) <- nom}
  # If it's a matrix, use colnames()
  else{colnames(out) <- nom}
  return(out)
}

# The function individual.pollen.output tells you the frequency of each pollen genotype produced by an individual.
# It requires the parent, a rate of gene conversion, and a matrix of all possible pollen genotypes.
individual.pollen.output <- function(parent, conv, possible.genos){
  # We have all the pollen genotypes.
  # We just need to calculate their frequencies.
  # This will be done by calling pollen.probability.
  # We'll put all the pollen genotypes into a list.
  pollen.list <-  split(t(possible.genos),
                        rep(1:ncol(t(possible.genos)),
                            each = nrow(t(possible.genos)
                                        )
                            )
                        )
  pollen.list <- lapply(pollen.list, FUN = name.names, nom = colnames(parent))
  # And then we'll use lapply to get a vector of pollen frequencies.
  # The output matrix will be the same size as possible.genos, except that it has an extra column for frequency.
  out <- cbind(possible.genos, rep(0, times = nrow(possible.genos)))
  colnames(out) <- c(colnames(possible.genos),'p')
  out[,'p'] <- unlist(lapply(pollen.list, FUN = pollen.probability, parent = parent, conv = conv))
  return(out)
}

# The  function all.pollen generates every pollen genotype for genomes of the specified size, regardless of whether any individual can make the pollen.

all.pollen <- function(n.rnase){
  # First, let's permute the pollen genotypes.
  pollen.perms <- permutations(n = 2,
                               r = n.rnase,
                               v = c('-','+'),
                               repeats.allowed = TRUE)
  # Now that we have pollen.perms, we know how big the output matrix will be.
  # It will be the number of RNases times the number of pollen permutations.
  # There will be one column for every SLF (including for the ancestral and derived RNases), and one row for the RNase locus.
  out <- matrix(nrow = (n.rnase * nrow(pollen.perms)), ncol = n.rnase+1)
  # Now we've got to name our columns.
  colnames(out) <- c('S','A','D',as.character(1:(n.rnase-2)))
  slf.options <- colnames(out)[2:length(colnames(out))]
  # The first column will consist of repeats of each RNase equal to the number of pollen permutations.
  out[,'S'] <- rep(slf.options, times = 1, each = nrow(pollen.perms))
  # The matrix will consist of the pollen.perms repeated once for every RNase.
  for(i in 1:n.rnase){
    repeated.slf <- rep(slf.options[i], times = nrow(pollen.perms))
    out[(1+(i-1)*nrow(pollen.perms)):(i*nrow(pollen.perms)),] <- cbind(repeated.slf,
                                                                      pollen.perms)
  }
  return(out)
}

# The function get.p will grab the column from a matrix with the column name 'p'

get.p <- function(mat){
  return(as.numeric(mat[,'p']))
}

# The function generate.pollen.pool will generate an infinite pollen pool with pollen genotypes proportional to the probability of creating them in the population.
# It requires a population and a gene conversion rate.
# It will have to be called once every generation because the population will change, but that's still better than being called once per individual.

generate.pollen.pool <- function(population, conv){
  # First, generate all pollen genotypes.
  all.genos <- all.pollen(ncol(population)-1)
  # We will now apply individual.pollen.output to every individual in the population.
  # But first we need to split the population into individuals.
  individuals <- lapply(split(population,rownames(population)), matrix, nrow = 2, ncol = ncol(population))
  individuals <- lapply(individuals, name.names, nom = colnames(population))
  # Now for each individual, use individual.pollen.output.
  pollen.outputs <- lapply(individuals, individual.pollen.output, possible.genos = all.genos, conv = conv)
  just.p <- lapply(pollen.outputs, get.p)
  out <- cbind(all.genos,Reduce('+',just.p)/(nrow(population)/2))
  colnames(out) <- c(colnames(population),'p')
  # Now that we have a list of matrices of pollen outputs, we need to add them together.
  return(out)
}

# The function construct.individual creates a diploid genotype with minimal hassle.
# It takes as an argument the number of RNase alleles in the population, and assumes that there's one SLF paralog for every RNase allele.
# It assumes that each haplotype can detoxify everything except itself and the derived RNase.
# It needs a diploid RNase genotype.
# n.rnases includes the ancestral and derived rnases.

construct.individual <- function(n.rnases, hap1, hap2){
  # Initialize the individual, an 2x(n.rnases+1) matrix because there is one RNase locus and n.rnases SLF loci.
  individual <- matrix(nrow = 2, ncol = n.rnases+1)
  # Columns are loci.
  colnames(individual) <- c('S','A','D',1:(n.rnases-2))
  # The RNase haplotypes present give the diploid RNase genotype.
  individual[,1] <- c(hap1,hap2)
  # It can detoxify everything except itself and D.
  individual[1,which(colnames(individual) != 'S')] <- '+'
  individual[1,which(colnames(individual) == 'D')] <- '-'
  individual[1,which(colnames(individual) == individual[1,1])] <- '-'
  individual[2,which(colnames(individual) != 'S')] <- '+'
  individual[2,which(colnames(individual) == 'D')] <- '-'
  individual[2,which(colnames(individual) == individual[2,1])] <- '-'
  return(individual)
}

####################################
### DEPRECATED FUNCTION DUMPSTER ###
####################################

### DEPRECATED ###
# The function mate.individual will keep trying to make an offspring through selfing and outcrossing with the appropriate probabilties.
# If outcross.individual or self.individual returns 'incompatible', it will try again.
# If self.individual returns 'dead', it will stop trying and let a different ovule parent mate.
# If outcross.individual or self.individual returns an offspring matrix, it will output this.
# s is the selfing rate.
# 
# mate.individual <- function(population, ovule.parent.id, delta, conv, s){
#   # Keep trying for some number of attempts.
#   attempts <- 0
#   tolerance <- 1000
#   while(attempts < tolerance){
#     # Determine if the individual should self.
#     self <- rbinom(1, 1, prob = s) == 1
#     if(self){
#       selfing.parent <- population[which(row.names(population)==ovule.parent.id),]
#       selfing.outcome <- self.individual(selfing.parent, delta, conv)
#       # If the pollen was incompatible, just remain in the loop.
#       if(identical(selfing.outcome, 'incompatible')){}
#       # Otherwise, return the value of selfing.outcome, which may be an offspring matrix or the string 'dead'.
#       else{return(selfing.outcome)}
#     }
#     # If self is not true, outcross instead.
#     else{
#       outcross.result <- outcross.individual(population, ovule.parent.id, conv)
#       # Just stay in the loop if the pollen is incompatible.
#       if(identical(outcross.result, 'incompatible')){}
#       # Otherwise, return the value of outcross.result.
#       else{return(outcross.result)}
#     }
#     # If mating failed this time, then add one to attempts.
#     attempts <- attempts + 1
#   }
#   # If all allowed attempts have been made, then return 'failed'.
#   if(attempts >= tolerance){return('failed')}
# }

# The function gamete.from.individual takes one individual and produces one gamete from it.
# This gamete can be either parental or recombinant.
# Recombinant gaametes are only produced by gene conversion.
# Gene conversion happens at rate conv.

# gamete.from.individual <- function(parent, conv){
#   # First, pick one parental gamete to be the base.
#   winner <- sample(c(1,2),1)
#   if(winner == 1){loser <- 2}
#   if(winner == 2){loser <- 1}
#   base <- parent[winner,]
#   # O.K., now we've got our RNase.
#   out <- vector(length = length(base))
#   names(out) <- names(base)
#   out['S'] <- base['S']
#   # Distinguish the RNase site from the SLF sites.
#   slf.sites <- which(colnames(parent) != 'S')
#   # Identify heterozygous SLF loci.
#   het.sites <- which(parent[1,slf.sites] != parent[2,slf.sites])
#   # Roll the dice for each site to convert.
#   converted <- vector(length = length(slf.sites))
#   converted[het.sites] <- rbinom(n = length(het.sites), size = 1, prob = conv) > 0
#   # Update out with the gene conversions.
#   out[slf.sites][which(converted)] <- parent[loser,][slf.sites][which(converted)]
#   out[slf.sites][which(!converted)] <- parent[winner,][slf.sites][which(!converted)]
#   return(out)
# }

# The function outcross.individual will take as arguments a population and the ID of an ovule parent.
# It will only attempt outcrossing once.
# If it fails, it will return the string 'incompatible'.

# outcross.individual <- function(population, ovule.parent.id, conv){
#   # Grab the ovule parent.
#   ovule.parent <- population[which(row.names(population) == ovule.parent.id),]
#   # Initialize the offspring.
#   offspring <- matrix(nrow = nrow(ovule.parent), ncol = ncol(ovule.parent))
#   # Pick a pollen parent at random from the population.
#   pollen.parent.id <- sample(1:(nrow(population)/2), 1)
#   pollen.parent <- population[which(row.names(population) == pollen.parent.id),]
#   pollen.grain <- gamete.from.individual(pollen.parent, conv)
#   # Check that the pollen grain can detoxify both of the ovule parent's RNases.
#   if(pollen.grain[ovule.parent[1,'S']] == '+' & pollen.grain[ovule.parent[2,'S']] == '+'){
#     offspring[1,] <- pollen.grain
#     offspring[2,] <- gamete.from.individual(ovule.parent, conv)
#     return(offspring)
#   }
#   else{return('incompatible')}
# }

# The function self.individual works like outcross.individual, but the pollen parent is the ovule parent,
# and the offspring only survives with probability 1 - delta, where delta is the inbreeding depression.

# self.individual <- function(ovule.parent, delta, conv){
#   # Determine if fertilization is successful.
#   offspring <- matrix(nrow = nrow(ovule.parent), ncol = ncol(ovule.parent))
#   pollen.grain <- gamete.from.individual(ovule.parent, conv)
#   if(pollen.grain[ovule.parent[1,'S']] == '+' & pollen.grain[ovule.parent[2,'S']] == '+'){
#     # Now that fertilization has occurred,
#     # determine if the offspring survives.
#     survives <- rbinom(n = 1, size = 1, prob = (1 - delta))
#     if(!survives){return('dead')}
#     else{
#       offspring[1,] <- pollen.grain
#       offspring[2,] <- gamete.from.individual(ovule.parent, conv)
#       return(offspring)
#     }
#   }
#   else{return('incompatible')}
# }
```
