## Supplementary material for "Diversification or collapse of self-incompatibility haplotypes as a rescue process": All scripts: convert_nb.R

```
eq <- read.table('../data/equilibria.txt', stringsAsFactors = F)
names <- eq[1,]
names <- do.call(rbind,strsplit(names, ','))
names <- gsub('\\{', '', names)
names <- gsub('\\}', '', names)
eq <- do.call(rbind,strsplit(eq[-1,1], "\\}\\},"))

sols <- eq[,1]
params <- eq[,2]

sols <- 
  
  sapply(strsplit(sols, '\\},\\{'),function(X){
    as.numeric(gsub("p->","",ifelse(length(X==3),X[2],NA)))
    })

params <- gsub('\\}', '', params)
params <- do.call(rbind,strsplit(params, ","))
params <- matrix(as.numeric(params), ncol = 3)

eq <- cbind(sols, params)
colnames(eq) <- names[1,]

#slice <- eq[eq[,2]==0.0001 & eq[,3]==0.1,]
#plot(x = slice[,4], y = slice[,1])
#par(mfrow = c(2,2)
plot(c(NA,NA), xlim = c(7,40), y = c(0,0.06), xlab = 'RNases', ylab = 'Equilibrium frequency',
     cex.axis = 2,
     cex.lab = 2)
library(RColorBrewer)
palette <- brewer.pal(10, 'RdBu')
for(i in 2:10){
  slice <- eq[eq[,2]==0.0003 & eq[,3]==unique(eq[,3])[i],]
  points(x = slice[,4], y= slice[,1], xlab = 'RNases', ylab = 'Equilibrium frequency', type = 'l', col = palette[i])
}
legend('topright', fill = palette[2:10], legend = unique(eq[,3])[2:10], ncol = 2, title = 'Selfing rate',
       cex=0.9,
       text.width=2)
```
