## Supplementary material for "Diversification or collapse of self-incompatibility haplotypes as a rescue process": All scripts: distribution_plot.R

```
# Pick an R value.
R <- 0.01


overR <- function(R){
# Load a transition matrix.
transition <- as.matrix(read.csv(paste('../data/transition_L=2_R=', R, '.csv', sep = ''), check.names = F, row.names = 1))

# The transition probabilities out of the first row were not previously calculated.
# If there are no unluckies, then it is impossible to produce more haplotypes.
# The first row is therefore an absorbing state.
transition[1,] <- c(1, rep(0, (ncol(transition)-1)))
# The exception might be if the first row is two haplotypes.
# In this case, the assumption of zero pollen limitation must logically break down because no one can fertilize anyone else.
# It might therefore make sense for it to be a reflecting boundary.
if('2' %in% colnames(transition)){
  # If there exists a state with only two haplotypes, make it impossible to transition to it.
  # All density that would transition into 2 is instead reflected to 3.
  transition[,'3'] <- transition[,'3'] + transition[,'2']
  transition[,'2'] <- 0
}

# Assuming we start with one unlucky, see what happens when we iterate forward.
starting_num <- 3
initial <- c(rep(0,(starting_num-1)), 1, rep(0, ncol(transition)-starting_num))
# Assume instead a uniform initial distribution.
#initial <- rep(1/ncol(transition), ncol(transition))
names(initial) <- colnames(transition)

# Now iterate.
generations <- 1000
current <- initial
for(i in 1:generations){
  current <- t(transition) %*% current
  # Try to restore rounding errors. This seems like a weird step, and I think it's probably wrong.
  #current <- current/(sum(current))
}
return(cbind(R=R, n =2:42, prob = current))
}
library(tidyverse)
library(ggthemes)
data.frame(do.call(rbind,lapply(c(.01,.1,.2,.3,1), overR))) %>%
  ggplot(aes(x = n, y = V3, fill = factor(R)))+
  geom_density(stat = "identity", alpha = .5,n = 7^2, show.legend = FALSE)+
  theme_light()+
  annotate(geom = "text",x = c(5,9,12,38,35), y = c(.9,.3,.2,.2,.9), label =  paste("R =", c(0.01,.1,.2,.3,1)))

outpath <- paste('../figures/stable_R=', R, '.pdf', sep = '')

pdf(outpath)
par(mar = c(8,9,4,2), mgp = c(6, 2, 0), cex.axis = 4, cex.lab = 4, lwd = 2)
plot(current,
     x = as.numeric(rownames(current)),
     xlab = 'Total haplotypes',
     ylab = 'Frequency',
     type = 'h',
     #main = paste('R=', R, sep = '')
     main = NULL
)
dev.off()
```
