## Supplementary material for "Diversification or collapse of self-incompatibility haplotypes as a rescue process": All scripts: equilibration.R

```
# This script finds the equilibrium genotype frequencies after the introduction of an RNase mutation.

# Load the functions.

source('scripts/rnase_functions.R')

# Initialize the population.

my_gens <- 1000 # The number of generations you want to track.
L <- 2 # The number of haplotypes that have a variant with the new key.
U <- 2 # The number of haplotypes that have no variant with the new key.
luck_frac <- 0.01 # The fraction of each of the L haplotypes that actually has the new key.
spite_frac <- 0.01 # The fraction of the lock-mutant's ancestor that actually mutated to the lock-mutant.

initialize(L,
           U,
           luck_frac,
           spite_frac)

# Create trajectories of diploid genotype frequencies.
# Plot trajectories of haplotype frequencies.

traj <- track.diplos(population, my_gens)
al.traj <- matrix(nrow = nrow(traj), ncol = 5)
colnames(al.traj) <- c('L','C','U','A','S')
al.traj[,'L'] <- traj[,'LL'] + traj[,'LC']/2 + traj[,'LU']/2 + traj[,'LA']/2 + traj[,'LS']/2
al.traj[,'C'] <- traj[,'LC']/2 + traj[,'CC'] + traj[,'CU']/2 + traj[,'CA']/2 + traj[,'CS']/2
al.traj[,'U'] <- traj[,'LU']/2 + traj[,'CU']/2 + traj[,'UU'] + traj[,'UA']/2 + traj[,'US']/2
al.traj[,'A'] <- traj[,'LA']/2 + traj[,'CA']/2 + traj[,'UA']/2
al.traj[,'S'] <- traj[,'LS']/2 + traj[,'CS']/2 + traj[,'US']/2

traj.for.fit <- split(traj, 1:nrow(traj))
named.traj.for.fit <- lapply(traj.for.fit, rename, nom = colnames(traj))
fit.traj <- unlist(lapply(named.traj.for.fit, get.mean.pollen.fitness))

plot(al.traj[,'L'], col = 1, type = 'l', xlab = 'Generation', ylab = 'Frequency', ylim = c(0,1))
points(al.traj[,'C'], col = 2, type = 'l')
points(al.traj[,'U'], col = 3, type = 'l')
points(al.traj[,'A'], col = 4, type = 'l')
points(al.traj[,'S'], col = 5, type = 'l')
points(fit.traj, col = 6, type = 'l')
legend(x = 'topleft', fill = c(1,2,3,4,5,6), legend = c('L','C','U','A','S','W'))
```
