## Supplementary material for "Diversification or collapse of self-incompatibility haplotypes as a rescue process": All scripts: explicit_haplotyes.R

```
library(gtools)
library(plyr)
# This is another attempt to get the recursions right on my S-RNase idea.
# For some reason, the diploid genotype recursions I programmed into equilibrate_ancestral.R do not exactly
# recapitulate the predictions I made on paper, though they do have the same requirements for coexistence.

############################
# INITIALIZATION FUNCTIONS #
############################

# The function generate.haplos will create a vector with names for all your haplotypes.
generate.haplos <- function(U, L, A = 1, S = 1){
  # Assertions
  stopifnot(A == 0 | A == 1, # There can be a maximum of one new RNase.
            S == 0 | S == 1)
  haplos <- vector(length = U + L + A + S) # Count up the haplotypes.
  haplos[1:U] <- paste('U', 1:U, sep = '') # Name the us.
  haplos[(U+1):(U+L)] <- paste('L', 1:L, sep = '') # Name the Ls.
  if(A==1){haplos[(U+L+A)] <- 'A'} # Name A, if it exists.
  if(S==1){haplos[(U+L+A+S)] <- 'S'} # Name S, if it exists.
  # Now we have the names of the haplotypes.
  return(haplos)
}

# The function generate.diplos will generate every possible diploid genotype given the different haplotypes.
# The only argument is haplos, which is the output of generate.haplos.
generate.diplos <- function(haplos){
  # The number of genotypes is like n choose 2, except that certain genotypes are impossible.
  # The impossible genotypes are homozygotes (all of them), and AS.
  # First, create a matrix containing all the combinations.
  conceivable_diplos <- combinations(n = length(haplos), r = 2, v = haplos)
  # Now that we have all conceivable diploid genotypes, we need to mark the impossible ones.
  impossible_ids <- conceivable_diplos[,1] == conceivable_diplos[,2] | # The two haplotypes can't be identical.
    conceivable_diplos[,1] == 'A' & conceivable_diplos[,2] == 'S' | # AS is impossible.
    conceivable_diplos[,2] == 'A' & conceivable_diplos[,1] == 'S'   # AS is impossible.
  # Now that we know what's impossible, just subset to what's possible.
  possible_diplos <- conceivable_diplos[!impossible_ids,]
  # Sort alphabetically within each pair.
  for(i in 1:nrow(possible_diplos)){
    possible_diplos[i,] <- sort(possible_diplos[i,])
  }
  return(possible_diplos)
}

# The function is.compatible determines whether a pollen grain is compatible with a particular diploid maternal genotype.
# The argument pollen.haplo should be a single string representing the pollen's S-haplotype.
# The argument maternal.diplo should be a vector of two strings representing the maternal plant's genotype.
is.compatible <- function(pollen.haplo, maternal.diplo){
  # There are a few ways to be incompatible.
  # If none of them are satisfied, then the plants are compatible.
  # By default, set the flag compatible to TRUE.
  compatible <- T
  # The pollen is incompatible if it matches one of the maternal plant's two haplotypes.
  if(sum(maternal.diplo == pollen.haplo) > 0){compatible <- F}
  # The pollen is incompatible if it is unlucky and the maternal plant has an S.
  if(sum(unlist(strsplit(pollen.haplo, split = '')) == 'U') > 0 & sum(maternal.diplo == 'S') > 0){compatible <- F}
  # The pollen is incompatible if it is A and the maternal plant has an S.
  if(pollen.haplo == 'A' & sum(maternal.diplo == 'S') > 0){compatible <- F}
  # The pollen is incompatible if it is S and the maternal plant has an A.
  if(pollen.haplo == 'S' & sum(maternal.diplo == 'A') > 0){compatible <- F}
  return(compatible)
}

# Now that we know the possible genotypes, we can generate a compatibility matrix.
# The function generate.comp.mat will generate a matrix of which pollen haplotypes can fertilize which diploid genotypes.
# Its only argument is the the list of diploid genotypes, from which it can pull the pollen haplotypes.
generate.comp.mat <- function(diplos){
  # First, extract the pollen haplotypes.
  pollen_haplos <- unique(as.vector(diplos))
  # Now create a logical matrix with one row for each diploid genotype and one column for each pollen haplotype.
  # This matrix tells you which pollen each maternal genotype can accept.
  comp_mat <- matrix(nrow = nrow(diplos), ncol = length(pollen_haplos))
  rownames(comp_mat) <- paste(diplos[,1], diplos[,2], sep = '_')
  colnames(comp_mat) <- pollen_haplos
  # Now check each cross to see if it works.
  for(i in 1:ncol(comp_mat)){
    comp_mat[,i] <- unlist(lapply(split(diplos, 1:nrow(diplos)), is.compatible, pollen.haplo = pollen_haplos[i]))
  }
  return(comp_mat)
}

# The function generate.pollen.production.matrix will create a matrix of what diploid genotypes produce what pollen.
generate.pollen.production.matrix <- function(diplos){
  # First, identify all the haplotypes.
  haplos <- unique(as.vector(diplos[,1:2]))
  # These will be the column names of the output matrix.
  out <- matrix(nrow = nrow(diplos), ncol = length(haplos))
  rownames(out) <- paste(diplos[,1], diplos[,2], sep = '_')
  colnames(out) <- haplos
  # Now pull out which diploid genotypes actually produce each pollen type.
  for(i in 1:ncol(out)){
    out[,i] <- haplos[i] == diplos[,1] | haplos[i] == diplos[,2]
  }
  # Replace TRUEs with 0.5 and FALSEs with 0.
  out <- out * 0.5
  return(out)
}

# The function generate.proper.diplos will create a vector in which the names are all diploid permutations,
# not just the biologically possible genotypes.
# The contents of each element will be the proper permutation (one per combination) for that genotype.
generate.proper.diplos <- function(haplos){
  # Create the names.
  gen_perm_mat <- permutations(n = length(haplos), r = 2, v = haplos, repeats.allowed = T)
  gen_perm_vec <- paste(gen_perm_mat[,1], gen_perm_mat[,2], sep = '_')
  # Create a vector of sorted genotypes.
  out <- vector(length = length(gen_perm_vec))
  names(out) <- gen_perm_vec
  for(i in 1:nrow(gen_perm_mat)){
    out[i] <- paste(sort(c(gen_perm_mat[i,1], gen_perm_mat[i,2])), collapse = '_')
  }
  return(out)
}

#############################
# REPEATEDLY USED FUNCTIONS #
#############################

# The function extract.pollen.vector will pull a pollen frequency vector from a vector of diploid genotype frequencies.
extract.pollen.vector <- function(diplo.freqs, pollen.production.matrix){
  # Check to make sure the diploid genotype frequencies are named and arranged in the right order.
  stopifnot(identical(names(diplo.freqs), rownames(pollen.production.matrix)))
  # Just right multiply the pollen.production matrix by the vector of genotype frequencies to get the pollen frequencies.
  out <-  as.vector(diplo.freqs %*% pollen.production.matrix)
  names(out) <- colnames(pollen.production.matrix)
  return(out)
}

# The function generate.sire.matrix multiplies the compatibility matrix by a matrix of repeated pollen frequencies,
# and then divides each row by the sum of each row to get the relative contributions of each pollen
# type to fertilizing each maternal genotype.
generate.sire.matrix <- function(comp.mat, pollen.freqs){
  # Check to make sure the column names of the compatibility matrix are the same as the names of the pollen vector.
  stopifnot(identical(names(pollen.freqs), colnames(comp.mat)))
  # Generate a matrix of pollen frequencies so that each genotype can be multiplied by each pollen type.
  # It will be a diagonal matrix with the pollen frequencies on the diagonal.
  pollen_mat <- diag(x = pollen.freqs, nrow = length(pollen.freqs), ncol = length(pollen.freqs))
  # Multiply to get the contribution of each pollen to fertilizing each maternal genotype.
  out <- comp.mat %*% pollen_mat
  # Relativize.
  row_totals <- as.vector(rowsum(t(out), rep(1, times = ncol(out))))
  out <- out * 1/row_totals
  rownames(out) <- rownames(comp.mat)
  colnames(out) <- colnames(comp.mat)
  return(out)
}

# The function generate.fertilization.matrix will create a matrix of the frequencies of all crosses.
# It requires the genotype frequencies and the sire matrix.
generate.fertilization.matrix <- function(diplo.freqs, sire.matrix){
  # Check to make sure the names of the genotypes are the same as the row names of the sire matrix.
  stopifnot(identical(names(diplo.freqs), rownames(sire.matrix)))
  # Make a diagonal matrix of the genotype frequencies.
  geno_mat <- diag(x = diplo.freqs, nrow = length(diplo.freqs), ncol = length(diplo.freqs))
  # Now multiply the genotype matrix by the sire matrix to get the frequencies of each fertilization type.
  out <- geno_mat %*% sire.matrix
  rownames(out) <- rownames(sire.matrix)
  colnames(out) <- colnames(sire.matrix)
  return(out)
}

# The function add.distinct.diplos will take a genotype vector with repeated genotypes and uniquify the entries.
# It is useful for generate.new.generation because that function deals with redundant genotypes.
add.distinct.diplos <- function(diplo.freqs, proper.diplos){
  # Standardize names.
  names(diplo.freqs) <- proper.diplos[names(diplo.freqs)]
  # Put names and frequencies in a data frame.
  diplo_frame <- data.frame(name = names(diplo.freqs), freq = diplo.freqs)
  # Add up the frequencies.
  distinct_frame <- ddply(diplo_frame,'name',numcolwise(sum))
  # Turn it back into a vector.
  out <- distinct_frame$freq
  names(out) <- distinct_frame$name
  return(out)
}

# The function generate.new.generation will take a fertilization matrix and create a vector of diploid genotype frequencies.
# It will accomplish this by finding the genotype frequencies of all offspring that inherit the "left" and "right"
# ovules and adding them together.
# It takes redundant diplotype names as an argument.
# It also must be able to recognize redundant permutations.
generate.new.generation <- function(diplos, proper.diplos, fertilization.matrix){
  # Make sure that the diploid genotypes correspond to genotypes in the fertilization matrix.
  stopifnot(identical(paste(diplos[,1], diplos[,2], sep = '_'), rownames(fertilization.matrix)))
  # Figure out the contribution of "left" and "right" ovules to each kind of genotype.
  # Unfurl the fertilization matrix into two vectors with names concatenated from row and column names using left
  # and right ovules. Multiply these by one half and add the ones that have the same names.
  left_contribution <- as.vector(fertilization.matrix) * 0.5
  names(left_contribution) <- paste(rep(diplos[,1], times = ncol(fertilization.matrix)),
                                    rep(colnames(fertilization.matrix), each = nrow(fertilization.matrix)),
                                    sep = '_')
  right_contribution <- left_contribution
  names(right_contribution) <- paste(rep(diplos[,2], times = ncol(fertilization.matrix)),
                                    rep(colnames(fertilization.matrix), each = nrow(fertilization.matrix)),
                                    sep = '_')
  # Add up redundant genotypes into distinct classes.
  left_contribution <- add.distinct.diplos(left_contribution, proper.diplos)
  right_contribution <- add.distinct.diplos(right_contribution, proper.diplos)
  # Identify overlaps.
  overlaps <- match(left_contribution, right_contribution)
  right_unique <- is.na(match(right_contribution, left_contribution))
  # Add the overlapping values to the left contribution and remove them from the right contribution.
  left_contribution[which(!is.na(overlaps))] <- left_contribution[which(!is.na(overlaps))] +
                                                right_contribution[overlaps[!is.na(overlaps)]]
  right_contribution <- right_contribution[right_unique]
  # Merge the two vectors to make the output vector.
  out <- c(left_contribution, right_contribution)
  out <- add.distinct.diplos(out, proper.diplos)
  # Remove impossible genotypes.
  out <- out[paste(diplos[,1], diplos[,2], sep ='_')]
  # Add up genotypes that occur in both vectors.
  # overlap_sums <- left_contribution[which(!is.na(match))] + right_contribution[!is.na(match)]
  # names(overlap_sums) <- names(left_contribution[which(!is.na(match))])
  # Rearrange the right contribution to match the order of genotypes in the left contribution.
  # right_contribution <- right_contribution[names(left_contrib#ution)]
  # Add them to get the genotype frequencies.
  return(out)
}
```
