## Supplementary material for "Diversification or collapse of self-incompatibility haplotypes as a rescue process": All scripts: haplotype_iteration.R

```
# Decide the number of generations to iterate

gens <- 100

# Create vectors to hold the frequencies each generation

pui <- vector(length = gens)
pali <- vector(length = gens)
pdli <- vector(length = gens)
pas <- vector(length = gens)
pds <- vector(length = gens)
pn <- vector(length = gens)
pnplus <- vector(length = gens)
# Define the initial frequencies

pui[1] <- (1/3)/3
pali[1] <- ((1/3)/2)/2
pdli[1] <- ((1/3)/2)/2
pas[1] <- (1/3)/2
pds[1] <- ((1/3)/2)
pn[1] <- 0
pnplus[1] <- 0
# Define the number of allele types in each class

u <- 3
l <- 2

# Make sure that all the frequencies add up each generation.
freqsum <- vector(length = gens)
freqsum[] <- NA
freqsum[1] <- u*pui[1] + l*pali[1] + l*pdli[1] + pas[1] + pds[1] + + pn[1] + pnplus[1]

for(i in 2:gens){
  w <- u * pui[i-1] * (1 - pui[i-1] - pds[i-1] - pnplus[i-1]) +
    l * pali[i-1] * (1 - pali[i-1] - pdli[i-1] - pds[i-1] - pnplus[i-1]) +
    l * pdli[i-1] * (1 - pali[i-1] - pdli[i-1]) +
    pas[i-1] * (1 - pas[i-1] - pn[i-1] - pds[i-1] - pnplus[i-1]) +
    pds[i-1] * (1 - pas[i-1] - pn[i-1] - pds[i-1] - pnplus[i-1]) +
    pn[i-1] * (1 - pas[i-1] - pn[i-1]) +
    pnplus[i-1] * (1 - pds[i-1] - pnplus[i-1])
  pui[i] <- pui[i-1] * (1 - pui[i-1] - pds[i-1] - pnplus[i-1]) / w
  pali[i] <- pali[i-1] * (1 - pali[i-1] - pdli[i-1] - pds[i-1] - pnplus[i-1]) / w
  pdli[i] <- pdli[i-1] * (1 - pali[i-1] - pdli[i-1]) / w
  pas[i] <- pas[i-1] * (1 - pas[i-1] - pn[i-1] - pds[i-1] - pnplus[i-1]) / w
  pds[i] <- pds[i-1] * (1 - pas[i-1] - pn[i-1] - pds[i-1] - pnplus[i-1]) / w
  pn[i] <- pn[i-1] * (1 - pas[i-1] - pn[i-1]) / w
  pnplus[i] <- pnplus[i-1] * (1 - pds[i-1] - pnplus[i-1])/ w
  freqsum[i] <- u*pui[1] + l*pali[1] + l*pdli[1] + pas[1] + pds[1] + pnplus[1]
}

plot(pui, col = 1, ylim = c(0,0.5), type = 'l', xlab = "Generation", ylab = "Frequency")
points(pali, col = 2, type = 'l')
points(pdli, col = 3, type = 'l')
points(pas, col = 4, type = 'l')
points(pds, col = 5, type = 'l')
points(pn, col = 6, type = 'l')
points(pnplus, col = 7, type = 'l')
```
