## Supplementary material for "Diversification or collapse of self-incompatibility haplotypes as a rescue process": All scripts: long-term.R

```
pdf('../figures/stable.pdf')
# Set graphical parameters.
par(lwd = 2, mfrow = c(2,3), mar = c(2,2,3,1), oma = c(5,5,0,0))
# Pick R values.
R <- c(0.01, 0.1, 1, 0.2, 0.3, 0.4)
# Loop through R values.
for(j in 1:6){

  # Load a transition matrix.
  transition <- as.matrix(read.csv(paste('../data/transition_L=2_R=', R[j], '.csv', sep = ''), check.names = F, row.names = 1))

  # The transition probabilities out of the first row were not previously calculated.
  # If there are no unluckies, then it is impossible to produce more haplotypes.
  # The first row is therefore an absorbing state.
  transition[1,] <- c(1, rep(0, (ncol(transition)-1)))
  # The exception might be if the first row is two haplotypes.
  # In this case, the assumption of zero pollen limitation must logically break down because no one can fertilize anyone else.
  # It might therefore make sense for it to be a reflecting boundary.
  if('2' %in% colnames(transition)){
    # If there exists a state with only two haplotypes, make it impossible to transition to it.
    # All density that would transition into 2 is instead reflected to 3.
    transition[,'3'] <- transition[,'3'] + transition[,'2']
    transition[,'2'] <- 0
  }

  # Assuming we start with one unlucky, see what happens when we iterate forward.
  starting_num <- 3
  initial <- rep(0, ncol(transition))
  initial[starting_num-1] <- 1 # Subtract 1 since we start at 2.
  # Assume instead a uniform initial distribution.
  #initial <- rep(1/ncol(transition), ncol(transition))
  names(initial) <- colnames(transition)

  # Now iterate.
  # "Generations" is a misnomer: this is the number of transitions.
  generations <- 1000
  current <- initial
  for(i in 1:generations){
    current <- t(transition) %*% current
    # Try to restore rounding errors. This seems like a weird step, and I think it's probably wrong.
    #current <- current/(sum(current))
  }

  # outpath <- paste('../figures/stable_R=', R, '.pdf', sep = '')

  # pdf(outpath)
  # par(mar = c(8,9,4,2), mgp = c(6, 2, 0), cex.axis = 4, cex.lab = 4, lwd = 2)
  print(current)
  plot(current,
       x = as.numeric(rownames(current)),
       xlab = NA,
       ylab = NA,
       type = 'h',
       yaxt = c('s','n','n',
                's','n','n')[j],
       xaxt = c('n','n','n',
                's','s','s')[j],
       main = eval(substitute(expression(R[conversion] == x), list(x = R[j]))),
       cex.main = 2,
       cex.axis = 2,
       ylim = c(0,1)
       )
}
# Add axis labels.
par(mgp = c(2,1,0))
title(xlab = 'Total haplotypes',
      ylab = 'Frequency',
      cex.lab = 3,
      outer = T)
dev.off()

# Output transition matrix as a heatmap
library(fields)

R <- 0.3
# Load a transition matrix.
transition <- as.matrix(read.csv(paste('../data/transition_L=2_R=', R, '.csv', sep = ''), check.names = F, row.names = 1))

# The transition probabilities out of the first row were not previously calculated.
# If there are no unluckies, then it is impossible to produce more haplotypes.
# The first row is therefore an absorbing state.
transition[1,] <- c(1, rep(0, (ncol(transition)-1)))
# The exception might be if the first row is two haplotypes.
# In this case, the assumption of zero pollen limitation must logically break down because no one can fertilize anyone else.
# It might therefore make sense for it to be a reflecting boundary.
if('2' %in% colnames(transition)){
  # If there exists a state with only two haplotypes, make it impossible to transition to it.
  # All density that would transition into 2 is instead reflected to 3.
  transition[,'3'] <- transition[,'3'] + transition[,'2']
  transition[,'2'] <- 0
}

par(mgp = c(2.75,1,0),
    mar = c(5,5,4,2))

library(viridis)
pdf('../figures/tmatrix.pdf')
# Plot the transition matrix, leaving out impossible states.
image.plot(x = as.numeric(rownames(transition)[2:nrow(transition)]),
           y = as.numeric(colnames(transition)[2:ncol(transition)]),
           z = transition[2:nrow(transition),2:ncol(transition)],
           xlab = 'Haplotypes before transition',
           ylab = 'Haplotypes after transition',
           col = viridis(nrow(transition)),
           legend.lab = 'Transition probability',
           cex.axis = 2,
           cex.lab = 2,
           legend.cex = 2,
           legend.line = 3,
           legend.mar = 7.1
           )
abline(a = 0, b = 1, lwd = 3, col = 'white')
dev.off()

# Simulate trajectories of haplotype number through time using the transition matrix
runs <- 5
duration <- 100
# Create a matrix in which each row is a run and each column is a timestep
number_trajectories <- matrix(nrow = runs, ncol = duration+1)
# Everything starts with three haplotypes
number_trajectories[,1] <- 3
for(j in 1:runs){
  for(i in 2:(duration+1)){
    # Look up the state you're in: that's your row
    # That is also the distribution from which you will be sampling
    number_trajectories[j,i] <- sample(x = as.numeric(colnames(transition)),
                                   size = 1,
                                   prob = transition[as.character(number_trajectories[j,i-1]),]
    )
  }
}
# Plot the trajectories
pdf('../figures/trajectories.pdf')
par(mar = c(5,5,3,2))
plot(NA,
     xlim = c(0,duration),
     ylim = c(0,max(number_trajectories)),
     xlab = 'Time (transitions)',
     ylab = 'Haplotype number',
     cex.lab = 2,
     cex.axis = 2
)
for(j in 1:runs){
  points(number_trajectories[j,], type = 'l')
}
dev.off()
```
