## Supplementary material for "Diversification or collapse of self-incompatibility haplotypes as a rescue process": All scripts: lumped_to_split.R

```
# This script takes a genotype frequency trajectory that DOESN'T pay attention to the frequencies of RNase variants within an SLF class
# and turns it into one that does.
# This obviously only works with some assumption about the relative frequencies of RNase variants since we're just making up that info.
# The script therefore assumes that all RNase variants within an SLF class (besides A and S) are at equal frequencies.
# The frequencies of A and S are already specified by the lumped trajectory.
# Need to double-check that the trajectories produced by splitting a lumped trajectory are the same as those that were split from the start.

source('rnase_functions.R')

# We need to know how many Ls and Us there are.

args <- commandArgs(trailingOnly = T)
L <- eval(parse(text=args[1]))
U <- eval(parse(text=args[2]))

# L <- 2
# U <- 2

# Load the trajectory.

lumped_traj <- read.csv(paste('../data/L=', L, '_U=', U, '_lumped_traj.csv', sep = ''), row.names = 1)

# Generate all the genotypes that can exist.
genos <- generate.genos(L, U, monster = T)

# Now that we have a list of the genotypes, find out which fall into which categories.
of_what_kind <- vector(length = nrow(genos))
of_what_kind[grepl('L', genos[,1]) & grepl('L', genos[,2])] <- 'LL'
of_what_kind[grepl('C', genos[,1]) & grepl('C', genos[,2])] <- 'CC'
of_what_kind[grepl('U', genos[,1]) & grepl('U', genos[,2])] <- 'UU'
of_what_kind[(grepl('L', genos[,1]) & grepl('C', genos[,2])) | (grepl('C', genos[,1]) & grepl('L', genos[,2]))] <- 'LC'
of_what_kind[(grepl('L', genos[,1]) & grepl('U', genos[,2])) | (grepl('U', genos[,1]) & grepl('L', genos[,2]))] <- 'LU'
of_what_kind[(grepl('L', genos[,1]) & grepl('A', genos[,2])) | (grepl('A', genos[,1]) & grepl('L', genos[,2]))] <- 'LA'
of_what_kind[(grepl('L', genos[,1]) & grepl('S', genos[,2])) | (grepl('S', genos[,1]) & grepl('L', genos[,2]))] <- 'LS'
of_what_kind[(grepl('C', genos[,1]) & grepl('U', genos[,2])) | (grepl('U', genos[,1]) & grepl('C', genos[,2]))] <- 'CU'
of_what_kind[(grepl('C', genos[,1]) & grepl('A', genos[,2])) | (grepl('A', genos[,1]) & grepl('C', genos[,2]))] <- 'CA'
of_what_kind[(grepl('C', genos[,1]) & grepl('S', genos[,2])) | (grepl('S', genos[,1]) & grepl('C', genos[,2]))] <- 'CS'
of_what_kind[(grepl('U', genos[,1]) & grepl('A', genos[,2])) | (grepl('A', genos[,1]) & grepl('U', genos[,2]))] <- 'UA'
of_what_kind[(grepl('U', genos[,1]) & grepl('S', genos[,2])) | (grepl('S', genos[,1]) & grepl('U', genos[,2]))] <- 'US'
of_what_kind[(grepl('A', genos[,1]) & grepl('S', genos[,2])) | (grepl('S', genos[,1]) & grepl('A', genos[,2]))] <- 'AS'

# We need to remember that Ls corresponding to Us do not exist.
UsLs <- (L+1):(L+1+U)
bad_Ls <- paste('L', UsLs, sep = '')

# Create a vector of which genotypes are extant.
extant <- rep(T, nrow(genos))
extant[genos[,1] %in% bad_Ls | genos[,2] %in% bad_Ls] <- F

# Now that we know which kind is which, we just need to divide up the lumped categories into the split categories.
# They should be divided evenly, I think.
# If they aren't, then that already contradicts the assumption that they would have shared the same trajectory.

split_traj <- matrix(nrow = nrow(lumped_traj), ncol = nrow(genos))
colnames(split_traj) <- paste(genos[,1], genos[,2], sep = '_')

for(i in 1:nrow(split_traj)){
  split_traj[i,(of_what_kind == 'LL' & extant)] <- lumped_traj[i,'LL'] / sum(of_what_kind == 'LL' & extant)
  split_traj[i,(of_what_kind == 'LC' & extant)] <- lumped_traj[i,'LC'] / sum(of_what_kind == 'LC' & extant)
  split_traj[i,(of_what_kind == 'LU' & extant)] <- lumped_traj[i,'LU'] / sum(of_what_kind == 'LU' & extant)
  split_traj[i,(of_what_kind == 'LA' & extant)] <- lumped_traj[i,'LA'] / sum(of_what_kind == 'LA' & extant)
  split_traj[i,(of_what_kind == 'LS' & extant)] <- lumped_traj[i,'LS'] / sum(of_what_kind == 'LS' & extant)
  split_traj[i,(of_what_kind == 'CC' & extant)] <- lumped_traj[i,'CC'] / sum(of_what_kind == 'CC' & extant)
  split_traj[i,(of_what_kind == 'CU' & extant)] <- lumped_traj[i,'CU'] / sum(of_what_kind == 'CU' & extant)
  split_traj[i,(of_what_kind == 'CA' & extant)] <- lumped_traj[i,'CA'] / sum(of_what_kind == 'CA' & extant)
  split_traj[i,(of_what_kind == 'CS' & extant)] <- lumped_traj[i,'CS'] / sum(of_what_kind == 'CS' & extant)
  split_traj[i,(of_what_kind == 'UU' & extant)] <- lumped_traj[i,'UU'] / sum(of_what_kind == 'UU' & extant)
  split_traj[i,(of_what_kind == 'UA' & extant)] <- lumped_traj[i,'UA'] / sum(of_what_kind == 'UA' & extant)
  split_traj[i,(of_what_kind == 'US' & extant)] <- lumped_traj[i,'US'] / sum(of_what_kind == 'US' & extant)
  split_traj[i,(of_what_kind == 'AS' & extant)] <- lumped_traj[i,'AS'] / sum(of_what_kind == 'AS' & extant)
}

# Remove NAs.
split_traj[is.na(split_traj)] <- 0

out_name <- paste('../data/L=', L, '_U=', U, '_split_traj.csv', sep = '')
write.csv(split_traj, file = out_name)
```
