## Supplementary material for "Diversification or collapse of self-incompatibility haplotypes as a rescue process": All scripts: make_lumped_trajectory.R

```
# This script iterates the population through the evolutionary rescue phase.

# Load the functions.

source('rnase_functions.R')

# # Load the starting population after equilibration.
# 
# 
# traj_name <- paste('../data/L=', L, '_U=', U, '_initial.csv', sep = '')
# 
# # R FORCES EVERYTHING TO BE A DATA FRAME EVEN WHEN YOU EXPLICITLY TELL IT NOT to BE.
# fake_population <- (read.csv(traj_name))
# population <- as.numeric(read.csv(traj_name))
# names(population) <- colnames(fake_population)

# Instead of loading a pre-equilibrated trajectory, load a trajectory before equilibration occurs
args <- commandArgs(trailingOnly = T)
L <- eval(parse(text=args[1]))
U <- eval(parse(text=args[2]))
luck_frac <- 0.01 # The fraction of each of the L haplotypes that actually has the new key.
spite_frac <- 0.01 # The fraction of the lock-mutant's ancestor that actually mutated to the lock-mutant.

initialize(L,
           U,
           luck_frac,
           spite_frac)

# Now we need to introduce the monster allele.
# The most correct way to do this would be to turn all of the As into a new class of U.
# The problem is that the As won't be at the same frequency as the other Us.
# Another way would be to change the compatibility relationships, though this could have unforeseen consequences.


# Create trajectories of diploid genotype frequencies.
# Plot trajectories of haplotype frequencies.

my_gens <- 4000 # The number of generations you want to track.
traj <- track.diplos(population, my_gens, monster = T)

traj_name <- paste('../data/L=', L, '_U=', U, '_lumped_traj.csv', sep = '')
write.csv(traj, traj_name)
```
