## Supplementary material for "Diversification or collapse of self-incompatibility haplotypes as a rescue process": All scripts: make_rescue_barplot.R

```
library(knitr)

# Choose graphical parameters.
par(mar = c(8,9,4,2), mgp = c(6, 2, 0), cex.axis = 4, cex.lab = 4, lwd = 2)

# Load rescue data.

rescue <- read.csv('../data/rescue.csv', check.names = F)

# Let's hold L = 2.
# The rescue probability is higher for low L, except when L < 2, in which case the new RNase can't invade.
# Let's also hold U = 20 because diversification at high diversity is absent in most models.
data <- rescue[,2:ncol(rescue)]

# Let's vary N while holding gene conversion constant.
vary_N <- data[data$conversion_rate == (10^-5) & ((rescue$L == 2) & rescue$U == 2),c('N','rescue_probability')]
# Sort by N.
vary_N <- vary_N[order(vary_N$N),]
barplot(height = (vary_N$rescue_probability)[1:5], names.arg = (vary_N$N)[1:5],  beside = T, ylab = 'P(all rescued)', ylim = c(0,1), xlab = 'N',
        col = 'red')

# Let's vary the gene conversion rate while holding N constant.
vary_conv <- data[(data$N == 10000) & ((rescue$L == 2) & rescue$U == 2),]
vary_conv <- vary_conv[nrow(vary_conv):1,] # Flip it so the smaller ones end up on the left.
# Sort by conversion rate.
vary_conv <- vary_conv[order(vary_conv$conversion_rate),]
barplot(height = vary_conv$rescue_probability, names.arg = vary_conv$conversion_rate, ylab = 'P(all rescued)', xlab = 'Conversion rate',
        col = 'red')

# Let's vary U while holding L constant at 2, N at 2000, and conversion rate at 10^-5.
vary_U <- rescue[(rescue$L == 2) & (rescue$N==10000) & rescue$conversion_rate==(10^-5),2:ncol(rescue)]
barplot(height = vary_U$rescue_probability, names.arg = vary_U$U, xlab = 'Normal haplotypes', ylab = 'P(all rescued)', ylim = c(0,1),
        col = 'red')
```
