## Supplementary material for "Diversification or collapse of self-incompatibility haplotypes as a rescue process": All scripts: make_rescue_curves.R

```
library(knitr)

# Load rescue data.

rescue <- read.csv('../data/rescue.csv', check.names = F)

# Remove the first column (a bunch of ones)
rescue <- rescue[,2:ncol(rescue)]

# Lump the N and gene conversion rates into a product column.
Rconversion <- rescue$N * rescue$conversion_rate
rescue <- data.frame(L = rescue$L, U = rescue$U, Rconversion, rescue_probability = rescue$rescue_probability)
rm(Rconversion)

# Order by Rconversion

rescue <- rescue[order(rescue$Rconversion),]

# Filter out the tiny ones.
rescue <- rescue[(rescue$Rconversion >= 0.1),]

# Filter out the huge ones.
rescue <- rescue[(rescue$Rconversion <= 10),]


# Generate a palette.
#my_pal = c(2,3,4,5)

# Loop through different L values.

# Choose graphical parameters.
pdf(file = '../figures/rescue_curves.pdf')
par(lwd = 2, mfrow = c(2,2), mar = c(4,3,2,1), oma = c(3,4,0,0),
    cex.axis = 2, cex.lab = 2)
# Plot blanks.
i <- 1
for(i in 1:4){
  current_main <- substitute(expression(n[L] == x), list(x = unique(rescue$L)[i]))
  plot(NA,
       xlab = NA,
       ylab = NA,
       xlim = c(min(log(rescue$Rconversion, base = 10)),max(log(rescue$Rconversion, base = 10))),
       ylim = c(min(log(rescue$rescue_probability, base = 10))-5,max(log(rescue$rescue_probability, base = 10))),
       main = eval(current_main))
}
i <- 1
a <- c(1,2,1,2)
b <- c(1,1,2,2)
for(current_L in unique(rescue$L)){
  # Switch between the four plots.
  par(mfg = c(a[i], b[i]))
  # Plot for U = 2
  i <- i+1
  points(x = log(rescue[rescue$U == 2 & rescue$L == current_L, 'Rconversion'], base = 10),
         y = log(rescue[rescue$U == 2  & rescue$L == current_L, 'rescue_probability'], base = 10),
         type = 'l',
         lty = 1)
  
  # Plot for U = 5
  points(x = log(rescue[rescue$U == 5 & rescue$L == current_L, 'Rconversion'], base = 10),
         y = log(rescue[rescue$U == 5  & rescue$L == current_L, 'rescue_probability'], base = 10),
         type = 'l',
         # col = my_pal[2],
         lty = 2)

  # U = 10
  points(x = log(rescue[rescue$U == 10 & rescue$L == current_L, 'Rconversion'], base = 10),
         y = log(rescue[rescue$U == 10  & rescue$L == current_L, 'rescue_probability'], base = 10),
         type = 'l',
         # col = my_pal[3],
         lty = 3)

  # U = 20
  points(x = log(rescue[rescue$U == 20 & rescue$L == current_L, 'Rconversion'], base = 10),
         y = log(rescue[rescue$U == 20  & rescue$L == current_L, 'rescue_probability'], base = 10),
         type = 'l',
         # col = my_pal[4],
         lty = 4)

}
# Create a legend.
par(mfg = c(1,1))
  legend('bottomright',
         legend = c(expression(n[U] == 2),
                    expression(n[U] == 5),
                    expression(n[U] == 10),
                    expression(n[U] == 20)),
         cex = 0.9,
         lty = 1:4,
         bg = 'white')

par(mgp = c(1,1,0))
title(xlab = expression(paste(log[10], (R[conversion]))),
      ylab = expression(paste(log[10], (P[expansion]))),
      outer = T,
      cex.lab = 2)

dev.off()
```
