## Supplementary material for "Diversification or collapse of self-incompatibility haplotypes as a rescue process": All scripts: make_rescue_table.R

```
library(knitr)

# Load rescue data.

rescue <- read.csv('../data/rescue_N=100.csv', row.names = 1, check.names = F)

# Put them in a table.

sink('../tables/rescue_prob_N=100.txt')
kable(t(rescue), format = 'latex', caption = 'caption', row.names = T)
sink()

# Load rescue data.

rescue <- read.csv('../data/rescue_N=1000.csv', row.names = 1, check.names = F)

# Put them in a table.

sink('../tables/rescue_prob_N=1000.txt')
kable(t(rescue), format = 'latex', caption = 'caption', row.names = T)
sink()

# Load rescue data.

rescue <- read.csv('../data/rescue_N=10000.csv', row.names = 1, check.names = F)

# Put them in a table.

sink('../tables/rescue_prob_N=10000.txt')
kable(t(rescue), format = 'latex', caption = 'caption', row.names = T)
sink()

# Load rescue data.

rescue <- read.csv('../data/rescue_N=1e+05.csv', row.names = 1, check.names = F)

# Put them in a table.

sink('../tables/rescue_prob_N=1e+05.txt')
kable(t(rescue), format = 'latex', caption = 'caption', row.names = T)
sink()

# Load rescue data.

rescue <- read.csv('../data/rescue_N=1e+06.csv', row.names = 1, check.names = F)

# Put them in a table.

sink('../tables/rescue_prob_N=1e+06.txt')
kable(t(rescue), format = 'latex', caption = 'caption', row.names = T)
sink()
```
