## Supplementary material for "Diversification or collapse of self-incompatibility haplotypes as a rescue process": All scripts: migration.R

```
# Here's a little side-project.
# I don't intend to put this in this manuscript.
# I'm actually hoping it's enough to write a different manuscript about.
# The idea is to model the introduction of foreign locks and keys.
# It would answer the question, how much introgression should we expect as a function of migration rate and S-allele overlap?
# This could include both introgression of S-alleles and introgression of neutral sites, either linked or unlinked.
# It would be difficult to add additional sites while maintaining the same underlying R processes because we modeled only one locus.
# However, it might be possible to say something about it given a genotype history.


source('rnase_functions.R')

args <- commandArgs(trailingOnly=TRUE)

# Choose parameters
L <- eval(parse(text=args[1]))
U <- eval(parse(text=args[2]))
luck_frac <- 1
spite_frac <- 1
my_generations <- 4000

# Initialize the population from those parameters.
my_genos <- generate.genos(L,U)
my_pollen_prod <- generate.pollen.production(my_genos)
my_hap_comp <- generate.haplo.compatibility(my_genos)
my_dip_comp <- generate.diplo.compatibility(my_hap_comp, my_genos)
my_geno_dist <- sim.initialize(my_genos, luck_frac = luck_frac, spite_frac = spite_frac)
names(my_geno_dist) <- paste(my_genos[,1], my_genos[,2], sep = '_')

# Determine the expected genotype frequency trajectories during the rescue phase.
expected_trajectory <- iterate(my_genos, my_geno_dist, my_pollen_prod, my_dip_comp, generations = my_generations)
trajectory_name <- paste('../data/', 'L=', L, '_U=', U, '_traj.csv', sep = '')
```
