## Supplementary material for "Diversification or collapse of self-incompatibility haplotypes as a rescue process": All scripts: outcome_distribution.R

```
# This script calculates the probability distribution of the number of rescued haplotypes.
# It is very similar to rescue_from_trajectory except that it calculates the whole distribution not just the probability of a particular event.

pdf('../figures/outcomes.pdf')
# Set up a plotting window.
par(mfrow = c(3,3),
    oma = c(4,5,0,0),
    mar = c(2,5,2,2),
    cex.main = 2,
    cex.axis = 2,
    cex.lab = 2,
    mgp = c(3,1,0),
    lwd = 2)

# Hold N and U constant.
N <- 1000
U <- 10

# Decide on L and conversion_rate parameters to loop through.
Ls <- c(2,3,5)
conversion_rates <- c(0.00001, 0.00002, 0.0001)
# Create a vector of above-figure text, which tells the reader how many Ls there are.
# above_figure <- c(substitute(expression(n[L] == x), list(x = Ls[1])),
#                   substitute(expression(n[L] == x), list(x = Ls[2])),
#                   substitute(expression(n[L] == x), list(x = Ls[3])),
#                   NA,NA,NA,NA,NA,NA)
# Create a vector of beside-figure text, which tells the reader the conversion rate.
beside_figure <- c(paste('R =', N*conversion_rates[1]), NA, NA,
                   paste('R =', N*conversion_rates[2]), NA, NA,
                   paste('R =', N*conversion_rates[3]), NA, NA)

# Loop through L.
# Create an index for above- and beside-figure text.
param_ind <- 1
for(conversion_rate in conversion_rates){
  for(L in Ls){
    
    source('rnase_functions.R')
    
    # Take the filepath to the trajectory as an argument.
    # This allows us to calculate the rescue probability from multiple trajectories with a bash script.
    # args <- commandArgs(trailingOnly=TRUE)
    # L <- eval(parse(text=args[1]))
    # U <- eval(parse(text=args[2]))
    # N <- eval(parse(text=args[3]))
    # conversion_rate <- eval(parse(text=args[4]))
    
    trajectory_path <- paste('../data/', 'L=', L, '_U=', U, '_split_traj.csv', sep = '')
    survival_path <- paste('../data/', 'L=', L, '_U=', U, '_survive.csv', sep = '')
    
    # Load the expected trajectory.
    
    expected_trajectory <- as.matrix(read.csv(trajectory_path, row.names = 1))
    
    # Load the probability of survival through time.
    
    expected_survival <- as.matrix(read.csv(survival_path, row.names = 1, header = F))
    
    # Load a separate survival probability for A.
    
    survival_path_A <- paste('../data/', 'L=', L, '_U=', U, '_survive_A.csv', sep = '')
    expected_survival_A <- as.matrix(read.csv(survival_path_A, row.names = 1, header = F))
    
    # Identify which genotypes could generate rescuing convertants.
    # These genotypes are heterozygous for a U and an L.
    
    is_L <- grepl('L', colnames(expected_trajectory))
    first_U_start <- paste('^U_', L+1, sep = '')
    first_U_end <- paste('U', L+1, '$', sep = '')
    is_first_U <- grepl(first_U_start, colnames(expected_trajectory)) | grepl(first_U_end, colnames(expected_trajectory))
    is_A <- grepl('A', colnames(expected_trajectory))
    is_convertible_U <- is_L & is_first_U
    convertible_U <- rowSums(expected_trajectory[,is_convertible_U])
    is_convertible_A <- is_L & is_A
    # Create a vector of the frequency of convertible individuals through time.
    convertible_A <- rowSums(expected_trajectory[,is_convertible_A])
    
    # Multiply by the population size to get the number of convertible individuals through time.
    convertible_num <- N * convertible_U
    convertible_A_num <- N * convertible_A
    
    # Get the probability of NOT surviving.
    loss <- 1 - expected_survival
    loss_A <- 1 - expected_survival_A
    
    # Find the probability that all COPIES are lost.
    lost <- loss ^ (convertible_num * conversion_rate)
    lost_A <- loss_A ^ (convertible_A_num * conversion_rate)
    all_lost <- prod(lost)
    all_lost_A <- prod(lost_A)
    
    rescue <- 1 - all_lost
    rescue_A <- 1 - all_lost_A
    
    # Now we know p in the binomial.
    # There are basically two random variables: the number of rescued As (0 or 1) and the number of rescued Us.
    # All we need to do is make two distributions for the number of Us rescued: one conditional on A being lost and one on being rescued.
    
    # The distributions are the same except that if A is rescued, then the distribution shifts up one unit.
    U_rescue_dist <- dbinom(0:(U), U, prob = rescue)
    
    # Now get the full distribution.
    rescue_dist <- vector(length = length(U_rescue_dist)+1)
    names(rescue_dist) <- 0:(U+1)
    rescue_dist[1:(length(rescue_dist)-1)] <- rescue_dist[1:(length(rescue_dist)-1)] + (U_rescue_dist * all_lost_A)
    rescue_dist[2:(length(rescue_dist))] <- rescue_dist[2:(length(rescue_dist))] + (U_rescue_dist * rescue_A)
    
    # Just adding the number of lucky alleles to the number of rescued alleles gives you the final number of alleles.
    final_dist <- rescue_dist
    names(final_dist) <- (0:(U+1)) + L
    
    # Before plotting, save the outcome distribution as a row in a transition matrix.
    # If we're going to run this script repeatedly, we might need to find some way to make sure we don't get duplicate rows.
    # One way would be to open the transition matrix file and overwrite any row with the same parameters.
    # Intuitively, this seems kind of wasteful, but I guess write.csv() has to open the file to edit it anyway.
    
    # Create an entry into the transition matrix.
    # The minimum will be L and the maximum will be L+40.
    states <- L:(L+40)
    # If the final element of final_dist extends beyond forty, collapse everything >= forty into the forty category.
    if(as.numeric(names(final_dist)[length(final_dist)]) > (L+40)){
      entry <- c(final_dist[1:which(as.numeric(names(final_dist))==(L+39))],sum(final_dist[which(as.numeric(names(final_dist))==(L+40)):length(final_dist)]))
    }else{
      entry <- c(final_dist, rep(0, 41-length(final_dist)))
    }
    names(entry) <- states
    
    # Create a transition matrix file if it doesn't already exist or open it if it does.
    # This file will be named after L and R (the population-wide rate of gene conversion, N * conversion_rate) on the assumption that it's transitioning between different U states.
    transition_path <- paste('../data/transition_L=', L, '_R=', as.character(N*conversion_rate), '.csv', sep = '')
    # run <- function(){
    if(file.exists(transition_path)){
      transition_matrix <- read.csv(transition_path, row.names = 1, check.names = F)
      # Now add the new entry, overwriting anything that currently occupies that row.
      transition_matrix[as.character(L+U),] <- entry
    }else{
      transition_matrix <- matrix(nrow = length(states), ncol = length(states))
      print(paste('L=', L, 'U=', U, 'N=', N, 'conv=', conversion_rate, sep = ''))
      rownames(transition_matrix) <- states
      colnames(transition_matrix) <- states
      # Now add the entry, overwriting anything that currently occupies that row.
      transition_matrix[as.character(L+U),] <- entry
    }
    # }
    # debug(run)
    # run()
    # With the transition matrix updated, write it to a file.
    # write.csv(transition_matrix, file = transition_path)
    
    # Plot.
    # name_of_plot <- paste('../figures/', 'L=', L, '_U=', U, '_N=', N, '_conv=', conversion_rate, '_outcome.pdf', sep = '')
    # pdf(name_of_plot)
    
    # Only plot the axes on the edges of the multipanel figure.
    if(L == 2){
      current_yaxt <- 's'
    }else{
      current_yaxt <- 'n'
    }
    if(conversion_rate == 0.0001){
      current_xaxt <- 's'
    }else{
      current_xaxt <- 'n'
    }
    # Put the right nL above the top panels.
    if(!is.na(above_figure[param_ind])){
      current_main <- eval(substitute(expression(n[L] == x), list(x = L)))
    }else{
      current_main <- above_figure[param_ind]
    }
    barplot(final_dist,
            xlab = NA,
            ylab = beside_figure[param_ind],
            xaxt = current_xaxt,
            yaxt = current_yaxt,
            ylim = c(0,1),
            main = if(!is.na(above_figure[param_ind])){eval(substitute(expression(n[L] == x), list(x = L)))}else{NA}
            # main = paste('L = ', L, ', R = ', N*conversion_rate, sep = '')
    )
    param_ind <- param_ind + 1
  }
}
par(mgp = c(2,1,0))
title(xlab = 'Total haplotypes',
      ylab = 'Probability',
      cex.lab = 3,
      outer = T)
dev.off()
```
