## Supplementary material for "Diversification or collapse of self-incompatibility haplotypes as a rescue process": All scripts: pollen_limitation.R

```
# Start by listing what you know will be arbitrary parameters

##################
### PARAMETERS ###
##################

# The number of ovules per carpel
ovules <- 10

# The number of pollen grains that can fit on a stigma
# We assume that there is enough total pollen to cover the stigma, even if not all of it is compatible
# We're going to vary the pollen/ovule ratio by varying pollen
pollen <- 100:1000

# The number of lucky haplotypes
nL <- 2

# The proportion of lucky haplotypes within L + C
dL <- c(0.01,0.1,0.5)

# The number of unlucky haplotypes
nU <- c(3, 9, 27)

# Proportion of pollen that is self-pollen
# We assume prior selfing at a given rate (might expand to allow competing selfing).
#prior = T
# Assuming prior selfing, a fraction of SC plants' ovules are wasted prior to receiving cross pollen.
#self <- 0.1

# Put all parameter combinations in a matrix, and leave empty columns for the outputs.
params <- matrix(nrow = length(ovules)*length(pollen)*length(nL)*length(dL)*length(nU),
                 ncol = 7)
colnames(params) <- c('ovules','pollen','nL','dL','nU','damFit','sireFit')

k <- 1
for(ov in ovules){
  for(pol in pollen){
    for(l in nL){
      for(d in dL){
        for(u in nU){
          params[k,] <- c(ov,pol,l,d,u,NA,NA)
          k <- k+1
        }
      }
    }
  }
}

######################
### DERIVED VALUES ###
######################

for(i in 1:nrow(params)){
# The probability that a given pollen grain is compatible with the dam's genotype
# This depends on the dam's genotype
#!!!!!!!!!!!!!!!!!!!!!!!!!!!###
### THINK ABOUT SELF POLLEN ###
#!!!!!!!!!!!!!!!!!!!!!!!!!!!!!#
compUM <- (params[i,'nL']*params[i,'dL'])/(params[i,'nL'] + params[i,'nU']) # If the other allele is a U, the pollen must merely be an L
compLM <- ((params[i,'nL'] - 1)*params[i,'dL'])/(params[i,'nL'] + params[i,'nU']) # If the other allele is an L, the pollen must be a different L
compUU <- (params[i,'nL'] + params[i,'nU'] - 2)/(params[i,'nL'] + params[i,'nU']) # All other combinations can take all but matching alleles

# The distribution of compatible pollen on the stigma is binomial with probability of success compM

# All these genotypes are SI, so they all get 
distCompUM <- dbinom(0:params[i,'pollen'], size = params[i,'pollen'], prob = compUM) 
distCompLM <- dbinom(0:params[i,'pollen'], size = params[i,'pollen'], prob = compLM)
distCompUU <- dbinom(0:params[i,'pollen'], size = params[i,'pollen'], prob = compUU)


# Get a function that relates the number of compatible pollen grains to the number of fertilized ovules
unfert <- ovules - (0:params[i,'pollen'])
unfert[unfert < 0] <- 0
fert <- ovules - unfert

# The vector fert relates the number of compatible pollen grains (index of the vector minus one) to the
# number of ovules fertilized (a given element of the vector)
# For example, the first element is always zero because no ovules are fertilized when no compatible pollen
# is received.

# Assuming prior selfing, SC plants further reduce their ovule success by (selfing rate) * (total ovules)
# homSelfFert <- fert - self * ovules
# hetSelfFert <- fert - self * ovules / 2

# Ovule success is the proportion of cross-fertilized ovules to all ovules
ovuleSuccess <- fert/ovules
#homSelfSuccess <- homSelfFert/ovules
#hetSelfSuccess <- hetSelfFert/ovules

# plot(x = success,
#      y = distCompUM,
#      type = 'l')
# points(x = success,
#      y = distCompLM,
#      type = 'l')
# points(x = success,
#      y = distCompUU,
#      type = 'l')

# sum(success*distCompUM)
# sum(success*distCompLM)
# sum(success*distCompUU)

# Assuming that UM and LM occur proportional to the frequencies of U and L,
# which should be the case immediately upon the origin of the mutant,
# we can calculate the mean ovule success of mutant-bearing genotypes
params[i,'damFit'] <- ((sum(ovuleSuccess*distCompLM))*(params[i,'nL']*params[i,'dL'])/(params[i,'nL'] + params[i,'nU']) +
                       (sum(ovuleSuccess*distCompUM))*(params[i,'nU'] + params[i,'nL']*(1-params[i,'dL']))/(params[i,'nL']+params[i,'nU'])
                      ) /
                      sum(ovuleSuccess*distCompUU)

# Now calculate pollen success.
params[i,'sireFit'] <- 1 /
                       ((params[i,'nL'] + params[i,'nU'] - 2) / (params[i,'nL'] + params[i,'nU']))
}

# Dam and sire fitness are calculated for the MUTANT relative to ALL NONMUTANT SIs.
# Assuming the intermediate I and mutant M haplotypes occur at negligible frequencies,
# the ovule success of double-nonmutants should equal the mean ovule success,
# and the pollen success of nonmutant pollen should equal the mean pollen success.
# Might be worth expanding to account for explicit genotype frequencies.

############
### PLOT ###
############

# Make 9 plots of net fitness vs. pollen supply,
# where each plot assumes a different nU.
# pdf('../figures/pollen_limitation.pdf')
# par(mfrow = c(3,3))
# for(u in nU){
#   for (d in dL){
#     desired <- params[,'nU'] == u & params[,'dL'] == d
#     plot(x = params[desired,'pollen']/params[desired,'ovules'],
#          y = params[desired,'sireFit']/2 + params[desired,'damFit']/2,
#          xlab = 'Pollen received per ovule',
#          ylab = 'Mutant fitness',
#          ylim = c(0,2),
#          type = 'l',
#          lwd = 2,
#          main = paste('nU = ', u,
#                       ', ',
#                       'dL = ', d,
#                       sep = '')
#     )
#     abline(h = 1)
#   }
# }
# dev.off()

# Tidy version
library(tidyverse)

pdf('../figures/pollen_limitation.pdf')
data.frame(params) %>%
  mutate(Fitness = sireFit/2+damFit/2) %>%
  mutate(nU = factor(nU),
         Pollen = pollen,
         Ovules = ovules) %>%
  ggplot(aes(x = Pollen/Ovules, y = Fitness, lty = nU))+
           geom_line(size = 1)+
           facet_wrap(. ~dL,labeller = 'label_both')+
           scale_y_continuous(limits = c(0,2))+
           geom_hline(yintercept = 1)+
           theme_light()+
           theme(axis.text=element_text(size=16), axis.title=element_text(size=16))
dev.off()
```
