## Supplementary material for "Diversification or collapse of self-incompatibility haplotypes as a rescue process": All scripts: preconversion.R

```
# This script finds the equilibrium genotype frequencies after the introduction of an RNase mutation but before gene conversion.

# Load the functions.

source('rnase_functions.R')

# Initialize the population.

args <- commandArgs(trailingOnly = T)
L <- eval(parse(text=args[1]))
U <- eval(parse(text=args[2]))

my_gens <- 4000 # The number of generations you want to track.
# L <- 2 # The number of haplotypes that have a variant with the new key.
# U <- 2 # The number of haplotypes that have no variant with the new key.
luck_frac <- 0.01 # The fraction of each of the L haplotypes that actually has the new key.
spite_frac <- 0.01 # The fraction of the lock-mutant's ancestor that actually mutated to the lock-mutant.

initialize(L,
           U,
           luck_frac,
           spite_frac)

# Create trajectories of diploid genotype frequencies.
traj <- track.diplos(population, my_gens)

# Plot trajectories of haplotype frequencies.
al_traj <- matrix(nrow = nrow(traj), ncol = 5)
colnames(al_traj) <- c('L', 'C', 'U', 'A', 'S')
for(i in 1:nrow(traj)){
  al_traj[i,] <- diplos.to.haplos(traj[i,])
}

traj_name <- paste('../data/L=', L, '_U=', U, '_initial.csv', sep = '')
write.table(t(traj[nrow(traj),]), traj_name, sep = ',')
```
