## Supplementary material for "Diversification or collapse of self-incompatibility haplotypes as a rescue process": All scripts: rescue_from_trajectory.R

```
# This script calculates the probability that all doomed haplotypes are rescued.

source('rnase_functions.R')

# Take the filepath to the trajectory as an argument.
# This allows us to calculate the rescue probability from multiple trajectories with a bash script.
args <- commandArgs(trailingOnly=TRUE)
L <- eval(parse(text=args[1]))
U <- eval(parse(text=args[2]))
N <- eval(parse(text=args[3]))
conversion_rate <- eval(parse(text=args[4]))

trajectory_path <- paste('../data/', 'L=', L, '_U=', U, '_split_traj.csv', sep = '')
survival_path <- paste('../data/', 'L=', L, '_U=', U, '_survive.csv', sep = '')

# Load the expected trajectory.

expected_trajectory <- as.matrix(read.csv(trajectory_path, row.names = 1))

# Load the probability of survival through time.

expected_survival <- as.matrix(read.csv(survival_path, row.names = 1, header = F))

# Load a separate survival probability for A.

survival_path_A <- paste('../data/', 'L=', L, '_U=', U, '_survive_A.csv', sep = '')
expected_survival_A <- as.matrix(read.csv(survival_path_A, row.names = 1, header = F))

# Identify which genotypes could generate rescuing convertants.
# These genotypes are heterozygous for a U and an L.

is_L <- grepl('L', colnames(expected_trajectory))
first_U <- paste('U', L+1, sep = '')
is_first_U <- grepl(first_U, colnames(expected_trajectory))
is_A <- grepl('A', colnames(expected_trajectory))
# PROBLEM HERE vvvvvvvvvvvv
# The probability that a given U is rescued depends on the supply of convertants of THAT TYPE of U, not on all Us.
# Any two EXTANT convertible genotypes (other than A) should be at equal frequencies.
is_convertible_U <- is_L & is_first_U
convertible_U <- rowSums(expected_trajectory[,is_convertible_U])
is_convertible_A <- is_L & is_A
# Create a vector of the frequency of convertible individuals through time.
convertible_A <- rowSums(expected_trajectory[,is_convertible_A])

# Multiply by the population size to get the number of convertible individuals through time.
convertible_num <- N * convertible_U
convertible_A_num <- N * convertible_A

# Get the probability of NOT surviving.
loss <- 1 - expected_survival
loss_A <- 1 - expected_survival_A

# Find the probability that all are lost.
lost <- loss ^ (convertible_num * conversion_rate)
lost_A <- loss_A ^ (convertible_A_num * conversion_rate)
all_lost <- prod(lost)
all_lost_A <- prod(lost_A)

# The probability that not all are lost is the complement.
rescue <- 1 - all_lost
rescue_A <- 1 - all_lost_A
#rescue

# Now that we've got the probability that one haplotype is rescued, calculate the probability that they're all rescued.
# Assume that these are independent, identically distributed events.
# First, we need to figure out how many Us there are.
haplo_names <- names(hap.from.dip(expected_trajectory[1,]))
hap_is_U <- grepl('U', haplo_names)
U <- sum(hap_is_U)

# The probability that all haplotypes are rescued is the product of their individual rescue probabilities.
rescue_all_haps <- rescue_A * (rescue ^ U)
#rescue_all_haps

out <- t(c(L, U, N, conversion_rate, rescue_all_haps))
write.table(out, file = '../data/rescue.csv', append = T, col.names = F, sep = ',')
```
