## Supplementary material for "Diversification or collapse of self-incompatibility haplotypes as a rescue process": All scripts: rescue_prob.R

```
# This script explores parameter space and calculates the probability that all hapltoypes are rescued.

# Choose a population size.

N <- 10000

# Load the functions.

source('rnase_functions.R')

# Create a matrix to hold the rescue probabilities.
# This matrix will vary the number of doomed alleles and the per-heterozygote rate of gene conversion.
rescue_mat <- matrix(nrow = 6, ncol = 10)
rownames(rescue_mat) <- c(10^-8,
                          10^-7,
                          10^-6,
                          10^-5,
                          10^-4,
                          10^-3
                          )
colnames(rescue_mat) <- 1:10

# Calculate rescue probabilities.
for(U in 1:10){
  for(conversion.rate in c(10^-8,
                           10^-7,
                           10^-6,
                           10^-5,
                           10^-4,
                           10^-3
                           )
  ){

# Initialize

my_gens <- 1000
L <- 2
# U is defined already because it is an iterator.
luck_frac <- 1 # The fraction of the lucky RNase haplotypes that also have the new SLF.
spite_frac <- 1 # The fraction of the ancestral RNase that is replaced with the novel RNase.

initialize(L,
           U,
           luck_frac,
           spite_frac)

# Create trajectories of diploid genotype frequencies.
# Plot trajectories of haplotype frequencies.

traj <- track.diplos(population, my_gens)

# To calculate the supply of gene convertants of a particular type, we need a vector of the frequencies of diploid genotypes
# that can produce that convertant.

my_convertible_traj <- traj[,'LU']/U

my_supply <- generate.supply(my_convertible_traj, N, conversion.rate)
my_survive <- determine.survival.prob(traj,N)

# You can back-calculate the selection coefficients from the survival = 2s approximation and put the resulting s into series.solver
my_series_survive <- sapply(my_survive/2, series.solver, guesses = seq(from = 0, to = 1, by = 0.01), terms = 100)
my_series_rescue <- rescue.prob(my_supply, my_series_survive)

# This is the probability that all rescues occur, assuming they're independent of each other.
total_rescue <- my_series_rescue ^ U
rescue_mat[as.character(conversion.rate),as.character(U)] <- total_rescue
  }
}
write.csv(rescue_mat, paste('../data/rescue_N=', N, '.csv', sep = ''))
```
