## Supplementary material for "Diversification or collapse of self-incompatibility haplotypes as a rescue process": All scripts: rnase_functions.R

```
########################
# Function definitions #
########################

# The function one.generation iterates the population for one generation.
# The population is a vector of diploid genotype frequencies.
# NEW FEATURE: takes into account whether the monster haplotype has arrived.
# If it has, the only difference is that the ancestor functions like just another U.

one.generation <- function(pop, monster = F){
  # I should probably make U and L arguments, but for now I will leave them as global variables.
  current.pop <- pop
  # Create a pollen pool.
  pollen.pool <- c(current.pop['LL']+current.pop['LC']/2+current.pop['LU']/2+current.pop['LA']/2+current.pop['LS']/2,
                   current.pop['LC']/2+current.pop['CC']+current.pop['CU']/2+current.pop['CA']/2+current.pop['CS']/2,
                   current.pop['LU']/2+current.pop['CU']/2+current.pop['UU']+current.pop['UA']/2+current.pop['US']/2,
                   current.pop['LA']/2+current.pop['CA']/2+current.pop['UA']/2+current.pop['AS']/2,
                   current.pop['LS']/2+current.pop['CS']/2+current.pop['US']/2+current.pop['AS']/2)
  names(pollen.pool) <- c('L','C','U','A','S')
  # The proportions of genotypes produced by each maternal plant depend on the frequency of each pollen type AMONG COMPATIBLE POLLEN TYPES.
  compatible.with.LL <- c(((L-2)/L)*pollen.pool['L'],
                          ((L-2)/L)*pollen.pool['C'],
                          pollen.pool['U'],
                          pollen.pool['A'],
                          pollen.pool['S'])
  compatible.with.LC <- c(((L-2)/L)*pollen.pool['L'],
                          ((L-2)/L)*pollen.pool['C'],
                          pollen.pool['U'],
                          pollen.pool['A'],
                          pollen.pool['S'])
  compatible.with.LU <- c(((L-1)/(L))*pollen.pool['L'],
                          ((L-1)/(L))*pollen.pool['C'],
                          ((U-1)/(U))*pollen.pool['U'],
                          pollen.pool['A'],
                          pollen.pool['S'])
  if(monster){ # A can be fertilized by S if S is a monster.
    compatible.with.LA <- c(((L-1)/(L))*pollen.pool['L'],
                            ((L-1)/(L))*pollen.pool['C'],
                            pollen.pool['U'],
                            0,
                            pollen.pool['S'])
    }
  else{
  compatible.with.LA <- c(((L-1)/(L))*pollen.pool['L'],
                          ((L-1)/(L))*pollen.pool['C'],
                          pollen.pool['U'],
                          0,
                          0)
  }
  compatible.with.LS <- c(((L-1)/(L))*pollen.pool['L'],
                          0,
                          0,
                          0,
                          0)
  compatible.with.CC <- c(((L-2)/L)*pollen.pool['L'],
                          ((L-2)/L)*pollen.pool['C'],
                          pollen.pool['U'],
                          pollen.pool['A'],
                          pollen.pool['S'])
  compatible.with.CU <- c(((L-1)/(L))*pollen.pool['L'],
                          ((L-1)/(L))*pollen.pool['C'],
                          ((U-1)/(U))*pollen.pool['U'],
                          pollen.pool['A'],
                          pollen.pool['S'])
  if(monster){ # A can be fertilized by a monster S.
    compatible.with.CA <-c(((L-1)/(L))*pollen.pool['L'],
                           ((L-1)/(L))*pollen.pool['C'],
                           pollen.pool['U'],
                           0,
                           pollen.pool['S'])
    }
  else{
  compatible.with.CA <- c(((L-1)/(L))*pollen.pool['L'],
                          ((L-1)/(L))*pollen.pool['C'],
                          pollen.pool['U'],
                          0,
                          0)
  }
  compatible.with.CS <- c(((L-1)/(L))*pollen.pool['L'],
                          0,
                          0,
                          0,
                          0)
  compatible.with.UU <- c(pollen.pool['L'],
                          pollen.pool['C'],
                          ((U-2)/(U))*pollen.pool['U'],
                          pollen.pool['A'],
                          pollen.pool['S'])
  if(monster){ # A can be fertilized by a monster S.
    compatible.with.UA <- c(pollen.pool['L'],
                            pollen.pool['C'],
                            ((U-1)/(U))*pollen.pool['U'],
                            0,
                            pollen.pool['S'])
    }
  else{
  compatible.with.UA <- c(pollen.pool['L'],
                          pollen.pool['C'],
                          ((U-1)/(U))*pollen.pool['U'],
                          0,
                          0)
  }
  compatible.with.US <- c(pollen.pool['L'],
                          0,
                          0,
                          0,
                          0)
  # If there's a monster, there should also needs to be an AS category.
  # AS only exists if the monster exists, so it doesn't need an if statement.
  # Only L pollen can fertilize it.
  compatible.with.AS <- c(pollen.pool['L'],
                          0,
                          0,
                          0,
                          0)
  # Negative or undefined values are the same as zeroes.
  compatible.with.LL[!(compatible.with.LL > 0)] <- 0
  compatible.with.LU[!(compatible.with.LU > 0)] <- 0
  compatible.with.LA[!(compatible.with.LA > 0)] <- 0
  compatible.with.LS[!(compatible.with.LS > 0)] <- 0
  compatible.with.LC[!(compatible.with.LC > 0)] <- 0
  compatible.with.CC[!(compatible.with.CC > 0)] <- 0
  compatible.with.CU[!(compatible.with.CU > 0)] <- 0
  compatible.with.CA[!(compatible.with.CA > 0)] <- 0
  compatible.with.CS[!(compatible.with.CS > 0)] <- 0
  compatible.with.UU[!(compatible.with.UU > 0)] <- 0
  compatible.with.UA[!(compatible.with.UA > 0)] <- 0
  compatible.with.US[!(compatible.with.US > 0)] <- 0
  compatible.with.AS[!(compatible.with.AS > 0)] <- 0
  # Convert to frequencies AMONG COMPATIBLE POLLEN.
  # We want to relativize.
  # However, some vectors add up to zero.
  # For these vectors, just leave them at zero
  if(sum(compatible.with.LL) > 0){compatible.with.LL <- compatible.with.LL/sum(compatible.with.LL)}
  if(sum(compatible.with.LU) > 0){compatible.with.LU <- compatible.with.LU/sum(compatible.with.LU)}
  if(sum(compatible.with.LA) > 0){compatible.with.LA <- compatible.with.LA/sum(compatible.with.LA)}
  if(sum(compatible.with.LS) > 0){compatible.with.LS <- compatible.with.LS/sum(compatible.with.LS)}
  if(sum(compatible.with.LC) > 0){compatible.with.LC <- compatible.with.LC/sum(compatible.with.LC)}
  if(sum(compatible.with.CC) > 0){compatible.with.CC <- compatible.with.CC/sum(compatible.with.CC)}
  if(sum(compatible.with.CU) > 0){compatible.with.CU <- compatible.with.CU/sum(compatible.with.CU)}
  if(sum(compatible.with.CA) > 0){compatible.with.CA <- compatible.with.CA/sum(compatible.with.CA)}
  if(sum(compatible.with.CS) > 0){compatible.with.CS <- compatible.with.CS/sum(compatible.with.CS)}
  if(sum(compatible.with.UU) > 0){compatible.with.UU <- compatible.with.UU/sum(compatible.with.UU)}
  if(sum(compatible.with.UA) > 0){compatible.with.UA <- compatible.with.UA/sum(compatible.with.UA)}
  if(sum(compatible.with.US) > 0){compatible.with.US <- compatible.with.US/sum(compatible.with.US)}
  if(sum(compatible.with.AS) > 0){compatible.with.AS <- compatible.with.AS/sum(compatible.with.AS)}
  # Name the elements of the compatible pollen vectors.
  names(compatible.with.LL) <- names(pollen.pool)
  names(compatible.with.LU) <- names(pollen.pool)
  names(compatible.with.LA) <- names(pollen.pool)
  names(compatible.with.LS) <- names(pollen.pool)
  names(compatible.with.LC) <- names(pollen.pool)
  names(compatible.with.CC) <- names(pollen.pool)
  names(compatible.with.CU) <- names(pollen.pool)
  names(compatible.with.CA) <- names(pollen.pool)
  names(compatible.with.CS) <- names(pollen.pool)
  names(compatible.with.UU) <- names(pollen.pool)
  names(compatible.with.UA) <- names(pollen.pool)
  names(compatible.with.US) <- names(pollen.pool)
  names(compatible.with.AS) <- names(pollen.pool)  
  # Figure out each genotype's contribution.
  LL.contribution <- current.pop['LL'] * c(compatible.with.LL['L'], # LL offspring
                                           compatible.with.LL['C'],# LC offspring
                                           compatible.with.LL['U'], # LU offspring
                                           compatible.with.LL['A'], # LA offspring
                                           compatible.with.LL['S'], # LS offspring
                                           0, # CC offspring
                                           0, # CU offspring
                                           0, # CA offspring
                                           0, # CS offspring
                                           0, # UU offspring
                                           0, # UA offspring
                                           0, # US offspring
                                           0) #AS offspring
  LC.contribution <- current.pop['LC'] * c(compatible.with.LC['L']/2, # LL offspring
                                           compatible.with.LC['L']/2 + compatible.with.LC['C']/2,# LC offspring
                                           compatible.with.LC['U']/2, # LU offspring
                                           compatible.with.LC['A']/2, # LA offspring
                                           compatible.with.LC['S']/2, # LS offspring
                                           compatible.with.LC['C']/2, # CC offspring
                                           compatible.with.LC['U']/2, # CU offspring
                                           compatible.with.LC['A']/2, # CA offspring
                                           compatible.with.LC['S']/2, # CS offspring
                                           0, # UU offspring
                                           0, # UA offspring
                                           0, # US offspring
                                           0) # AS offspring
  LU.contribution <- current.pop['LU'] * c(compatible.with.LU['L']/2, # LL offspring
                                           compatible.with.LU['C']/2, # LC OFFSPRING
                                           compatible.with.LU['U']/2 + compatible.with.LU['L']/2, # LU offspring
                                           compatible.with.LU['A']/2, # LA offspring
                                           compatible.with.LU['S']/2, # LS offspring
                                           0, # CC offspring
                                           compatible.with.LU['C']/2, # CU offspring
                                           0, # CA offspring
                                           0, # CS offspring
                                           compatible.with.LU['U']/2, # UU offspring
                                           compatible.with.LU['A']/2, # UA offspring
                                           compatible.with.LU['S']/2, # US offspring
                                           0) # AS offspring
  LA.contribution <- current.pop['LA'] * c(compatible.with.LA['L']/2, # LL offspring
                                           compatible.with.LA['C']/2, # LC OFFSPRING
                                           compatible.with.LA['U']/2, # LU offspring
                                           compatible.with.LA['L']/2 + compatible.with.LA['A']/2, # LA offspring
                                           compatible.with.LA['S']/2, # LS offspring
                                           0, # CC offspring
                                           0, # CU offspring
                                           compatible.with.LA['C']/2, # CA offspring
                                           0, # CS offspring
                                           0, # UU offspring
                                           compatible.with.LA['U']/2, # UA offspring
                                           0, # US offspring
                                           compatible.with.LA['S']/2) # AS offspring
  LS.contribution <- current.pop['LS'] * c(compatible.with.LS['L']/2, # LL offspring
                                           compatible.with.LS['C']/2, # LC OFFSPRING
                                           compatible.with.LS['U']/2, # LU offspring
                                           compatible.with.LS['A']/2, # LA offspring
                                           compatible.with.LS['S']/2 + compatible.with.LS['L']/2, # LS offspring
                                           0, # CC offspring
                                           0, # CU offspring
                                           0, # CA offspring
                                           compatible.with.LS['C']/2, # CS offspring
                                           0, # UU offspring
                                           0, # UA offspring
                                           compatible.with.LS['U']/2, # US offspring
                                           compatible.with.LS['A']/2) # AS offspring
  CC.contribution <- current.pop['CC'] * c(0, # LL offspring
                                           compatible.with.CC['L'], # LC offspring
                                           0, # LU offspring
                                           0, # LA offspring
                                           0, # LS offspring
                                           compatible.with.CC['C'], # CC offspring
                                           compatible.with.CC['U'], # CU offspring
                                           compatible.with.CC['A'], # CA offspring
                                           compatible.with.CC['S'], # CS offspring
                                           0, # UU offspring 
                                           0, # UA offspring
                                           0, # US offspring
                                           0) # AS offspring
  CU.contribution <- current.pop['CU'] * c(0, # LL offspring
                                           compatible.with.CU['L']/2, # LC offspring
                                           compatible.with.CU['L']/2, # LU offspring
                                           0, # LA offspring
                                           0, # LS offspring
                                           compatible.with.CU['C']/2, # CC offspring
                                           compatible.with.CU['C']/2 + compatible.with.CU['U']/2, # CU offspring
                                           compatible.with.CU['A']/2, # CA offspring
                                           compatible.with.CU['S']/2, # CS offspring
                                           compatible.with.CU['U']/2, # UU offspring 
                                           compatible.with.CU['A']/2, # UA offspring
                                           compatible.with.CU['S']/2, # US offspring
                                           0) # AS offspring
  CA.contribution <- current.pop['CA'] * c(0, # LL offspring
                                           compatible.with.CA['L']/2, # LC offspring
                                           0, # LU offspring
                                           compatible.with.CA['L']/2, # LA offspring
                                           0, # LS offspring
                                           compatible.with.CA['C']/2, # CC offspring
                                           compatible.with.CA['U']/2, # CU offspring
                                           compatible.with.CA['C']/2 + compatible.with.CA['A']/2, # CA offspring
                                           compatible.with.CA['S']/2, # CS offspring
                                           0, # UU offspring 
                                           compatible.with.CA['U']/2, # UA offspring
                                           0, # US offspring
                                           compatible.with.CA['S']/2) # AS offspring
  CS.contribution <- current.pop['CS'] * c(0, # LL offspring
                                           compatible.with.CS['L']/2, # LC offspring
                                           0, # LU offspring
                                           0, # LA offspring
                                           compatible.with.CS['L']/2, # LS offspring
                                           compatible.with.CS['C']/2, # CC offspring
                                           compatible.with.CS['U']/2, # CU offspring
                                           compatible.with.CS['A']/2, # CA offspring
                                           compatible.with.CS['C']/2 + compatible.with.CS['S']/2, # CS offspring
                                           0, # UU offspring 
                                           0, # UA offspring
                                           compatible.with.CS['U']/2, # US offspring
                                           compatible.with.CS['A']/2) # AS offspring
  UU.contribution <- current.pop['UU'] * c(0, # LL offspring
                                           0, # LC OFFSPRING
                                           compatible.with.UU['L'], # LU offspring
                                           0, # LA offspring
                                           0, # LS offspring
                                           0, # CC offspring
                                           compatible.with.UU['C'], # CU offspring
                                           0, # CA offspring
                                           0, # CS offspring
                                           compatible.with.UU['U'], # UU offspring
                                           compatible.with.UU['A'], # UA offspring
                                           compatible.with.UU['S'], # US offspring
                                           0) # AS offspring
  UA.contribution <- current.pop['UA'] * c(0, # LL offspring
                                           0, # LC OFFSPRING
                                           compatible.with.UA['L']/2, # LU offspring
                                           compatible.with.UA['L']/2, # LA offspring
                                           0, # LS offspring
                                           0, # CC offspring
                                           compatible.with.UA['C']/2, # CU offspring
                                           compatible.with.UA['C']/2, # CA offspring
                                           0,# CS offspring
                                           compatible.with.UA['U']/2, # UU offspring
                                           compatible.with.UA['U']/2 + compatible.with.UA['A']/2, # UA offspring
                                           compatible.with.UA['S']/2, # US offspring
                                           compatible.with.UA['S']/2) # AS offspring
  US.contribution <- current.pop['US'] * c(0, # LL offspring
                                           0, # LC offspring
                                           compatible.with.US['L']/2, # LU offspring
                                           0, # LA offspring
                                           compatible.with.US['L']/2, # LS offspring
                                           0, # CC offspring
                                           compatible.with.US['C']/2, # CU offspring
                                           0, # CA offspring
                                           compatible.with.US['C']/2, # CS offspring
                                           compatible.with.US['U']/2, # UU offspring
                                           compatible.with.US['A']/2, # UA offspring
                                           compatible.with.US['S']/2 + compatible.with.US['U']/2, # US offspring
                                           compatible.with.US['A']/2) # AS offspring
  AS.contribution <- current.pop['AS'] * c(0, # LL offspring
                                           0, # LC offspring
                                           0, # LU offspring
                                           compatible.with.AS['L']/2, # LA offspring
                                           compatible.with.AS['L']/2, # LS offspring
                                           0, # CC offspring
                                           0, # CU offspring
                                           compatible.with.AS['C']/2, # CA offspring
                                           compatible.with.AS['C']/2, # CS offspring
                                           0, # UU offspring
                                           compatible.with.AS['U']/2, # UA offspring
                                           compatible.with.AS['U']/2, # US offspring
                                           compatible.with.AS['A']/2 + compatible.with.AS['S']/2) # AS offspring
  # Add up the contributions into a single vector.
  new.pop <- LL.contribution +
    LC.contribution +
    LU.contribution +
    LA.contribution +
    LS.contribution +
    CC.contribution +
    CU.contribution +
    CA.contribution +
    CS.contribution +
    UU.contribution +
    UA.contribution +
    US.contribution +
    AS.contribution
  # Name the elements of the vector.
  names(new.pop) <- names(current.pop)
  return(new.pop)
}

# The function track.diplos tracks the diploid genotypes through time for a specified number of generations.
# The output is a matrix of genotype frequencies with one row per generation.
# This can later be converted to haplotype frequencies through time.

track.diplos <- function(pop, last.gen, monster = F){
  # First create the matrix.
  out <- matrix(nrow = last.gen, ncol = length(pop))
  colnames(out) <- names(pop)
  out[1,] <- pop[]
  for(i in 2:last.gen){
    out[i,] <- one.generation(out[i-1,], monster)
    # Debug the last step.
    # if(i == last.gen - 1){debug(one.generation)}
    # if(i == last.gen){undebug(one.generation)}
  }
  return(out)
}

# The function rename will put names back in.
rename <- function(x,nom){
  names(x) <- nom
  return(x)
}

# The function diplos.to.haplos converts a vector of diploid genotypes into a vector of haplotype frequencies.
diplos.to.haplos <- function(pop){
  pL <- (pop['LL'])+(pop['LC']/2)+(pop['LU']/2)+(pop['LS']/2)+(pop['LA']/2)
  pC <- (pop['LC']/2)+(pop['CC'])+(pop['CU']/2)+(pop['CA']/2)+(pop['CS']/2)
  pU <- (pop['LU']/2)+(pop['CU']/2)+(pop['US']/2)+pop['UU']+(pop['UA']/2)
  pA <- (pop['LA']/2)+(pop['CA']/2)+(pop['UA']/2)
  pS <- (pop['LS']/2)+(pop['CS']/2)+(pop['US']/2)
  out <- c(pL,pC,pU,pA,pS)
  names(out) <- c('L','C','U','A','S')
  return(out)
}

# The function get.mean.pollen.fitness counts the fraction of pollen grains expected to land on compatible plants.
get.mean.pollen.fitness <- function(pop){
  current.pop <- pop
  # Create a pollen pool.
  pollen.pool <- c(current.pop['LL']+current.pop['LC']/2+current.pop['LU']/2+current.pop['LA']/2+current.pop['LS']/2,
                   current.pop['LC']/2+current.pop['CC']+current.pop['CU']/2+current.pop['CA']/2+current.pop['CS']/2,
                   current.pop['LU']/2+current.pop['CU']/2+current.pop['UU']+current.pop['UA']/2+current.pop['US']/2,
                   current.pop['LA']/2+current.pop['CA']/2+current.pop['UA']/2,
                   current.pop['LS']/2+current.pop['CS']/2+current.pop['US']/2)
  names(pollen.pool) <- c('L','C','U','A','S')
  # The fraction rejected depends on the rejecting phenotypes.
  compatible.with.LL <- c(((L-2)/L)*pollen.pool['L'],
                          ((L-2)/L)*pollen.pool['C'],
                          pollen.pool['U'],
                          pollen.pool['A'],
                          pollen.pool['S'])
  compatible.with.LC <- c(((L-2)/L)*pollen.pool['L'],
                          ((L-2)/L)*pollen.pool['C'],
                          pollen.pool['U'],
                          pollen.pool['A'],
                          pollen.pool['S'])
  compatible.with.LU <- c(((L-1)/(L))*pollen.pool['L'],
                          ((L-1)/(L))*pollen.pool['C'],
                          ((U-1)/(U))*pollen.pool['U'],
                          pollen.pool['A'],
                          pollen.pool['S'])
  compatible.with.LA <- c(((L-1)/(L))*pollen.pool['L'],
                          ((L-1)/(L))*pollen.pool['C'],
                          pollen.pool['U'],
                          0,
                          0)
  compatible.with.LS <- c(((L-1)/(L))*pollen.pool['L'],
                          0,
                          0,
                          0,
                          0)
  compatible.with.CC <- c(((L-2)/L)*pollen.pool['L'],
                          ((L-2)/L)*pollen.pool['C'],
                          pollen.pool['U'],
                          pollen.pool['A'],
                          pollen.pool['S'])
  compatible.with.CU <- c(((L-1)/(L))*pollen.pool['L'],
                          ((L-1)/(L))*pollen.pool['C'],
                          ((U-1)/(U))*pollen.pool['U'],
                          pollen.pool['A'],
                          pollen.pool['S'])
  compatible.with.CA <- c(((L-1)/(L))*pollen.pool['L'],
                          ((L-1)/(L))*pollen.pool['C'],
                          pollen.pool['U'],
                          0,
                          0)
  compatible.with.CS <- c(((L-1)/(L))*pollen.pool['L'],
                          0,
                          0,
                          0,
                          0)
  compatible.with.UU <- c(pollen.pool['L'],
                          pollen.pool['C'],
                          ((U-2)/(U))*pollen.pool['U'],
                          pollen.pool['A'],
                          pollen.pool['S'])
  compatible.with.UA <- c(pollen.pool['L'],
                          pollen.pool['C'],
                          ((U-1)/(U))*pollen.pool['U'],
                          0,
                          0)
  compatible.with.US <- c(pollen.pool['L'],
                          0,
                          0,
                          0,
                          0)
  # Negative or undefined values are the same as zeroes.
  compatible.with.LL[!(compatible.with.LL > 0)] <- 0
  compatible.with.LC[!(compatible.with.LC > 0)] <- 0
  compatible.with.LU[!(compatible.with.LU > 0)] <- 0
  compatible.with.LA[!(compatible.with.LA > 0)] <- 0
  compatible.with.LS[!(compatible.with.LS > 0)] <- 0
  compatible.with.CC[!(compatible.with.CC > 0)] <- 0
  compatible.with.CU[!(compatible.with.CU > 0)] <- 0
  compatible.with.CA[!(compatible.with.CA > 0)] <- 0
  compatible.with.CS[!(compatible.with.CS > 0)] <- 0
  compatible.with.UU[!(compatible.with.UU > 0)] <- 0
  compatible.with.UA[!(compatible.with.UA > 0)] <- 0
  compatible.with.US[!(compatible.with.US > 0)] <- 0
  incompatible.with.LL <- 1 - compatible.with.LL
  incompatible.with.LC <- 1 - compatible.with.LC
  incompatible.with.LU <- 1 - compatible.with.LU
  incompatible.with.LA <- 1 - compatible.with.LA
  incompatible.with.LS <- 1 - compatible.with.LS
  incompatible.with.CC <- 1 - compatible.with.CC
  incompatible.with.CU <- 1 - compatible.with.CU
  incompatible.with.CA <- 1 - compatible.with.CA
  incompatible.with.CS <- 1 - compatible.with.CS
  incompatible.with.UU <- 1 - compatible.with.UU
  incompatible.with.UA <- 1 - compatible.with.UA
  incompatible.with.US <- 1 - compatible.with.US
  # Now count up how often incompatible pollinations are expected.
  rejected <- pollen.pool*incompatible.with.LL * current.pop['LL'] +
    pollen.pool*incompatible.with.LC * current.pop['LC'] +
    pollen.pool*incompatible.with.LU * current.pop['LU'] +
    pollen.pool*incompatible.with.LA * current.pop['LA'] +
    pollen.pool*incompatible.with.LS * current.pop['LS'] +
    pollen.pool*incompatible.with.CC * current.pop['CC'] +
    pollen.pool*incompatible.with.CU * current.pop['CU'] +
    pollen.pool*incompatible.with.CA * current.pop['CA'] +
    pollen.pool*incompatible.with.CS * current.pop['CS'] +
    pollen.pool*incompatible.with.UU * current.pop['UU'] +
    pollen.pool*incompatible.with.UA * current.pop['UA'] +
    pollen.pool*incompatible.with.US * current.pop['US']
  names(rejected) <- NULL
  return(1- sum(rejected))
}

# Now I need to find the probability that at least one gene conversion survives.
# I need to multiply a vector of frequencies of LU by a vector of fixation probabilities for each generation.
# The vector of LU frequencies per generation is easy to extract.
# The survival probabilities depend on the genotype frequencies at that time.
# Therefore, the survival probabilities will need to be extracted from the population history.
# I can plug in the allele frequencies to get the expected number of offspring copies per copy of the converted haplotype.
# If I assume Poisson distributed offspring per copy, I have to add up all infinite possible numbers of offspring copies.
# The best that a normal allele could do would be to be the ancestor of all alleles in the next generation.
# For an S-allele, the best it could do would be to be the ancestor of half the alleles in the next generation.
# If I assume the population is at carrying capacity, then I just have to add up all possibilities from 0 offspring to N offspring.
# However, there remains a mathematical difficulty.
# I need to solve for the fixation probability at time t.
# Otto and Whitlock manage to do this by approximating a conversion factor between P_t and P_t+1.
# Assuming weak positive selection, the conversion factor was a ratio of population sizes.
# Whereas population size changes in their model, the selection coefficient changes in mine.

# The function generate.supply generates a vector of the number of potentially rescued gene convertants appear each generation.
generate.supply <- function(convertible.traj, N, conversion.rate){
  return(convertible.traj * N * conversion.rate)
}

# The function generate.binomial.supply is like generate.supply, but creates a distribution of convertant numbers instead of
# just the mean.
# generate.binomial.supply <- function(convertible.traj, N, conversion.prob){
#   # Create a probability vector of the number of gene convertants that could have occurred in a generation.
#   convertant_dist <- dbinom(seq(from = 1, to = ceiling(convertible.traj*N), by = 1), prob = convertion.prob)
#   names(convertant_dist) <- 1:ceiling(convertible.traj*N)
#   return(convertant_dist)
# }

# The function determine.s calculates survival probability using the selection coefficient (Haldane's definition:
# expected offspring in excess of the average) of a hypothetical gene convertant.
# The mean absolute fitness of a haplotype equals one in a population at carrying capacity.
# So all we need to do is to combine frequency changes with population size to calculate the absolute fitness of gene convertant.
# Then, just subtract one to get Haldane's s.
# The function will do this every generation given a trajectory of genotype frequencies.
# Since there are several kinds of haplotype that can be doomed, the kind must be specified in the arguments.
# It could be a U haplotype that did not acquire the new key in the invasion phase.
# It could be the A haplotype acquiring its descendant's key.
# It could be a C haplotype if equilibrium was not reached in the invasion phase.
# It could even be the S haplotype if its ancestor acquired the descendant's key first.
# By default, it is one of the U haplotypes.
# Once s is calculated, the function multiplies by two to get Haldane's approximation for the survival probability.

determine.survival.prob <- function(traj, N, doomed.kind = 'U'){
  # Initialize a vector for the per-generation survival probabilities.
  out <- vector(length = nrow(traj))
  for(i in 1:nrow(traj)){
    current.pop <- traj[i,]
    # Create a pollen pool.
    pollen.pool <- c(current.pop['LL']+current.pop['LC']/2+current.pop['LU']/2+current.pop['LA']/2+current.pop['LS']/2,
                     current.pop['LC']/2+current.pop['CC']+current.pop['CU']/2+current.pop['CA']/2+current.pop['CS']/2,
                     current.pop['LU']/2+current.pop['CU']/2+current.pop['UU']+current.pop['UA']/2+current.pop['US']/2,
                     current.pop['LA']/2+current.pop['CA']/2+current.pop['UA']/2,
                     current.pop['LS']/2+current.pop['CS']/2+current.pop['US']/2)
    names(pollen.pool) <- c('L','C','U','A','S')
    # The fraction rejected depends on the rejecting phenotypes.
    compatible.with.LL <- c(((L-2)/L)*pollen.pool['L'],
                            ((L-2)/L)*pollen.pool['C'],
                            pollen.pool['U'],
                            pollen.pool['A'],
                            pollen.pool['S'])
    compatible.with.LC <- c(((L-2)/L)*pollen.pool['L'],
                            ((L-2)/L)*pollen.pool['C'],
                            pollen.pool['U'],
                            pollen.pool['A'],
                            pollen.pool['S'])
    compatible.with.LU <- c(((L-1)/(L))*pollen.pool['L'],
                            ((L-1)/(L))*pollen.pool['C'],
                            ((U-1)/(U))*pollen.pool['U'],
                            pollen.pool['A'],
                            pollen.pool['S'])
    compatible.with.LA <- c(((L-1)/(L))*pollen.pool['L'],
                            ((L-1)/(L))*pollen.pool['C'],
                            pollen.pool['U'],
                            0,
                            0)
    compatible.with.LS <- c(((L-1)/(L))*pollen.pool['L'],
                            0,
                            0,
                            0,
                            0)
    compatible.with.CC <- c(((L-2)/L)*pollen.pool['L'],
                            ((L-2)/L)*pollen.pool['C'],
                            pollen.pool['U'],
                            pollen.pool['A'],
                            pollen.pool['S'])
    compatible.with.CU <- c(((L-1)/(L))*pollen.pool['L'],
                            ((L-1)/(L))*pollen.pool['C'],
                            ((U-1)/(U))*pollen.pool['U'],
                            pollen.pool['A'],
                            pollen.pool['S'])
    compatible.with.CA <- c(((L-1)/(L))*pollen.pool['L'],
                            ((L-1)/(L))*pollen.pool['C'],
                            pollen.pool['U'],
                            0,
                            0)
    compatible.with.CS <- c(((L-1)/(L))*pollen.pool['L'],
                            0,
                            0,
                            0,
                            0)
    compatible.with.UU <- c(pollen.pool['L'],
                            pollen.pool['C'],
                            ((U-2)/(U))*pollen.pool['U'],
                            pollen.pool['A'],
                            pollen.pool['S'])
    compatible.with.UA <- c(pollen.pool['L'],
                            pollen.pool['C'],
                            ((U-1)/(U))*pollen.pool['U'],
                            0,
                            0)
    compatible.with.US <- c(pollen.pool['L'],
                            0,
                            0,
                            0,
                            0)
    # Negative or undefined values are the same as zeroes.
    compatible.with.LL[!(compatible.with.LL > 0)] <- 0
    compatible.with.LC[!(compatible.with.LC > 0)] <- 0
    compatible.with.LU[!(compatible.with.LU > 0)] <- 0
    compatible.with.LA[!(compatible.with.LA > 0)] <- 0
    compatible.with.LS[!(compatible.with.LS > 0)] <- 0
    compatible.with.CC[!(compatible.with.CC > 0)] <- 0
    compatible.with.CU[!(compatible.with.CU > 0)] <- 0
    compatible.with.CA[!(compatible.with.CA > 0)] <- 0
    compatible.with.CS[!(compatible.with.CS > 0)] <- 0
    compatible.with.UU[!(compatible.with.UU > 0)] <- 0
    compatible.with.UA[!(compatible.with.UA > 0)] <- 0
    compatible.with.US[!(compatible.with.US > 0)] <- 0
    incompatible.with.LL <- 1 - compatible.with.LL
    incompatible.with.LC <- 1 - compatible.with.LC
    incompatible.with.LU <- 1 - compatible.with.LU
    incompatible.with.LA <- 1 - compatible.with.LA
    incompatible.with.LS <- 1 - compatible.with.LS
    incompatible.with.CC <- 1 - compatible.with.CC
    incompatible.with.CU <- 1 - compatible.with.CU
    incompatible.with.CA <- 1 - compatible.with.CA
    incompatible.with.CS <- 1 - compatible.with.CS
    incompatible.with.UU <- 1 - compatible.with.UU
    incompatible.with.UA <- 1 - compatible.with.UA
    incompatible.with.US <- 1 - compatible.with.US
    # Now count up how often incompatible pollinations are expected.
    rejected <- pollen.pool*incompatible.with.LL * current.pop['LL'] +
      pollen.pool*incompatible.with.LC * current.pop['LC'] +
      pollen.pool*incompatible.with.LU * current.pop['LU'] +
      pollen.pool*incompatible.with.LA * current.pop['LA'] +
      pollen.pool*incompatible.with.LS * current.pop['LS'] +
      pollen.pool*incompatible.with.CC * current.pop['CC'] +
      pollen.pool*incompatible.with.CU * current.pop['CU'] +
      pollen.pool*incompatible.with.CA * current.pop['CA'] +
      pollen.pool*incompatible.with.CS * current.pop['CS'] +
      pollen.pool*incompatible.with.UU * current.pop['UU'] +
      pollen.pool*incompatible.with.UA * current.pop['UA'] +
      pollen.pool*incompatible.with.US * current.pop['US']
    names(rejected) <- NULL
    mean.fit <- 0.5 + 0.5*(1 - sum(rejected))
    # The mean fitness is now known.
    # The fitness of the gene convertant can be calculated from the following formula.
    # Assuming that the gene convertant is a variant of U,
    if(doomed.kind == 'U'){
      convert.fit <- 0.5 + 0.5*(current.pop['LL']/sum(compatible.with.LL) +
                                  current.pop['LC']/sum(compatible.with.LC) +
                                  ((U - 1)/U) * current.pop['LU']/sum(compatible.with.LU) +
                                  current.pop['LA']/sum(compatible.with.LA) +
                                  current.pop['LS']/sum(compatible.with.LS) +
                                  current.pop['CC']/sum(compatible.with.CC) +  
                                  ((U - 1)/U) * current.pop['CU']/sum(compatible.with.CU) +
                                  current.pop['CA']/sum(compatible.with.CA) +
                                  current.pop['CS']/sum(compatible.with.CS) +
                                  ((U - 2)/U) * current.pop['UU']/sum(compatible.with.UU) +
                                  ((U - 1)/U) * current.pop['UA']/sum(compatible.with.UA) +
                                  ((U - 1)/U) * current.pop['US']/sum(compatible.with.US))
    }
    # Assuming that the gene convertant is a variant of C,
    if(doomed.kind == 'C'){
      convert.fit <- 0.5 + 0.5*(((L - 2)/L) * current.pop['LL']/sum(compatible.with.LL) +
                                  ((L - 2)/L) * current.pop['LC']/sum(compatible.with.LC) +
                                  ((L - 1)/L) * current.pop['LU']/sum(compatible.with.LU) +
                                  ((L - 1)/L) * current.pop['LA']/sum(compatible.with.LA) +
                                  ((L - 1)/L) * current.pop['LS']/sum(compatible.with.LS) +
                                  ((L - 2)/L) * current.pop['CC']/sum(compatible.with.CC) +
                                  ((L - 1)/L) * current.pop['CU']/sum(compatible.with.CU) +
                                  ((L - 1)/L) * current.pop['CA']/sum(compatible.with.CA) +
                                  ((L - 1)/L) * current.pop['CS']/sum(compatible.with.CS) +
                                  current.pop['UU']/sum(compatible.with.UU) +
                                  current.pop['UA']/sum(compatible.with.UA) +
                                  current.pop['US']/sum(compatible.with.US))
    }
    # Assuming that the gene convertant is a variant of A,
    if(doomed.kind == 'A'){
      convert.fit <- 0.5 + 0.5*(current.pop['LL']/sum(compatible.with.LL) +
                                  current.pop['LC']/sum(compatible.with.LC) +
                                  current.pop['LU']/sum(compatible.with.LU) +
                                  0 * current.pop['LA']/sum(compatible.with.LA) +
                                  current.pop['LS']/sum(compatible.with.LS) +
                                  current.pop['CC']/sum(compatible.with.CC) +
                                  current.pop['CU']/sum(compatible.with.CU) +
                                  0 * current.pop['CA']/sum(compatible.with.CA) +
                                  current.pop['CS']/sum(compatible.with.CS) +
                                  current.pop['UU']/sum(compatible.with.UU) +
                                  0 * current.pop['UA']/sum(compatible.with.UA) +
                                  current.pop['US']/sum(compatible.with.US))
    }
    # Assuming that the gene convertant is a variant of S,
    if(doomed.kind == 'S'){
      convert.fit <- 0.5 + 0.5*(current.pop['LL']/sum(compatible.with.LL) +
                                  current.pop['LC']/sum(compatible.with.LC) +
                                  current.pop['LU']/sum(compatible.with.LU) +
                                  current.pop['LA']/sum(compatible.with.LA) +
                                  0 * current.pop['LS']/sum(compatible.with.LS) +
                                  current.pop['CC']/sum(compatible.with.CC) +
                                  current.pop['CU']/sum(compatible.with.CU) +
                                  current.pop['CA']/sum(compatible.with.CA) +
                                  0 * current.pop['CS']/sum(compatible.with.CS) +
                                  current.pop['UU']/sum(compatible.with.UU) +
                                  current.pop['UA']/sum(compatible.with.UA) +
                                  0 * current.pop['US']/sum(compatible.with.US))
    }
    # Now calculate Haldane's selection coefficient.
    s <- (convert.fit - mean.fit)/mean.fit
    # Finally, multiply by two to get the survival probability.
    out[i] <- 2 * s
  }
  return(out)
}

# Given the supply of gene convertants and their probabilities of survival, we can calculate the probability that no convertants
# both arise and survive.
# Currently, this is the probability of rescue conditional on exactly the expected number of convertants arising.
# This could be improved by making the number of gene conversions a distribution rather than a single rate.
# One possibility would be to have a per-individual conversion probability, and make the number that is actually converted binomial.
# Once we calculate the probability that no convertants are rescued, we can take the complement to get the probability of rescue.
# This is the probability of rescue for a single haplotype.
# The process can be repeated for each doomed haplotype and multiplied, or the probability can just be raised to the appropriate
# power if all trajectories are identical, as in the case of only doomed U haplotypes.
# Its inputs are a supply vector and a vector of survival probabilities.

rescue.prob <- function(supply.vec, survive.vec){
  # Convert the survival probability vector into a vector of loss probabilities.
  loss.vec <- 1 - survive.vec
  # Each generation, raise the probability of loss to a power equal to the expected number of convertants that generation.
  # Multiply these to get the total probability of no survivors.
  no.survivors.prob <- prod(loss.vec ^ supply.vec)
  # Now return the complement.
  return(1 - no.survivors.prob)
}

# The function series.solver approximates a fixation probability from a selection coefficient using Haldane's series approximation.
# When the selection coefficient is small, terms after the first in the series can be ignored.
# This is for when the selection coefficient is high.
# It will plug in a sequence of possible fixation probabilities specified by the user, and will output the one that results in the
# selection coefficient closest to its actual value.
# The user also specifies how many terms to use in the infinite series.

series.solver <- function(sel, guesses, terms){
  guess.s <- vector(length = length(guesses))
  guess.series <- vector(length = terms)
  for(i in 1:length(guesses)){
    for(j in 1:terms){
      guess.series[j] <- (guesses[i] ^ j)/(j + 1)
    }
    guess.s[i] <- sum(guess.series)
  }
  best.guess <- which(abs(guess.s - sel) == min(abs(guess.s - sel)))
  return(guesses[best.guess])
}

initialize <- function(L, U, luck_frac, spite_frac){
  # There are initially n = U + L + 1 RNases in the population.
  n <<- U + L + 1
  
  # Each RNase starts at frequency 1/n, but the SLF genotypes are determined by luck_frac
  pL <- (1/n) * luck_frac * L # Multiply by the number of RNases contained in this class because we're lumping them together.
  pC <- (1/n) * (1 - luck_frac) * L
  pU <- (1/n) * U
  pA <- (1/n) * (1 - spite_frac)
  pS <- (1/n) * spite_frac
  
  # Define the initial diploid genotype frequencies.
  # We assume that, conditional on the two haplotypes being compatible, the diploid genotype is independent of which particular compatible alleles are together.
  # This is justified because, immediately before the spiteful RNase appeared, all haplotypes were fully functional.
  # Therefore, no haplotypes would be particularly good at fertilizing particular other haplotypes.
  
  # It's the probability of randomly sampling both of them, given that RNase homozygotes are forbidden, times the number of combinations
  # that are classified as the event.
  
  # We must first find the probability of a successful fertilization and condition on that.
  prob_success <- 1 - 
    (pL ^ 2) * 1/L -
    pL * pC * 2 * 1/L -
    (pC ^ 2) * 1/L -
    (pU ^ 2) * 1/U -
    pA ^ 2 -
    pA * pS * 2 -
    pS ^ 2
  
  pLL <- ((pL ^ 2) * (L - 1)/L) / prob_success
  pLC <- (pL * pC * 2 * (L - 1)/L) / prob_success
  pLU <- (pL * pU * 2) / prob_success
  pLA <- (pL * pA * 2) / prob_success
  pLS <- (pL * pS * 2) / prob_success
  pCC <- ((pC ^ 2) * (L - 1)/L) / prob_success
  pCU <- (pC * pU * 2) / prob_success
  pCA <- (pC * pA * 2) / prob_success
  pCS <- (pC * pS * 2) / prob_success
  pUU <- ((pU ^ 2) * (U - 1)/U) / prob_success
  pUA <- (pU * pA * 2) / prob_success
  pUS <- (pU * pS * 2) / prob_success
  # pAA <- (0 * pA ^ 2) / prob_success
  pAS <- (0 * pA * pS * 2) / prob_success # AS is possible after the monster arises, but initially exists at frequency zero.
  # pSS <- (0 * pS ^ 2) / prob_success
  
  # Put the diploid genotype frequencies in a vector.
  
  population <<- c(pLL,
                   pLC,
                   pLU,
                   pLA,
                   pLS,
                   pCC,
                   pCU,
                   pCA,
                   pCS,
                   pUU,
                   pUA,
                   pUS,
                   # pAA,
                   pAS
                   # pSS
  )
  
  names(population) <<- c('LL',
                          'LC',
                          'LU',
                          'LA',
                          'LS',
                          'CC',
                          'CU',
                          'CA',
                          'CS',
                          'UU',
                          'UA',
                          'US',
                          # 'AA',
                          'AS'
                          # 'SS'
  )
}

# Initialize given that there can be an arbitrary number of genotypes.
# In fact, take the list of genotypes as an argument.
# Since the only difference between U and C is whether there is a corresponding L, make Us into Cs where the Ls have frequency 0.
# We need to recognize that the existence of a U implies the possibility of a corresponding L, even if it doesn't currently exist.
# Ls that correspond to Cs get positive frequencies, whereas Ls that correspond to Us get frequencies of zero.
sim.initialize <- function(genos, luck_frac, spite_frac){
  # We need a vector of all the haplotypes for several reasons.
  haplos <- unique(as.vector(genos))
  # Count the number of Ls and Us.
  # There should be as many EXTANT Ls as there are Cs, by definition.
  L <- sum(grepl('C', haplos))
  U <- sum(grepl('U', haplos))
  # There are initially n = U + L + 1 RNases in the population.
  n <- U + L + 1
  # Each CLASS OF RNase starts at frequency 1/n, but the SLF genotypes are determined by luck_frac
  pL <- (1/n) * luck_frac * L # Multiply by the number of RNases contained in this class because we're lumping them together.
  pC <- (1/n) * (1 - luck_frac) * L
  pU <- (1/n) * U
  pA <- (1/n) * (1 - spite_frac)
  pS <- (1/n) * spite_frac
  
  # Define the initial diploid genotype frequencies.
  # We assume that, conditional on the two haplotypes being compatible, the diploid genotype is independent of which particular compatible alleles are together.
  # This is justified because, immediately before the spiteful RNase appeared, all haplotypes were fully functional.
  # Therefore, no haplotypes would be particularly good at fertilizing particular other haplotypes.
  
  # It's the probability of randomly sampling both of them, given that RNase homozygotes are forbidden, times the number of combinations
  # that are classified as the event.
  
  # We must first find the probability of a successful fertilization and condition on that.
  prob_success <- 1 - 
    (pL ^ 2) * 1/L -
    pL * pC * 2 * 1/L -
    (pC ^ 2) * 1/L -
    (pU ^ 2) * 1/U -
    pA ^ 2 -
    pA * pS * 2 -
    pS ^ 2
  
  pLL <- ((pL ^ 2) * (L - 1)/L) / prob_success
  pLC <- (pL * pC * 2 * (L - 1)/L) / prob_success
  pLU <- (pL * pU * 2) / prob_success
  pLA <- (pL * pA * 2) / prob_success
  pLS <- (pL * pS * 2) / prob_success
  pCC <- ((pC ^ 2) * (L - 1)/L) / prob_success
  pCU <- (pC * pU * 2) / prob_success
  pCA <- (pC * pA * 2) / prob_success
  pCS <- (pC * pS * 2) / prob_success
  pUU <- ((pU ^ 2) * (U - 1)/U) / prob_success
  pUA <- (pU * pA * 2) / prob_success
  pUS <- (pU * pS * 2) / prob_success
  # pAA <- (0 * pA ^ 2) / prob_success
  # pAS <- (0 * pA * pS * 2) / prob_success
  # pSS <- (0 * pS ^ 2) / prob_success

  # We now know the expected frequencies of each CLASS OF genotype.
  # What we need to do is divide up these frequencies by the actual genotypes making up those classes.
  # To do that, we need to identify which genotypes fall into which classes.
  LL <- grepl('L', genos[,1]) & grepl('L', genos[,2]) | grepl('L', genos[,2]) & grepl('L', genos[,1])
  LC <- grepl('L', genos[,1]) & grepl('C', genos[,2]) | grepl('L', genos[,2]) & grepl('C', genos[,1])
  LU <- grepl('L', genos[,1]) & grepl('U', genos[,2]) | grepl('L', genos[,2]) & grepl('U', genos[,1])
  LA <- grepl('L', genos[,1]) & grepl('A', genos[,2]) | grepl('L', genos[,2]) & grepl('A', genos[,1])
  LS <- grepl('L', genos[,1]) & grepl('S', genos[,2]) | grepl('L', genos[,2]) & grepl('S', genos[,1])
  CC <- grepl('C', genos[,1]) & grepl('C', genos[,2]) | grepl('C', genos[,2]) & grepl('C', genos[,1])
  CU <- grepl('C', genos[,1]) & grepl('U', genos[,2]) | grepl('C', genos[,2]) & grepl('U', genos[,1])
  CA <- grepl('C', genos[,1]) & grepl('A', genos[,2]) | grepl('C', genos[,2]) & grepl('A', genos[,1])
  CS <- grepl('C', genos[,1]) & grepl('S', genos[,2]) | grepl('C', genos[,2]) & grepl('S', genos[,1])
  UU <- grepl('U', genos[,1]) & grepl('U', genos[,2]) | grepl('U', genos[,2]) & grepl('U', genos[,1])
  UA <- grepl('U', genos[,1]) & grepl('A', genos[,2]) | grepl('U', genos[,2]) & grepl('A', genos[,1])
  US <- grepl('U', genos[,1]) & grepl('S', genos[,2]) | grepl('U', genos[,2]) & grepl('S', genos[,1])

  # Remove non-extant Ls.
  non_extant_Ls <- paste('L', (L+1):(L+U), sep = '')
  non_extant_genos <- genos[,1] %in% non_extant_Ls | genos[,2] %in% non_extant_Ls
  LL[non_extant_genos] <- F
  LC[non_extant_genos] <- F
  LU[non_extant_genos] <- F
  LA[non_extant_genos] <- F
  LS[non_extant_genos] <- F
  # Now that we know which things are which, we can make the frequency of every member of a class equal to all other members of it.
  
  # Put the diploid genotype frequencies in a vector.
  
  population <- vector(length = nrow(genos))
  names(population) <- paste(genos[,1], genos[,2], sep = '_')
  population[LL] <- pLL/sum(LL)
  population[LC] <- pLC/sum(LC)
  population[LU] <- pLU/sum(LU)
  population[LA] <- pLA/sum(LA)
  population[LS] <- pLS/sum(LS)
  population[CC] <- pCC/sum(CC)
  population[CU] <- pCU/sum(CU)
  population[CA] <- pCA/sum(CA)
  population[CS] <- pCS/sum(CS)
  population[UU] <- pUU/sum(UU)
  population[UA] <- pUA/sum(UA)
  population[US] <- pUS/sum(US)
  population[non_extant_genos] <- 0
  return(population)
}

############################
# FUNCTIONS FOR SIMULATION #
############################

# Generate all genotypic combinations.
library(gtools)
generate.genos <- function(L, U, monster = F){
  # First, enumerate the haplotypes.
  haplos <- c(paste('L', 1:(L+U), sep = ''), # L haplotypes, extant and non-
              paste('C', 1:L, sep = ''), # C haplotypes
              paste('U', (L+1):(L+U), sep = ''), # U haplotypes
              'A',
              'S'
  )
  # Second, generate all combinations.
  raw <- combinations(length(haplos), 2, haplos)
  # Then, delete the illegitimate ones.
  legitimate <- rep(T, nrow(raw))
  # The easiest-to-spot illegitimate ones are the homozygotes.
  # These are always illegitimate.
  # RNase homozygotes with different SLF complements are harder.
  # grepping is dangerous because, for example, grepping "L1" will give you not only L1, but L10, L11, etc.
  # The safest way to do it would be to use RNase IDs again.
  # But this time, it will be for the RNases in raw instead of haplos.
  rnase_id <- gsub('U', '', raw)
  rnase_id <- gsub('L', '', rnase_id)
  # REMEMBER TO REMOVE Cs!
  rnase_id <- gsub('C', '', rnase_id)
  legitimate[rnase_id[,1] == rnase_id[,2]] <- F
  # AS heterozygotes are also easy to spot.
  # However, they are only illegitimate if the monster RNase hasn't shown up.
  if(monster){}
  else{
    legitimate[(raw[,1] == 'A' & raw[,2] == 'S' | raw[,1] == 'S' & raw[,2] == 'A')] <- F
  }
  return(raw[legitimate,])
}

# Generate a pollen production matrix.
generate.pollen.production <- function(genos){
  # The rows of the matrix will be the diploid genotype.
  # The columns will be that genotype's contribution to each type of pollen.
  # This will be 0.5 for things it can produce and 0 for things it can't.
  pollen_names <- unique(as.vector(genos))
  # Find which pollen haplotypes are produced by the left haplotype in the pollen parent.
  produced_by_left <- matrix(genos[,1] == rep(pollen_names, each = nrow(genos)),
                             nrow = nrow(genos),
                             ncol = length(pollen_names))
  produced_by_right <- matrix(genos[,2] == rep(pollen_names, each = nrow(genos)),
                              nrow = nrow(genos),
                              ncol = length(pollen_names))
  production <- 0.5 * (produced_by_left | produced_by_right)
  rownames(production) <- paste(genos[,1], genos[,2], sep = '_')
  colnames(production) <- pollen_names
  return(production)
}

# Produce pollen given genotype frequencies and a pollen production matrix.
produce.pollen <- function(geno.dist, pollen.production.matrix){
  return(colSums(geno.dist[rownames(pollen.production.matrix)] * pollen.production.matrix))
}

# Generate a compatibility matrix given the number of lucky haplotypes (L) and the number of unlucky haplotypes (U).
generate.haplo.compatibility <- function(genos, monster = F){
  haplos <- unique(as.vector(genos))
  out <- matrix(nrow = length(haplos), ncol = length(haplos))
  rownames(out) <- haplos
  colnames(out) <- haplos
  # The rules are:
  # Identical haplotypes are incompatible.
  # Haplotypes sharing a number are incompatible.
  # Delete any Ls or Us, then take whatever's left as the ID.
  rnase_id <- gsub('L', '', haplos)
  rnase_id <- gsub('U', '', rnase_id)
  # REMEMBER TO REMOVE Cs!
  rnase_id <- gsub('C', '', rnase_id)
  # This will be a letter if it's a one-character haplotype or a number if it's a multi-character haplotype.
  # Only L pollen is compatible with S.
  # S pollen is incompatible with A UNLESS monster is TRUE.
  # Otherwise, they're compatible.
  for(i in 1:nrow(out)){
    for(j in 1:ncol(out)){
      if(monster){
        incompatible <- rownames(out)[i] == rownames(out)[j] |   # Identical haplotypes are incompatible.
          rnase_id[i] == rnase_id[j] | # Haplotypes sharing a number are incompatible.
          grepl('S', colnames(out)[j]) & !(grepl('L', rownames(out)[i])) # Only L pollen is compatible with S.
        }
      else{
      incompatible <- rownames(out)[i] == rownames(out)[j] |   # Identical haplotypes are incompatible.
                      rnase_id[i] == rnase_id[j] | # Haplotypes sharing a number are incompatible.
                      grepl('S', colnames(out)[j]) & !(grepl('L', rownames(out)[i])) | # Only L pollen is compatible with S.
                      grepl('S', rownames(out)[i]) & grepl('A', colnames(out)[j])# S pollen is incompatible with A.
      }
      out[i,j] <- !incompatible # Otherwise, they're compatible.
    }
  }
  return(out)
}

# Use the incompatibility relationships of the haplotypes to generate the diploid incompatibility matrix.
generate.diplo.compatibility <- function(haplo.compatibility, genos){
  # We have a matrix of all the diploid genotypes.
  # These will be the rownames.
  # Create a matrix that shows whether the pollen haplotype is compatible with the left of the two RNases the dam has.
  compatible_with_left <- matrix(F, nrow = nrow(genos), ncol = ncol(haplo.compatibility))
  rownames(compatible_with_left) <- genos[,1]
  colnames(compatible_with_left) <- colnames(haplo.compatibility)
  compatible_with_left[,] <- t(haplo.compatibility)[genos[,1],]
  compatible_with_right <- matrix(F, nrow = nrow(genos), ncol = ncol(haplo.compatibility))
  rownames(compatible_with_right) <- genos[,2]
  colnames(compatible_with_right) <- colnames(haplo.compatibility)
  compatible_with_right[,] <- t(haplo.compatibility)[genos[,2],]
  # To be compatible with a genotype, the pollen haplotype has to be compatible with both dam haplotypes.
  out <- compatible_with_left & compatible_with_right
  rownames(out) <- paste(genos[,1], genos[,2], sep = '_')
  colnames(out) <- colnames(haplo.compatibility)
  # Sort the output.
  out <- out[sort(rownames(out)),]
  out <- out[,sort(colnames(out))]
  return(out)
}

# Get the expected genotype frequencies after a single generation of mating.
# It's possible that I accidentally included pollen limitation in this simulation.
# One way to test would be to give a very common haplotype a very rare pollen source.
# It can't be zero, because zero really does affect fertilization.
# But it shouldn't matter if it's rare or common.
expected.one.gen <- function(genos, geno.dist, pollen.production.matrix, diplo.compatibility){
  # Create the pollen pool.
  pollen_pool <- produce.pollen(geno.dist, pollen.production.matrix)
  # Fill a matrix with the frequency of each kind of compatible pollen.
  raw_crosses <- matrix(pollen_pool, nrow = nrow(diplo.compatibility), ncol = ncol(diplo.compatibility), byrow = T) *
                 diplo.compatibility
  # Normalize the the rows.
  normalized_crosses <- raw_crosses / (rowSums(raw_crosses))
  # Crosses that never happen never happen.
  normalized_crosses[is.na(normalized_crosses)] <- 0
  # Weight each row by the frequency of the dam involved.
  true_crosses <- normalized_crosses * geno.dist
  # Replicate the crosses matrix but change the names.
  left_ovule <- true_crosses * 0.5
  rownames(left_ovule) <- genos[,1]
  right_ovule <- true_crosses * 0.5
  rownames(right_ovule) <- genos[,2]
  # CAN'T FIGURE IT OUT, DOING LOOPS.
  # The data frame "collected" contains the ovule-by-pollen crosses involved in every diploid-by-pollen pollination.
  # So A_C1 x L1 produces one instance of A x L1 and one of C1 x L1.
  # A_L1 X L1 makes another A x L1 and a L1 x L1, neither of which works.
  # The third column is the frequency with which that ovule-by-pollen fertilization actually occurs.
  # Crosses between nonexistent plants or pollen don't show up, and neither do illegitimate pollinations.
  collected <- data.frame(c(rep(rownames(left_ovule), times = ncol(left_ovule)), rep(rownames(right_ovule), times = ncol(right_ovule))),
                          c(rep(colnames(left_ovule), each = nrow(left_ovule)), rep(colnames(right_ovule), each = nrow(right_ovule))),
                          c(as.vector(left_ovule), as.vector(right_ovule))
  )
  out <- geno.dist
  has_ever <- vector(length = nrow(collected))
  for(i in 1:length(out)){
    produces_it <- paste(collected[,1], collected[,2], sep = '_') == names(out[i]) |
                   paste(collected[,2], collected[,1], sep = '_') == names(out[i])
    has_ever <- has_ever | produces_it
    out[i] <- sum(collected[produces_it,3])
  }
  # Crosses that DON'T show up in has_ever should all produce genotypes that are not in the genotype vector because they are impossible.
  # Crosses that are merely impossible because their alleles don't exist SHOULD show up in has_ever.
  return(out)
}

# Simulate.
simulate <- function(genos, geno.dist, pollen.production.matrix, diplo.compatibility, generations, N){
  out <- matrix(nrow = generations+1, ncol = length(geno.dist))
  colnames(out) <- names(geno.dist)
  out[1,] <- geno.dist
  for(i in 2:(generations+1)){
    current_dist <- expected.one.gen(genos, out[i-1,], pollen.production.matrix, diplo.compatibility)
    current_sample <- sample(names(current_dist), size = N, replace = T, prob = current_dist)
    for(j in colnames(out)){
      out[i,j] <- mean(current_sample == j)
    }
  }
  return(out)
}

# Iterate.
# Identical to simulate, except you take the expected frequencies rather than sampling from that distribution.
iterate <- function(genos, geno.dist, pollen.production.matrix, diplo.compatibility, generations){
  out <- matrix(nrow = generations+1, ncol = length(geno.dist))
  colnames(out) <- names(geno.dist)
  out[1,] <- geno.dist
  for(i in 2:(generations+1)){
    out[i,] <- expected.one.gen(genos, out[i-1,], pollen.production.matrix, diplo.compatibility)
  }
  return(out)
}

# Calculate the diploid genotype frequencies to haplotype frequencies
hap.from.dip <- function(geno.dist){
  # First, split out the names.
  genos <- matrix(nrow = length(geno.dist), ncol = 2)
  for(i in 1:length(geno.dist)){
    genos[i,] <- strsplit(names(geno.dist)[i], split = '_')[[1]]
  }
  out <- vector(length = length(unique(as.vector(genos))))
  names(out) <- unique(as.vector(genos))
  for(i in 1:length(out)){
    out[i] <- 0.5 * sum(geno.dist[genos[,1] == names(out[i]) | genos[,2] == names(out[i])])
  }
  return(out)
}

# Get the selection coefficient.
get.sel <- function(geno.dist, pollen.production.matrix, diplo.compatibility, what.haplo){
  # The probability that a given plant is fertilized by pollen of a given type is the probability that,
  # out of all compatible pollen, the pollen grain was of that type.
  # First, get the pollen vector.
  pollen_pool <- produce.pollen(geno.dist, pollen.production.matrix)
  # Now get the frequency of ALL compatible pollen for each genotype.
  compatible_supply <- colSums(t(diplo.compatibility) * pollen_pool)
  # Divide each genotype frequency by the fraction of pollen that is compatible with it.
  # This is the probability that a given compatible pollen grain will fertilize a plant of this genotype.
  will_fertilize <- (matrix(geno.dist, nrow = length(geno.dist), ncol = length(pollen_pool)) *
                            diplo.compatibility) /
                            compatible_supply
  # Now that we know the fraction of ovules of each plant that will be fertilized by each pollen type, we need to weight it by
  # the frequency of that genotype.
  # fertilization <- fraction_of_compatible * geno.dist
  siring <- colSums(will_fertilize)
  weighted_siring <- siring * pollen_pool
  # Now, get the mean siring success of all haplotypes.
  mean_siring <- sum(weighted_siring)
  # The mean fitness is (mean sire success)/2 + 1/2 because ovules always succeed.
  mean_fit <- (mean_siring)/2 + 1/2
  # Now calculate the fitness of the haplotype in question.
  haplo_siring <- siring[what.haplo]
  haplo_fit <- (haplo_siring)/2 + 1/2
  # Calculate Haldane's selection coefficient.
  sel <- (haplo_fit - mean_fit)/mean_fit
  return(sel)
}

# Infer the selection coefficient by iterating forward a generation and seeing what happens to the allele frequency.
# Generate a single mutant (1/2N frequency)
realized.sel <- function(genos, geno.dist, pollen.production.matrix, diplo.compatibility, N){
  mutated <- geno.dist
  mutated['L1_U3'] <- mutated['L1_U3'] - 1/(N)
  mutated['L1_L3'] <- mutated['L1_L3'] + 1/(N)
  subsequent_generation <- iterate(genos, mutated, pollen.production.matrix, diplo.compatibility, generations = 1)[2,]
  subsequent_al_freq <- hap.from.dip(subsequent_generation)['L3']
  current_al_freq <- hap.from.dip(mutated)['L3']
  # s = (p' - p)/p
  s <- (subsequent_al_freq - current_al_freq)/current_al_freq
  return(s)
}
```
