## Supplementary material for "Diversification or collapse of self-incompatibility haplotypes as a rescue process": All scripts: sc.R

```
# SI-SC balance model based on evaluating transmission through ovules and transmission through pollen separately

#############
### MODEL ###
#############

### Parameters ###
# SC haplotypes (C) are generated from SI haplotypes (I) at rate m every generation
# CC individuals self at rate s, IC individuals self at rate s/2, and II individuals never self
# There exist n distinct SI haplotypes

### Pollen success ###
# I pollen is rejected at rate (n-2)/n, whereas C pollen is never rejected

### Ovule success ###
# Selfed ovules are inviable

### Assumptions ###
# C haplotypes are rare
# Mutation occurs in zygotes
# Pollen success is determined as if all individuals are heterozygous at the RNase locus
# The frequencies of all RNase alleles are equal

#################
### FUNCTIONS ###
#################

# The function delP evaluates the change in the frequency of C haplotypes
# This is a cubic function
# Its biologically relevant solutions occur in the range [0,1] in the real numbers
# Exploration with Mathematica suggests there are usually either zero or two solutions in that range
# Furthermore, it appears that, usually, the smaller solution is stable, and the larger solution is unstable
# When the larger solution violates the assumption that C is rare, it is not clearly meaningful
delP <- function(p, # The current frequency of C haplotypes
                 s, # The selfing rate of CC individuals
                 m, # The mutation rate from I to C
                 n){# The number of RNase alleles
  return(p*((1-s/2)/(2 - 2*s*p) + (1)/(2 - 4/n - 4*p/n) - 1 - m) + m)
}

##################
### PARAMETERS ###
##################

# Selfing rate
s <- 0.5

# Mutation rate
m <- 0.0001

# Number of RNase alleles
n <- 40:10

################
### PLOTTING ###
################
x <- seq(from = 0, to = 1, by = 0.0001)
y <- rep(NA, length(x))
for(i in 1:length(y)){
  y[i] <- delP(x[i], s, m, n[1])
}

plot(x, y, type = 'l')
abline(h=0)
abline(v=0)
for(i in 1:length(n)){
  for(j in 1:length(y)){
    y[i] <- delP(x[i], s, m, n[j])
  }
  points(y, type = 'l')
}
##################
### Evaluation ###
##################

# # For each parameter set, scan through the range [0,1] in increments of the smallest mutation rate
# for(i in s){ # Selfing rate
#   for(j in m){ # Mutation rate
#     for(k in n){ # RNase number
#       for(l in seq(from = 0, to = 1, by = m[1])){ # Current frequency
#         delVal[h] <- delP(p=l,s=i,m=j,n=k)
#         h <- h+1
#       }
#     }
#   }
# }
# 
# # Identify sign changes
# # Multiply each value by the next value
# signs <- delVal[1:(length(delVal)-1)] * delVal[2:length(delVal)]
# # Negative values imply sign changes
# changes <- signs<0
# # Count changes
# numChanges <- sum(changes)
```
