## Supplementary material for "Diversification or collapse of self-incompatibility haplotypes as a rescue process": All scripts: simulate_survival.sh

```
#!/bin/bash
for GEN in 10 20 30 40 50 60 70 80 90 100 110 120 130 140 150 160 170 180 190 200 210 220 230 240 250 260 270 280 290 300 310 320 330 340 350 360 370 380 390 400 410 420 430 440 450 460 470 480 490 500 510 520 530 540 550 560 570 580 590 600 610 620 630 640 650 660 670 680 690 700 710 720 730 740 750 760 770 780 790 800 810 820 830 840 850 860 870 880 890 900 910 920 930 940 950 960 970 980 990 1000; do
		Rscript stochastic_survival.R $GEN
done
```
