## Supplementary material for "Diversification or collapse of self-incompatibility haplotypes as a rescue process": All scripts: stochastic_survival.R

```
# The purpose of this script is to check the analytic approximation for survival probability of a new gene convertant.

source('rnase_functions.R')

expected_trajectory <- as.matrix(read.csv('../data/L=2_U=2_split_traj.csv', as.is = T, row.names = 1))

genos <- matrix(nrow = ncol(expected_trajectory), ncol = 2)
# Recover parameters from the trajectory.
for(i in 1:ncol(expected_trajectory)){
  genos[i,] <- strsplit(colnames(expected_trajectory)[i], split = '_')[[1]]
}
pollen_prod <- generate.pollen.production(genos)
hap_comp <- generate.haplo.compatibility(genos)
dip_comp <- generate.diplo.compatibility(hap_comp, genos)
# N is tricker because it cannot be recovered from the trajectory.
# I should probably put it in there somehow.
# For now, I'm hardcoding it at a constant N = 10000 for all scripts.
N <- 10000

args <- commandArgs(trailingOnly = TRUE)

start_gen <- eval(parse(text = args[1]))

sims <- 1

survival_proof <- 10

survived <- vector(length = sims)
for(i in 1:sims){
  sim_fodder <- expected_trajectory[start_gen,]
  sim_fodder['L1_U3'] <- sim_fodder['L1_U3'] - 1/(N)
  sim_fodder['L1_L3'] <- sim_fodder['L1_L3'] + 1/(N)
  # sim_fodder <- sim_fodder/sum(sim_fodder)
  result <- simulate(genos, sim_fodder, pollen_prod, dip_comp, generations = survival_proof, N = N)
  final_al <- hap.from.dip(result[nrow(result),])
  survived[i] <- final_al['L3'] > 0
}

out <- t(as.matrix(c(start_gen, mean(survived))))

write.table(out, file = '../data/sim_survival.csv', append = T, sep = ',', col.names = F, row.names = F)
```
