## Supplementary material for "Diversification or collapse of self-incompatibility haplotypes as a rescue process": All scripts: survival_sim.R

```
# This script simulates the probability of survival of a new gene convertant.
# The simulation begins with initial conditions drawn from a single point in time during the deterministic rescue process.
# From there, it simulates the fate of all diploid genotype frequencies.
# It stops when a) the convertant is lost or b) the convertant reaches its equilibrium frequency.
# It repeats this and outputs the fraction of simulations resulting in survival, which can be compared to the analytic approximation.

# Load the function definitions.

######################
# Initial conditions #
######################

# Generating initial conditions by running rescue_prob.R
# This will both generate a trajectory of diploid genotype frequencies and calculate survival probabilities throughout the process.
# We can then run the simulation from some arbitrary point and compare to the analytic approximation.

# Define parameters
# Some parameters, such as the number of doomed and lucky haplotypes, will already be defined after running rescue_prob.R.

##############
# Simulation #
##############

# The simulation will track the diploid genotype frequencies through time.
# Each generation, it will create a vector of the probability distribution of sampling a given offspring genotype.
# E.g., there is a 0.01 probability of sampling LS in the next generation given the parental genotype frequencies.
# Once this vector is done, the simulation will sample from this distribution once per individual in the population.
# The frequencies will then be calculated from the raw numbers, and the process will be repeated.
# The population size affects the number of draws, so drift is automatically incorporated.

# To do the simulation, we need the probability of each genotype as a function of the previous genotypes.
# This is basically already done in rnase_functions.R.
# In fact, we could probably use the deterministic one.generation function and just sample using the distribution it produces.

# The simulation will therefore use one.generation to create the expected frequencies (i.e., probabilities) of all genotypes,
# sample from that distribution N times,
# and divide by the sum to get the frequencies.

# The problem with this idea is that it treats the classes with multiple haplotypes (L, C, and U) as single entities.
# However, the fertilization probabilities change as the frequencies of haplotypes within a class diverge.
# E.g., if most class-L haplotypes are copies of the same haplotype, then LxL crosses will decrease.
# Furthermore, there are correspondences between members of different classes: for every C, there's an L.
# Simply creating vectors of the frequencies of these haplotypes wouldn't work because it would ignore covariance between alleles in
# a diploid
# One option would be to assume that the frequencies within a class behave deterministically, but that is unsatisfying.
# The other option is to track each haplotype separately, but this allows an arbitrary number of haplotypes and genotypes.
# One fortunate thing is that the only differences between haplotypes of the same class is their RNase specificity.
# However, the fertilization probabilities should still be affected by the covariance between alleles.

# The other options bite the bullet to various degrees.
# The first is just to do an individual-based simulation.
# Individual-based simulations are the worst, so I'm not doing that.
# The other option is to create a gentoype vector of arbitrary size depending on the number of haplotypes.
# Since the behavior of all members of a haplotype class is the same except with respect to itself, you wouldn't have to write new
# mating rules for each haplotype.
# I think I will go with this option.

# Unfortunately, the genotype-frequency recursions are not linear functions of genotype frequencies.
# Therefore, I can't populate a transition matrix and be done with it.
# One possibility is to do the same thing as the function one.generation, except supply a vector of frequencies instead of (L - 1 / L).
# Do this once for each multi-haplotype class.
# Furthermore, instead of having a compatible.with.XX variable for each genotype, create a compatible.with matrix with a row for each
# diploid genotype and a column for each pollen haplotype.
# If you right-multiply the compatible.with matrix by a matrix in which each column is just copies of one of the genotype frequencies,
# then right-multiply that by the vector of pollen haplotype frequencies, you should get the new vector of genotype frequencies.
# I don't think there's an easy way to describe the long-term behavior of a system like that, but you can still iterate matrix math.
# Once you've got the expected frequency distribution, sample from that to make the next generation.

# The hardest part will be calculating the compatibility matrix.
# I know the algorithm, but I don't know how to implement it in R.
# The part that bugs me is that there can be an arbitrary number of haplotypes.
# Then, you have to keep the genotypes in a particular order, and the size of each fertilization category (e.g., LLxL) is variable.
# I guess you can just assign the compatibilities as if there were no others of your kind, then go back and find the ones of your kind
# and set them to zero.
# As a matter of fact, if we're putting the pollen haplotype frequencies in a different vector, the compatibility matrix only has ones
# and zeros.
# Since the columns are just haplotypes rather than diploid genotypes, you can designate whole stretches as behaving identically.
# You could probably co-opt the hard-coded version by repeating certain compatibility classes a number of times equal to the number
# of haplotypes in each class, then go back and get rid of the ones where like pollen is landing.

# If we were to do an individual-based simulation, we could at least enumerate all the possible genotypes at the outset.
# But there might be a fairly large number of combinations.

# Here's how I would write an individual-based simulation.
# Each individual would consist of a vector of length two.
# Each element would be a string representing a haplotype.
# Each haplotype would be either one or two characters.
# E.g., L1, C2, U1, A, S.
# Another possibility would be to just give them numerical names, and store their phenotypic effects in a separate matrix.
# That would give each one a unique identifier.
# Somehow, I need to know whether they are compatible with each other.
# Besides storing their effects in a compatibility matrix, the other option is to base their effects on their names.
# This would only work if they have strings as names.
# The obvious thing to do would be not to let things that share at least one character to mate, but then U needs a different
# numerical series from L and C.
# It's not obvious to me whether it's better to hardcode it or use a compatibility matrix.
# I guess logically they use the same number of operations if you go through all the possible crosses, but maybe keeping the compatibility
# matrix in memory or consulting it is burdensome.
```
