## Supplementary material for "Diversification or collapse of self-incompatibility haplotypes as a rescue process": All scripts: transition_probs.R

```
# This script is like rescue_prob.R except that it calculates the probability distribution of the final number of alleles
# instead of merely the probability that it is n+1 or less than n+1.

# Choose a population size.

N <- 400000

# Load the functions.

source('rnase_functions.R')

# Initialize

my_gens <- 5000
doomed_max <- 40 # The maximum number of doomed alleles tracked by the transition matrix.
# The probability above this upper boundary will be collapsed into a point at the boundary.
# This is incorrect because all the numbers above the upper bound have different transition probabilities, but I will ignore them.
# This might not matter if the maximum is high because these high number universes contribute almost no probability anyway.
L <- 2 # L is held constant as a parameter even though the number of lucky haplotypes is actually subject to chance.
luck_frac <-0.01 # The fraction of the lucky RNase haplotypes that also have the new SLF.
spite_frac <- 0.01 # The fraction of the ancestral RNase that is replaced with the novel RNase.
conversion_rate <- 10 ^ -6 # The per-heterozygote probability of gene conversion.

# Fill out a transition matrix.
# It describes the probabilities of transition between any two
trans_mat <- matrix(nrow = doomed_max+2, ncol = doomed_max+2)
rownames(trans_mat) <- L:(L+doomed_max+1)
colnames(trans_mat) <- L:(L+doomed_max+1)
# Figure out the probability of transitioning from one number to any other number.
for(U in 1:(doomed_max+1)){
  initialize(L,
             U,
             luck_frac,
             spite_frac)
  
  # Create trajectories of diploid genotype frequencies.
  # Plot trajectories of haplotype frequencies.
  
  traj <- track.diplos(population, my_gens)
  
  # To calculate the supply of gene convertants of a particular type, we need a vector of the frequencies of diploid genotypes
  # that can produce that convertant.
  
  my_convertible_traj <- traj[,'LU']/U
  # Track the survival probability of A separately since it has a different starting frequency.
  my_convertible_traj_A <- traj[,'LA']
  
  my_supply <- generate.supply(my_convertible_traj, N, conversion_rate)
  my_survive <- determine.survival.prob(traj,N)
  my_supply_A <- generate.supply(my_convertible_traj_A, N, conversion_rate)
  my_survive_A <- determine.survival.prob(traj,N, doomed.kind='A')
  
  # You can back-calculate the selection coefficients from the survival = 2s approximation and put the resulting s into series.solver
  my_series_survive <- sapply(my_survive/2, series.solver, guesses = seq(from = 0, to = 1, by = 0.01), terms = 100)
  my_series_rescue <- rescue.prob(my_supply, my_series_survive)
  my_series_survive_A <- sapply(my_survive_A/2, series.solver, guesses = seq(from = 0, to = 1, by = 0.01), terms = 100)
  my_series_rescue_A <- rescue.prob(my_supply_A, my_series_survive_A)
    
  # Instead of getting a single probability that all haplotypes are rescued, calculate the probability distribution of the final number
  # of haplotypes assuming that only U haplotypes are doomed.
  survivors <- dbinom(0:doomed_max, U, my_series_rescue)
  survivors_A <- dbinom(0:1, 1, my_series_rescue_A)
  # If the maximum number has been reached, the probability that all are rescued is collapsed into the probability that all but one are.
  if(U == doomed_max+1){
    survivors[length(survivors)-1] <- survivors[length(survivors)]
    survivors[length(survivors)] <- 0
  }
  # There are essentially two distributions:
  # the number of Us conditional on A surviving and the number of Us conditional on A dying
  Us_and_A <- survivors * survivors_A[2]
  Us_no_A <- survivors * survivors_A[1]
  all_survivors <- c(Us_no_A[1],
                     Us_no_A[2:length(Us_no_A)] + Us_and_A[1:(length(Us_and_A)-1)],
                     Us_and_A[length(Us_and_A)]
  )
  trans_mat[U+1,] <- all_survivors
}

# Write the transition matrix to a file
outname <- paste('../data/transition_L=', L, '_R=', N*conversion_rate, '.csv', sep = '')
write.csv(trans_mat, outname)

# Plot the entire transition matrix as a heatmap
library(viridis)
library(fields)
mypal <- viridis(n = length(unique(trans_mat)))
# pdf('../figures/tmatrix.pdf')
image.plot(x = L:(L+doomed_max+1),
           y = L:(L+doomed_max+1),
           z = trans_mat,
           col = mypal,
           ylab = 'Haplotypes after transition',
           xlab = 'Haplotypes before transition',
           legend.lab = 'Transition probability'
           )
# dev.off()
  
# Can use this plot to visualize the probability distribution of transition probabilities from a certain starting point.
# plot(x = 0:(length(survivors)-1),
#      y = trans_mat[50,],
#      type = 'h',
#      xlab = 'Rescued haplotypes',
#      ylab = 'Probability')
```
