## Supplementary material for "Diversification or collapse of self-incompatibility haplotypes as a rescue process": All scripts: view_trajectories.R

```
# The purpose of this script is to create graphs of the haplotype trajectories.
source('rnase_functions.R')

my_gens <- 1000 # The number of generations you want to track.
L <- 2 # The number of haplotypes that have a variant with the new key.
U <- 2 # The number of haplotypes that have no variant with the new key.
luck_frac <- 0.01 # The fraction of each of the L haplotypes that actually has the new key.
spite_frac <- 0.01 # The fraction of the lock-mutant's ancestor that actually mutated to the lock-mutant.

initialize(L,
           U,
           luck_frac,
           spite_frac)

# Create trajectories of diploid genotype frequencies.
# Plot trajectories of haplotype frequencies.

traj <- track.diplos(population, my_gens, monster = F)

# Put underscores

# Turn the genotype trajectory into a haplotype trajectory.
al_traj <- matrix(nrow = nrow(traj), ncol = 5)
colnames(al_traj) <- c('L','C','U','A','S')
for(i in 1: nrow(al_traj)){
  al_traj[i,] <- diplos.to.haplos(traj[i,])
}

# library(RColorBrewer)
pdf('../figures/equilibration.pdf', width = 11, height = 8.5)
par(mar = c(8,9,4,2), mgp = c(6, 2, 0), cex.axis = 3, cex.lab = 4)
# my_pal <- brewer.pal(5, 'Dark2')
plot(al_traj[,'L'], type = 'l', ylim = c(0,0.7), xlim = c(0,my_gens),
     xlab = 'Generation',
     ylab = 'Frequency',
     # col = my_pal[1],
     lwd = 5, lty = 1
)
lines(al_traj[,'C'], #col = my_pal[2],
      lwd = 5, lty = 2)
lines(al_traj[,'U'], #col = my_pal[3],
      lwd = 5, lty = 3)
lines(al_traj[,'A'], #col = my_pal[4],
      lwd = 5, lty = 4)
lines(al_traj[,'S'], #col = my_pal[5],
      lwd = 5, lty = 5)
# legend(x= 'topright',
#        legend = c('L',
#                   'C',
#                   'U',
#                   'A',
#                   'S'
#                   ),
#        fill = my_pal,
#        ncol = 2,
#        cex = 2,
#        bg = 'white')
text(x = c(250), y = al_traj[250,c('L','C','U','A','S')]+0.03, labels = c('L', 'C', 'U', 'A','M'), cex = 2)
dev.off()

# Repeat for the invasion of the monster haplotype, starting where you left off.
# Strictly speaking, you're magically replacing every S with the monster version of S.
# In reality, there would be a new monster version of S that would compete with S.
# However, since S is expected to stay at low frequency and also be eliminated when monster-S shows up,
# it isn't that unrealistic to do magical replacement.

# Create trajectories of diploid genotype frequencies.
# Plot trajectories of haplotype frequencies.

monster_population <- traj[nrow(traj),]

monster_traj <- track.diplos(monster_population, my_gens, monster = T)

# Put underscores

# Turn the genotype trajectory into a haplotype trajectory.
monster_al_traj <- matrix(nrow = nrow(monster_traj), ncol = 5)
colnames(monster_al_traj) <- c('L','C','U','A','S')
for(i in 1: nrow(al_traj)){
  monster_al_traj[i,] <- diplos.to.haplos(monster_traj[i,])
}

# Plot
pdf('../figures/dwindle.pdf', width = 11, height = 8.5)
par(mar = c(8,9,4,2), mgp = c(6, 2, 0), cex.axis = 3, cex.lab = 4)
plot(monster_al_traj[,'L'], type = 'l', ylim = c(0,0.7), xlim = c(0,my_gens/10),
     xlab = 'Generation',
     ylab = 'Frequency',
     # col = my_pal[1],
     lwd = 5, lty = 1
)
lines(monster_al_traj[,'C'], #col = my_pal[2],
      lwd = 5, lty = 2)
lines(monster_al_traj[,'U'], #col = my_pal[3],
      lwd = 5, lty = 3)
lines(monster_al_traj[,'A'], #col = my_pal[4],
      lwd = 5, lty = 4)
lines(monster_al_traj[,'S'], #col = my_pal[5],
      lwd = 5, lty = 5)
# legend(x= 'topright',
#        legend = c('L',
#                   'C',
#                   'U',
#                   'A',
#                   'S*'
#        ),
#        fill = my_pal,
#        ncol = 2,
#        cex = 2,
#        bg = 'white')
text(x = c(30), y = monster_al_traj[30,c('L','C','U','A','S')]+0.03, labels = c('L', 'C', 'U', 'A', 'M'), cex = 2)
dev.off()
```
