## Supplementary figures and images for "Diversification or collapse of self-incompatibility haplotypes as a rescue process"

### test.pdf

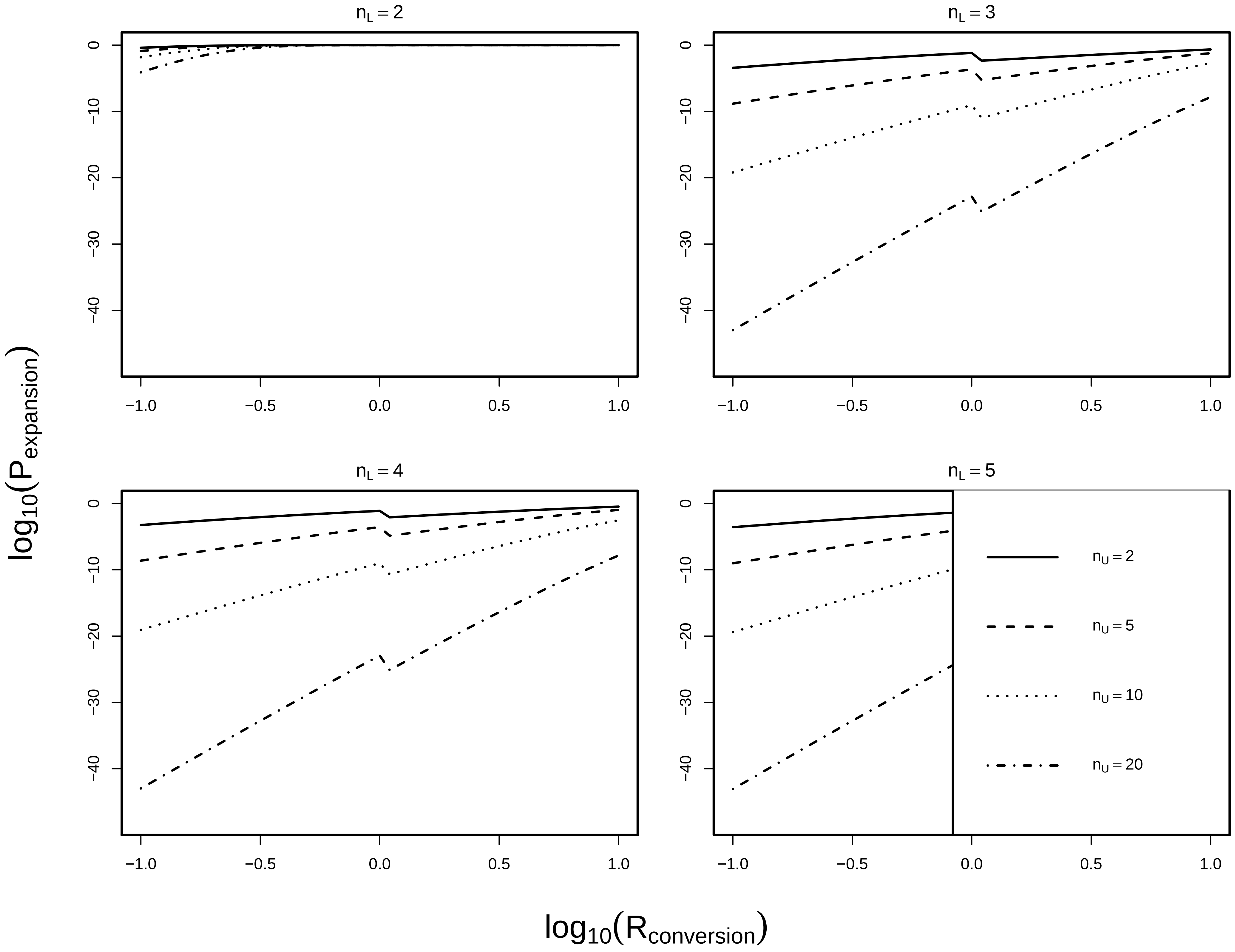
